# UniFORM: Towards Universal ImmunoFluorescence Normalization for Multiplex Tissue Imaging

**DOI:** 10.1101/2024.12.06.626879

**Authors:** Kunlun Wang, Kaoutar Ait-Ahmad, Sam Kupp, Zachary Sims, Eric Cramer, Zeynep Sayar, Jessica Yu, Melissa H. Wong, Gordon B. Mills, S. Ece Eksi, Young Hwan Chang

**Author notes:** Co-corresponding author: S. Ece Eksi, PhD, Assistant Professor, Cancer Early Detection Advanced Research (CEDAR), Knight Cancer Institute, Oregon Health and Science University, Portland, Oregon, Young Hwan Chang, PhD, Associate Professor, Department of Biomedical Engineering and Computational Biology Program, Knight Cancer Institute, Oregon Health and Science University, Portland, Oregon.

## Abstract

Multiplexed tissue imaging (MTI) technologies enable high-dimensional spatial analysis of tumor microenvironments but face challenges with technical variability in staining intensities. Existing normalization methods, including Z-score, ComBat, and MxNorm, often fail to account for the heterogeneous, right-skewed expression patterns of MTI data, compromising signal alignment and downstream analyses. We present UniFORM, a non-parametric, Python-based pipeline that uses an automated rigid landmark functional data registration approach for normalizing both feature- and pixel-level MTI data. Designed specifically for the distributional characteristics of MTI datasets, UniFORM operates without prior distributional assumptions and performs robustly regardless of distribution modality, including both unimodal and bimodal patterns. It removes technical variation by aligning the biologically invariant component of the signal, typically the negative (non-expressing) population, while preserving biologically meaningful variation in the positive population, thereby maintaining tissue-specific expression patterns essential for downstream analysis. Benchmarking across three distinct MTI platform datasets demonstrates that UniFORM outperforms existing methods in mitigating batch effects while maintaining biological signal fidelity. This is evidenced by improved marker distribution alignment and positive population preservation, enhanced kBET and Silhouette scores, and improved downstream analyses such as UMAP visualizations and Leiden clustering. UniFORM also introduces a novel guided fine-tuning option for complex and heterogeneous datasets. Although optimized for fluorescence-based platforms, UniFORM provides a scalable and robust solution for MTI data normalization, enabling accurate and biologically meaningful interpretations.

## Introduction

Multiplexed tissue imaging (MTI) technologies, including cyclic immunofluorescence (CyCIF)^1^, multiplexed immunohistochemistry (IHC)^2–4^, and CODEX^5^, have significantly advanced our ability to visualize and analyze the complexities of the tumor microenvironment at both single-cell and subcellular levels^6^. These techniques enable the simultaneous detection of multiple markers within a single tissue section, offering a detailed view of cellular interactions and functional states^6^. However, a major challenge remains in integrating data from various samples and batches due to technical variations in staining intensities^7^, which can arise from differences in tissue fixation, antibody concentrations, and imaging conditions^7–9^. These variations necessitate robust normalization procedures to ensure accurate biological interpretations.

MTI datasets typically exist in two formats: 1) feature-level data, which usually contains mean intensity values across markers for individual cells, and 2) pixel-level data, which comprises the raw registered multi-channel images^10^. Most existing intensity normalization methods are tailored to feature-level data, where intensity measurements of individual markers are adjusted across different samples to mitigate technical variability. Commonly used methods can be categorized into parametric and non-parametric approaches. Parametric methods assume the data follows a specific parametric distribution (e.g., Gaussian or normal distribution). Typical examples of this category include Z-score^11,12^ and ComBat^12,13^, where Z-score normalization standardizes data based on mean and standard deviation, and ComBat adjusts for batch effects using empirical Bayes methods. Non-parametric methods make no explicit assumptions about the data’s distribution. The MxNorm^8^ framework incorporates several such approaches, including mean division and B-spline registration. In mean division, intensity values are normalized by dividing by the mean intensity of each marker within each image. B-spline registration aligns intensity distributions across samples through functional data registration using B-spline basis functions.

Despite their utility, these methods often fall short when applied to MTI datasets. Z-score and ComBat normalization, both feature-level normalization methods, rely on the assumptions of normally or symmetrically distributed data and consistent cell composition across samples^13–15^. However, the right-skewed and heterogeneous expression patterns, driven by variations in cellular composition across tissues – typical characteristics of MTI data – violate these fundamental assumptions. Although MxNorm’s mean division and registration-based normalization algorithms do not assume normality or symmetry of the data distribution, they remain sensitive to distributional characteristics such as skewness and heterogeneity. Mean division relies on the arithmetic mean, which can be unstable in right-skewed, heterogenous marker distributions common in MTI data, leading to suboptimal normalization. B-spline registration assumes distributional differences are technical artifacts correctable by smooth warping, but it may underperform on highly skewed data and risk removing true biological variation. Consequently, these methods often face challenges in accurately aligning signals, preserving the original data distribution, and maintaining the integrity of the marker-positive population percentage. This makes them poorly suited for MTI feature normalization and can negatively impact downstream analyses, such as cell phenotyping and clustering.

Meanwhile, there is growing interest in pixel-level MTI applications such as image-to-image translation or virtual staining^17–19^, driven by recent advancements in deep learning. Notable examples include SHIFT^20^ and MIM-CyCIF^21^, both of which highlight the critical need for robust pixel-level normalization techniques to enhance performance. One of the few options is FLINO^9^, an R-based script that uses a grid-based quantile normalization workflow. While it may be effective in certain cases, FLINO fails to handle the right-skewness and heterogeneity of MTI data, resulting in overcorrection or undercorrection of marker intensities, misalignment of biologically meaningful signals, and potential amplification of batch effects or local intensity differences. Moreover, FLINO is computationally inefficient and lacks the scalability to handle large-scale MTI datasets.

A significant challenge in MTI applications is the lack of a universal normalization method that effectively handles both feature-level and pixel-level data. Existing approaches tend to prioritize one level over the other, limiting their ability to provide a comprehensive solution for the complex structure of MTI datasets. This underscores the need for a novel, unified approach that normalizes data at both levels, ensuring computational efficiency while accommodating the unique characteristics of MTI datasets to enable accurate and meaningful biological interpretations.

To address this need, we present UniFORM, a non-parametric and automated MTI intensity normalization pipeline developed in Python, designed for both feature-level and pixel-level datasets (**Figure 1a**). UniFORM uses our novel automated rigid landmark functional data registration approach that is not limited by any of the assumptions of existing methods and is tailored specifically to the complex nature of MTI data. A key innovation of UniFORM lies in its novel automatic normalization approach, which aligns marker intensity distributions while preserving both distribution shapes and the proportion of positive populations across samples— without requiring prior knowledge. Although this automated workflow performs effectively in most cases, guided fine-tuning options are also available for scenarios involving markers with high noise and heterogeneous expression pattern samples, allowing for further refinement when necessary.

**Figure 1:**
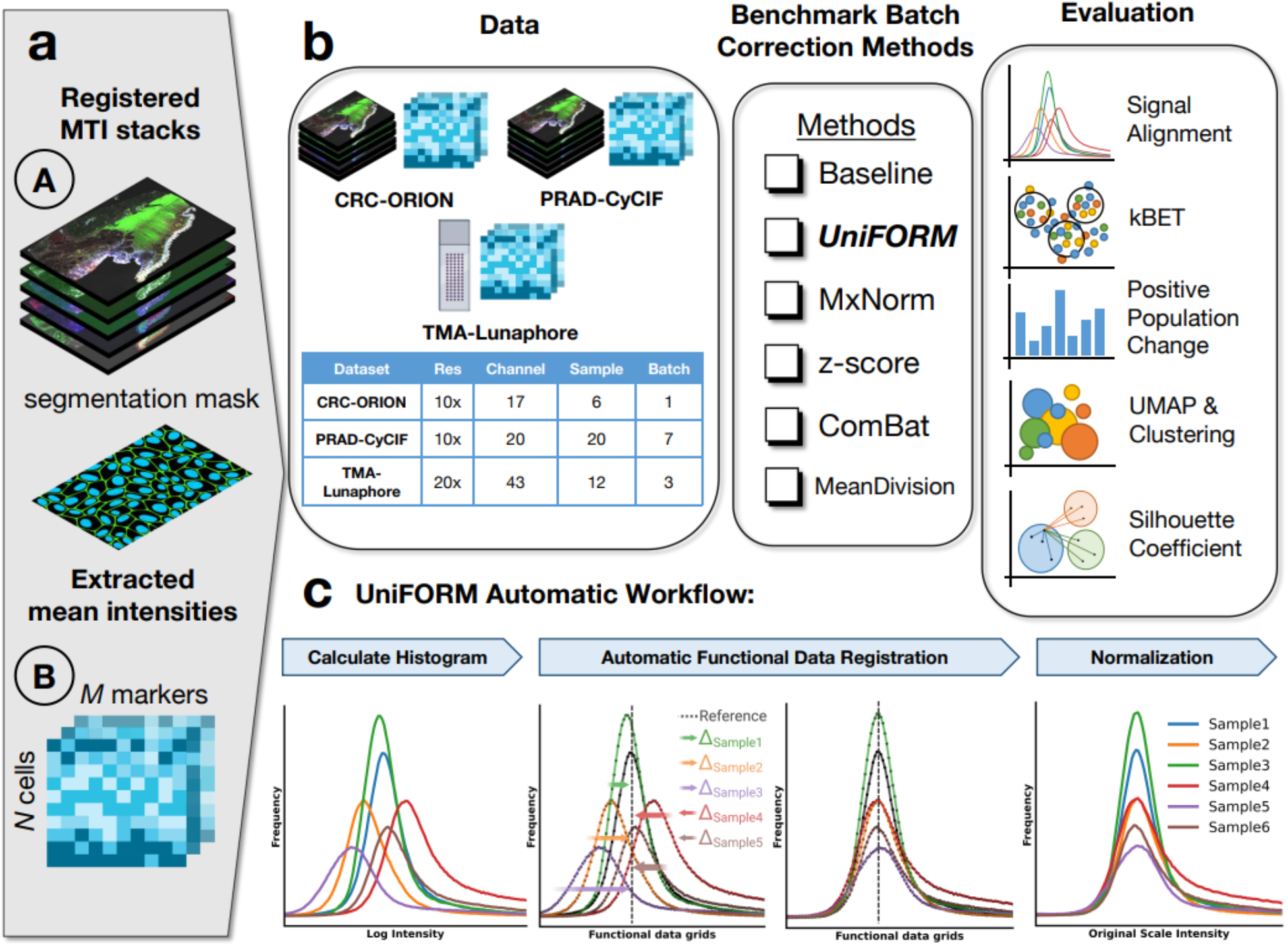
UniFORM offers universal automatic intensity normalization for both feature-level and pixel-level MTI data. **(a)** UniFORM accepts both pixel-level and feature-level data as input. Nucleus/cell segmentation masks are required for pixel-level normalization to exclude non-cell regions. **(b)** The study workflow includes datasets from two multiplex imaging platforms: CRC-ORION (17 channels, 6 patients, 1 batch), PRAD-CyCIF (20 channels, 20 patients, 7 batches), and TMA-Lunaphore (43 channels, 12 samples, 3 batches). UniFORM was benchmarked against commonly used normalization methods (MxNorm-registration, Z-score, ComBat, Mean Division) and a baseline. The performance of each method was evaluated through signal alignment, kBET analysis, positive population change, and UMAP-Leiden clustering. **(c)** Overview of the UniFORM automatic normalization workflow. The process begins with histogram calculation, followed by signal alignment using functional data registration, and concludes with normalization. A major novelty and highlight of UniFORM is its pixel-level normalization capability. It undergoes the same pipeline but uses cell/nuclei segmentation masks and a slightly different normalization step.

We demonstrate the feasibility of UniFORM as a universal normalization method for both feature-level and pixel-level data by benchmarking it against commonly used approaches, including MxNorm (registration option), Z-score, ComBat, and mean division across three cancer MTI datasets:

1. CRC-ORION (colorectal cancer whole slides, 6 samples from a single batch on the ORION platform by Rarecyte; **Table S1a**)
2. PRAD-CyCIF (prostate cancer whole slides, 20 samples across 7 batches on the CyCIF platform; **Table S1b**)
3. TMA-Lunaphore (a control slide composed of mixed tissue microarray cores from 12 tumor types, stained and imaged across three batches on the Lunaphore COMET platform; **Table S1c**).

Each dataset was chosen to assess specific normalization challenges: CRC-ORION for minor intra-batch technical variability, PRAD-CyCIF for correcting batch effects across heterogeneous samples, and TMA-Lunaphore for evaluating batch effect removal on identical tissue across batches. We evaluated each method’s performance by assessing its signal alignment, kBET, Silhouette score, positive population change, UMAP, and Leiden clustering^22^ before and after normalization (**Figure 1b**). Our results highlight the fundamental limitations of existing methods and show that UniFORM consistently outperforms them across all evaluation metrics, underscoring its effectiveness and suitability for MTI data.

## Results

Technical variations can significantly affect overall signal intensity in images, leading to inconsistencies across samples or batches. To address this, UniFORM utilizes an approach that aligns negative population peaks – cells that are not expressed for a given marker, which ideally represent background or baseline signals – to correct for these variations^7^. UniFORM does not require prior phenotyping or cell type annotations to identify this population, instead, they are inferred directly from the leftmost mode (lowest-intensity peak) of the marker’s histogram, which typically corresponds to non-expressing cells. In most cases, the negative population is observed to be the dominant component of the distribution. This alignment ensures that biologically invariant baseline signals remain consistent across samples, creating a stable foundation for comparing intensities across samples. In this study, we define the negative population as cells lacking expression of a given marker, and the positive population as those with detectable marker expression.

UniFORM’s automatic pipeline is based on two assumptions:

1. **Minimal Biological Variability in Negative Population:** For any given marker, the negative population across samples should exhibit minimal biological variability, serving primarily as an indicator of technical variation rather than biological differences.
2. **Consistent Distribution Patterns in Positive/Negative Population:** Marker intensity distributions are assumed to maintain a consistent pattern in positive and negative populations across samples.

The first assumption posits that the negative peak corresponds to cells lacking marker expression, making it a reliable baseline for background signals that account for technical variations. This assumption allows the negative peak to function as a stable baseline signal, typically reflecting background fluorescence and autofluorescence rather than specific marker expression. Because this negative peak reflects technical rather than biological variability, it serves as a stable baseline. Aligning these negative peaks across samples standardizes background signals, thereby correcting for technical variations and enabling comparable intensity measurements. The second assumption reinforces this approach by expecting consistency in the distribution of intensity patterns across samples, expecting a consistent pattern in marker intensity distributions across samples, where either the negative or positive peak consistently dominates. This consistency across samples allows us to align distributions automatically by maximizing the correlation between them, ensuring that intensity patterns remain comparable across samples. However, due to factors like tissue heterogeneity, marker specificity, or technical challenges, variations may occasionally arise. To address these situations, UniFORM includes a quick and guided fine-tuning option, allowing users to adjust as needed to maintain accuracy and consistency in their data.

### Feature-Level Normalization: Intensity Distribution Alignment and Batch Effect Removal

The feature-level MTI dataset, derived from cell or nuclei segmentation masks applied to pixel-level MTI data (see **Methods**), comprises mean cell intensity values along with metadata and spatial information. Our feature-level normalization approach (see **Methods**) is designed to be efficient and effective. Initially, we calculate the histogram distribution of the log-transformed cell intensities for each marker, allowing us to access insights into the central tendency, variability, skewness, modality, and inter-sample variations within the dataset. These distributions are then converted into functional data curves, which are aligned to an appropriate reference distribution using a maximum cross-correlation algorithm (**Figure 1c**). Finally, the raw feature data is normalized by applying the normalization factors estimated from this alignment process (see **Methods**). Once normalized with UniFORM, the resulting consistency in marker intensity distributions across samples enables the application of uniform gating thresholds across the entire dataset. This eliminates the need for manual, sample-specific gating—commonly required in unnormalized datasets—where technical variability often necessitates individually adjusted thresholds to account for non-biological differences.

Building on this approach, we performed the automatic normalization on the CRC-ORION, PRAD-CyCIF, and TMA-Lunaphore datasets (see **Data availability**). To evaluate the effectiveness of UniFORM in comparison to other methods, we first compared the alignment of the intensity distributions before and after normalization using MxNorm (registration option), Z-score, ComBat, and mean division against UniFORM. To further assess the effectiveness of batch correction, we used the kBET algorithm^23^ (**see Methods**), which constructs a k-nearest neighbor graph to evaluate the batch-label distribution within subsets of neighboring cells relative to the overall batch-label distribution. This approach allows us to quantify batch effect mitigation by measuring the degree of batch mixing within the data, providing a detailed comparison of UniFORM, MxNorm, ComBat, and z-score normalization methods specifically within the PRAD-CyCIF dataset. We further evaluated the preservation of biological composition using Silhouette^16^ coefficient analysis, focusing on three mutually exclusive cell types—tumor epithelial cells, immune cells, and non-immune stromal cells—to assess how well each normalization method maintains separation between distinct cell populations after batch correction.

**Figure 2a** shows examples of marker intensity distributions before and after normalization for the unimodal distribution of marker PD-L1 (CRC-ORION) and the bimodal distribution of marker ECAD (PRAD-CyCIF). For PD-L1, all methods seem to achieve a good alignment, however, only UniFORM preserves the distribution shape without alterations. For ECAD, UniFORM alone successfully aligns the peaks and retains the original distribution shape while other methods fail in both alignment and shape preservation. Additional marker alignments are provided in Supplementary **Figures S1-S2**. We performed intensity distribution alignment on the TMA-Lunaphore dataset as a separate analysis to further evaluate UniFORM’s performance. This dataset serves as a biologically controlled system, consisting of tissue microarray (TMA) cores imaged across different batches, specifically designed to assess batch correction effectiveness. Due to the dataset’s scale and its use as biological replicates, only UniFORM was applied for normalization in this context. The results (**Figure S3a–l**) reveal substantial batch effects even within the same TMA slide and demonstrate that UniFORM effectively corrects these technical variations while preserving biological signal.

**Figure 2:**
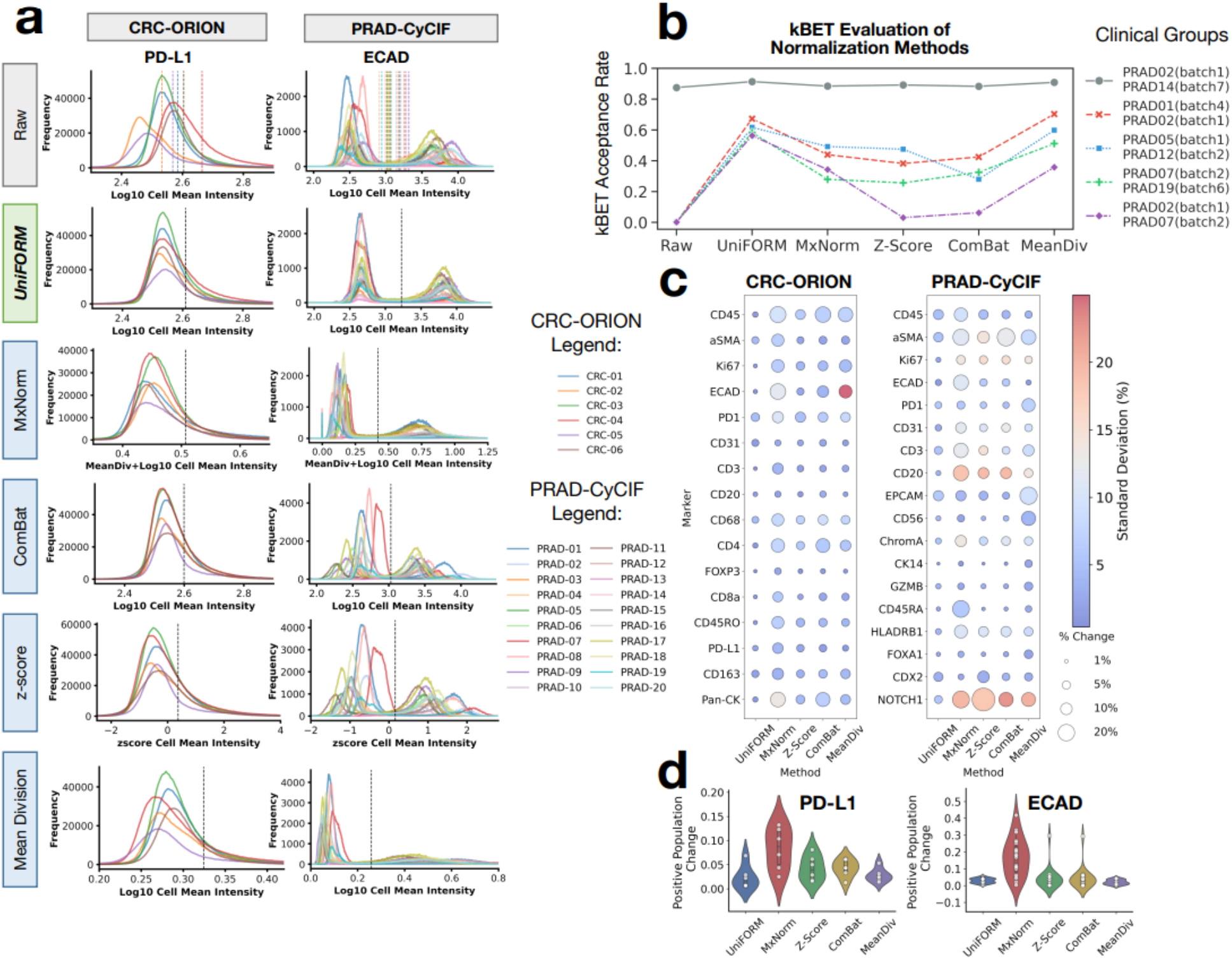
Feature-level normalization evaluation and comparison. **(a)** Evaluation of cell mean intensity signal alignment across different normalization methods of unimodal marker PD-L1 and bimodal marker ECAD, with local positive population thresholds (colorful lines) and global threshold (black line) annotated. **(b)** Evaluation of batch effect correction with kBET with five patient groups, each composed of patient samples with similar cell composition but of different batches, measured by kBET acceptance rate (higher rate indicates better batch correction). **(c)** Evaluation of positive population change post-normalization across different normalization methods. The size of the bubbles indicates the magnitude of the absolute positive population mean percentage change, and the color of the heatmap indicates the standard deviation of the mean percentage change. **(d)** Violin plots show the spread and central tendency of the positive percentage change for PD-L1 and ECAD.

More broadly, the distribution alignment results across all three MTI datasets—CRC-ORION, PRAD-CyCIF, and TMA-Lunaphore—highlight UniFORM’s robustness across diverse platforms, each presenting unique normalization challenges. These results confirm UniFORM’s ability to preserve distribution shape and achieve consistent signal alignment for both intra- and inter-batch normalization. Notably, UniFORM achieves near-perfect alignment of negative population peaks across all markers (**Figures S1–S3**) while maintaining the integrity of the original distribution shape. In contrast, existing methods distort signal distributions and fail to align intensities accurately (**Figures S1, S2**), which compromises cell phenotyping accuracy in MTI data spanning multiple batches.

To further evaluate the performance of UniFORM in batch normalization, we used kBET analysis on MTI data generated from multiple batches. kBET implicitly assumes that samples across batches share similar cellular compositions to accurately assess batch effects without being confounded by biological variability. To ensure a fair kBET analysis, we selected samples with comparable cellular compositions from different batches. Specifically, we focused on five groups with distinct cellular populations to maintain heterogeneity across the selected samples (**Table S2**). We evaluated batch effect removal using three metrics: kBET acceptance score, average Chi-square statistics, and average p-value (see **Methods**). A higher kBET acceptance score indicates better mixing of similar cell types, reflecting more effective batch correction. The Chi-square statistic measures the magnitude of batch imbalance, with lower values indicating better batch mixing. The average p-value reflects the statistical significance of batch mixing, with higher values indicating closer alignment to an ideal batch-free distribution.

As demonstrated in **Figure 2b**, group 1 (PRAD02 & PRAD14) exhibits significant population differences, with one sample much larger than the other. This discrepancy artificially inflated the kBET score. However, UniFORM still achieved the best performance in all metrics, highlighting its robustness even with imbalanced datasets. For the remaining groups, where samples exhibit similar population size and cell composition but also significant batch effect, UniFORM outperformed other methods by a significant margin (**Table 1** and **Table S2**), with the highest average kBET score of 0.60843 (raw=0.00098, MxNorm=0.38788, Z-score=0.28570, ComBat=0.27266, meandiv=0.54240), lowest average Chi-square statistics of 6.4473 (raw=20.093, MxNorm=9.9575, Z-score=14.823, ComBat=15.298, meandiv=7.6324), and highest average p-value of 0.25835 (raw=0.00145, MxNorm=0.15307, Z-score=0.11459, ComBat=0.10981, meandiv=0.22568). We have further demonstrated UniFORM’s effective batch correction through UMAP visualization shown in **Figure S4**.

**Table 1:** Summary of the average statistics of the kBET analysis. UniFORM outperformed all other methods by a significant margin in all three kBET evaluation statistics.

| Method | Average kBET Score | Average Chi-square Statistics | Average p-value |
| --- | --- | --- | --- |
| **UniFORM** | **0.60843** | **6.4473** | **0.25835** |
| Raw | 0.00098 | 20.093 | 0.00145 |
| MxNorm | 0.38788 | 9.9575 | 0.15307 |
| z-score | 0.28570 | 14.823 | 0.11459 |
| ComBat | 0.27266 | 15.298 | 0.10981 |
| MeanDiv | 0.54240 | 7.6324 | 0.22568 |

### Feature-Level Normalization: Positive Population Percentage Preservation

Effective normalization should adjust for technical variations and biases without altering the inherent biological distribution of marker expression. To evaluate this, we compared the percentage of positive populations in samples before and after normalization. The primary goal of normalization is to establish a reliable global threshold across samples that accurately differentiate positive and negative populations while maintaining the integrity of the positive population percentage. This consistency is essential for downstream analyses, particularly in cell phenotyping, where a global threshold ensures that any subsequent classification of cells into phenotypic categories remains consistent, reducing variability and minimizing bias from sample-specific thresholds.

To achieve this, we applied a two-component Gaussian Mixture Model^24^ (GMM, see **Methods**) to determine local thresholds for each raw sample and a global threshold for the normalized samples in each marker of the CRC-ORION and PRAD-CyCIF dataset. The mean percentage change (see **Methods**) was then calculated by comparing the local thresholds and the global threshold. While perfect preservation of the positive population percentage is unlikely, significant changes should be avoided to ensure data integrity.

UniFORM effectively preserves population levels from local thresholds (pre-normalization) to global thresholds (post-normalization), with minimal percentage changes and low standard deviation across all markers in both datasets. In contrast, other methods exhibited substantial shifts in thresholds, high mean percentage changes, and significant standard deviations (**Figure 2c, Figure 2d**). Notably, MxNorm and mean division resulted in pronounced percentage changes for key markers such as CD45, ECAD, and Pan-CK in the CRC-ORION dataset.

Similarly, all methods except UniFORM produced high percentage changes and variability for multiple markers in the PRAD-CyCIF dataset, particularly aSMA, CD20, and NOTCH1. These high standard deviations indicate inconsistent performance across markers and samples, reflecting poor alignment of distributions and signal distortion that adversely affected the GMM thresholding (**Table S4a, Table S4b, Figure S6a, Figure S6b**). Markers such as EPCAM and CD45 in the PRAD-CyCIF dataset also exhibited high mean percentage changes across all methods, which is likely attributable to their intrinsic biological heterogeneity across samples. Certain samples have a pronounced positive population, while others exhibit negligible positive populations. For samples with low positive populations, applying a local GMM threshold can lead to biases, as the minimal positive population may skew the threshold estimation. In contrast, a global GMM threshold, calculated across all samples, provides a more reliable and consistent threshold. However, the global threshold may differ from the local thresholds, particularly in heterogeneous datasets, contributing to the observed percentage change.

### Feature-Level Normalization: Impact on Downstream Analysis

To evaluate the impact of UniFORM normalization on downstream analysis, we performed UMAP visualizations and Leiden clustering (see **Methods**). As shown in **Figure 3a**, the pre-normalization UMAP revealed distinct sample separation, indicating the presence of batch effects across samples. In contrast, post-normalization UMAP embeddings showed a well-mixed distribution of samples, demonstrating that UniFORM effectively minimized batch effects. To further assess whether similar cell types from different batches clustered together after UniFORM, we examined the UMAP plots color-coded by the intensity values of key markers (CD45, αSMA, and ECAD). In the raw data, high-intensity regions for these markers were dispersed across multiple clusters, indicating batch-induced variability in marker expression.

**Figure 3:**
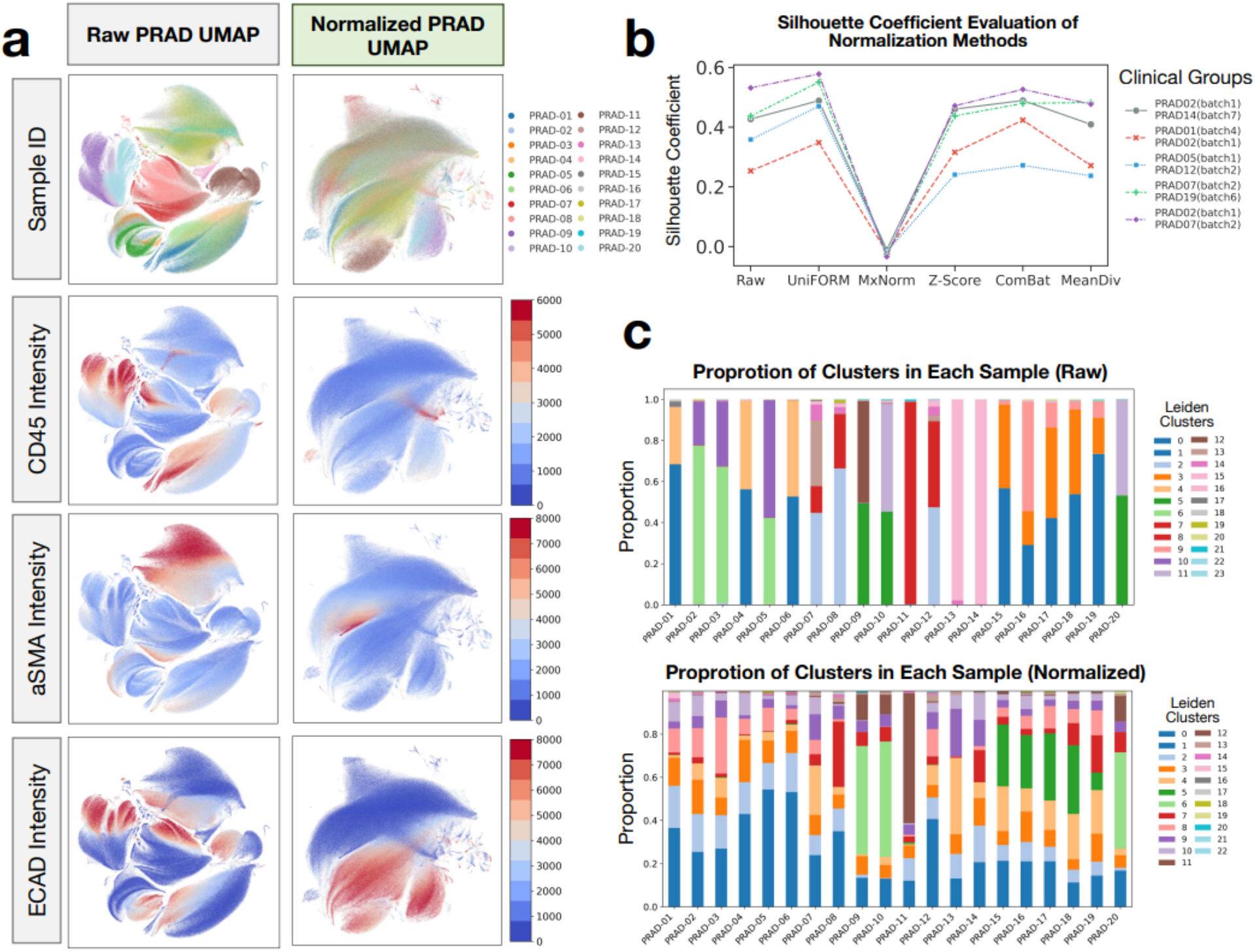
Impact of UniFORM on downstream analysis. **(a)** UMAP of raw and normalized PRAD-CyCIF dataset colored by sample IDs showing mixture after normalization and UMAP of raw and normalized PRAD-CyCIF dataset colored by the intensity value of marker CD45, α-SMA, and ECAD showing clustering of higher intensity regimen after normalization. (**b**) Silhouette coefficient evaluation of mutually exclusive marker separation with five patient groups, a higher Silhouette score indicates better cluster separation. **(c)** Leiden clustering of raw and normalized PRAD-CyCIF dataset showing enriched cluster proportions across samples after normalization.

After normalization, these high-intensity regions clustered together, highlighting that UniFORM successfully aligned positive cell populations across batches, supporting both improved normalization across samples.

Building on these qualitative findings, we next sought to quantitatively evaluate whether UniFORM preserved biological relationships between distinct cell types. Specifically, we assessed the preservation of mutual exclusivity among tumor, immune, and stromal cell types after normalization, we performed Silhouette score analysis across normalization methods. Silhouette scores measure clustering quality by comparing intra-cluster compactness and inter-cluster separation, with higher scores indicating better-defined, more separable clusters. Given the biological distinctness of epithelial (ECAD+), immune (CD45+), and stromal (αSMA+) cells, we expect normalization methods that preserve mutual exclusivity to yield higher Silhouette scores.

Using the same patient samples as the kBET analysis, we found that UniFORM achieved the highest average Silhouette score of 0.48688 (raw=0.40137, MxNorm=-0.02141, Z-score=0.38513, ComBat=0.43796, meandiv=0.37537), indicating best preservation of biological separation between these cell types (**Figure 3b, Table 2**). Mean division also performed well. Interestingly, ComBat achieved slightly higher Silhouette scores than UniFORM for two sample pairs (**Table S3**); however, this is primarily attributed to ComBat’s tendency to compress the data range, artificially reducing intra-cluster distances and inflating scores. Supporting this, scatter plots of mutually exclusive marker pairs (ECAD vs. CD45 and αSMA vs. ECAD) show that UniFORM and mean division effectively preserve the expected L-shaped mutually exclusive patterns (**Figure S5**), reinforcing biological interpretability. In contrast, methods such as MxNorm (registration option), Z-score, and ComBat distort marker relationships, weakening mutual exclusivity and potentially compromising downstream analyses. Notably, MxNorm yielded a negative Silhouette score, consistent with observed misalignments—such as instances where positive peaks align with negative peaks—potentially impacting mutual exclusivity and reducing biological interpretability.

**Table 2:** Summary of the average statistics of the Silhouette analysis. UniFORM outperformed all other methods the average Silhouette score statistic.

| Method | Average Silhouette Coefficient |
| --- | --- |
| **UniFORM** | **0.48688** |
| Raw | 0.40137 |
| MxNorm | -0.02141 |
| z-score | 0.38513 |
| ComBat | 0.43796 |
| MeanDiv | 0.37537 |

For the Leiden clustering analysis, we calculated the distribution of cluster proportions across samples before and after normalization. As shown in **Figure 3c**, before normalization, cluster proportions were highly skewed, with many samples dominated by a single cluster or containing only a limited subset of clusters. This pattern indicates sample-specific biases, likely driven by batch effects and technical variability, which masked the true biological heterogeneity. After normalization, cluster representation across samples became more balanced, with each sample contributing more uniformly to the overall clustering structure. This improved cluster consistency suggests that UniFORM successfully reduced technical artifacts, enabling more meaningful and biologically relevant clustering. This ensures clustering was driven by intrinsic biological variation rather than batch effect, which enhances downstream analyses, such as cell type annotation, and allows for more accurate comparisons between experimental conditions.

### Feature-Level Normalization: Computational Efficiency

To assess computational efficiency, we benchmarked UniFORM and MxNorm by running each method five times on the PRAD-CyCIF dataset under identical conditions (Apple MacBook Pro with an M2 Max chip and 32GB RAM). For a fair comparison, the registration-based option of MxNorm was selected, as it also uses functional data registration. UniFORM demonstrated significantly faster performance, with an average runtime of 30.58 ± 0.30 seconds, compared to 114.75 ± 4.35 seconds for MxNorm (**Figure S7**), representing nearly a fourfold improvement in efficiency.

### Evaluation of Pixel-Level Normalization Methods

A major novel contribution of UniFORM is the introduction of a robust pixel-level normalization workflow (see **Methods**), which effectively aligns marker signals while preserving distribution shape. Similar to feature-level normalization, this approach applies the normalization factor at the pixel level. UniFORM uses only pixels within the cell/nuclei segmentation masks, excluding non-cell regions to yield a cleaner and more accurate representation of the true signal, thereby enhancing the accuracy of signal alignment. UniFORM processes unsigned 16-bit images—the standard format for MTI images—and implements a mechanism to prevent range compression or overflow issues that can occur when the image is multiplied by a normalization factor and scaled back to 16-bit again. Consequently, the normalized images are also saved as 16-bit files without any unwanted artifacts.

### Pixel-Level Normalization: Intensity Distribution Alignment

Following our feature-level evaluation, we performed the automatic normalization on the CRC-ORION and PRAD-CyCIF pixel-level datasets and evaluated the efficacy of signal alignment. UniFORM demonstrated consistent effectiveness in aligning the pixel intensity distribution to the cell mean intensity distribution. **Figure 4a** shows that both PD-L1 and ECAD achieved good alignment while preserving the original shape of the distribution, and the rest of the markers also achieved good alignment (**Figure S8a-b, Figure S9a-b**).

**Figure 4:**
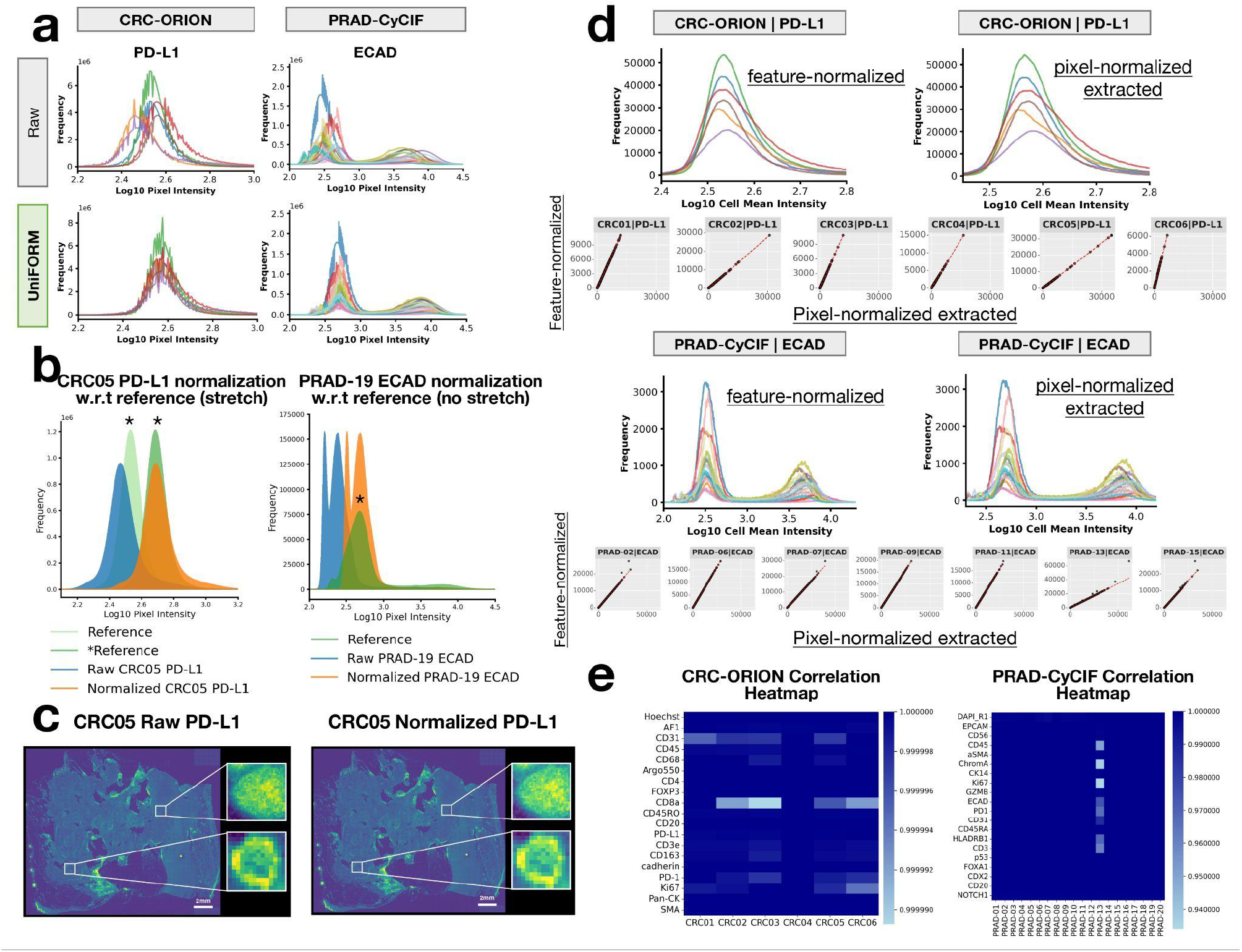
Pixel-level normalization evaluation. **(a)** Evaluation of UniFORM’s pixel intensity signal alignment of unimodal marker PD-L1 and bimodal marker ECAD **(b)** Demonstration of the sample signal alignment with respect to its reference and the effect of stretching mechanism. **(c)** Visual inspection of the PD-L1 channel before and after normalization confirms no artifacts are introduced. **(d)** Features extracted from normalized images achieve good signal alignment and almost perfect Spearman correlation with the feature-level normalized data. **(e)** Spearman correlation heatmap showing extracted features having almost perfect correlation with the feature-level normalized in both CRC-ORION and PRAD-CyCIF data. Refer to **Figure 2a** for legend information.

Pixel-level normalization differs from feature-level normalization in how the normalization factor affects data distribution. When a normalization factor < 1 is applied at the pixel level, the marker intensity distribution shifts left, resulting in a narrower range. The conversion back to uint16 introduces compression artifacts (‘spikes’) due to the limited intensity values representable within the fixed-bit range. **Figure S10** illustrates this effect, showing how the unstretched distribution displays prominent spiking artifacts from quantization, in contrast to the smoother, properly scaled (stretched) distribution. In contrast, feature-level normalization, which is maintained in float64, avoids this issue and preserves data precision. To address this compression issue, we incorporated a linear stretching mechanism (see **Methods**), applying a stretching factor to each sample to restore its range to pre-normalization levels. This approach ensures even the most compressed samples avoid artifacts associated with data compression while maintaining the original distribution shape.

**Figure 4b** illustrates the scenarios where a stretching factor is needed versus where it is not. For PD-L1, the raw CRC05 distribution (blue) is expected to align with the reference (light green). However, due to leftward shifts in some samples, a common stretching factor is calculated and applied uniformly to all samples, including the reference, after the normalization step. This process aligns the distributions while linearly stretching them preserving the original distribution shape. For ECAD, where the reference is already the right-most sample, no stretching is required as all distributions are shifted to the right. The reference sample remains unchanged (i.e., no stretching factor), and the raw PRAD-CyCIF sample aligns effectively with the reference distribution. In both cases, the signals align accurately with their references, addressing compression and overflow issues and ensuring robust pixel-level normalization.

To further illustrate that no image artifacts are introduced after normalization, we examined the PD-L1 channel of CRC05 before and after normalization. The results clearly show that no artifacts were introduced during the process, and the single-cell granularity was perfectly preserved (**Figure 4c**).

### Pixel-Level Normalization: correlation with feature-level normalization

Finally, to assess the relationship between features extracted from pixel-level normalized images and those from the feature-level normalized dataset, we computed Spearman correlations and plotted the line of best fit (see **Methods**). This analysis was performed using the PD-L1 marker from the CRC-ORION dataset and ECAD marker from PRAD-CyCIF dataset. The features extracted from pixel-level normalized images in both datasets showed great alignment comparable to those from the feature-level normalized dataset, displaying a linear relationship with near-perfect Spearman correlations (**Figure 4d**). These results indicate that normalizing the pixel-level image dataset also effectively normalizes the feature-level mean intensity dataset. Furthermore, the strong correlation (**Figure 4d, 4e**) suggests that features derived from normalized images consistently maintain the positive population percentage.

## Discussion

MTI datasets, both at pixel and feature levels, often exhibit right-skewed distributions, largely influenced by commonly observed trends. This skewness primarily arises from the biological nature of marker expression, where a subset of cells exhibits high-intensity signals while the majority show lower or background levels. These high-intensity signals contribute to the long tail of the distribution that varies across samples due to heterogeneous expression patterns inherent in tissue samples. The four benchmarked methods—MxNorm (registration option), Z-score, ComBat, and Mean division struggle to effectively account for the interplay of biological variability, technical noise, and distribution skewness. Consequently, these methods frequently misalign signals, distort original distributions, and change the proportions of positive populations, highlighting the need for approaches tailored to the unique characteristics of MTI data. A common limitation across existing normalization methods is the lack of explicit discrimination between negative and positive expression peaks. This non-selective adjustment risks overlapping biologically distinct populations — shifting positive populations into the range of negative populations and vice versa — which can severely confound downstream analyses such as cell type classification, phenotyping, or spatial neighborhood inference.

MxNorm addresses the challenge of evaluating normalization quality in the absence of a universal standard by benchmarking various methods, including log transformation, ComBat, mean division, and functional data registration. Based on its evaluation, MxNorm recommends mean division as the most practical default for MTI data, while suggesting that combining mean division with log transformation and functional data registration can further improve normalization, especially for correcting marker-specific discrepancies. For example, the authors cite an outlier case with CD8 where mean division alone was insufficient, but the combined approach resolved the issue.

To benchmark UniFORM—our method that also employs functional data registration—we compared it to MxNorm’s mean division and registration pipelines. The MxNorm registration method applies functional data registration using B-spline basis functions to align marker intensity histograms across samples. Specifically, each sample’s histogram is first smoothed and represented as a cubic functional data object. A monotonic warping function, parameterized by low-degree B-splines, is then estimated to align each curve to a target distribution (typically the mean distribution across samples). While this approach makes no explicit assumption of data normality or symmetry, it faces fundamental limitations when applied to the distributional characteristics of MTI data, which are often unimodal, right-skewed distributions with a large negative population and a long, sparse positive tail.

In these cases, the smooth warping function tends to compress or distort the long tail, altering the relative intensity of the positive population and disrupting the biologically meaningful separation between negative and positive cells. This distortion can significantly affect threshold-based classification and downstream phenotyping. Furthermore, the smoothness constraint imposed by the low-degree B-spline basis may distort the peak-to-tail ratio of the dominant negative population, further compromising biological interpretability.

Although the authors of the MxNorm paper report that the B-spline registration performs well for bimodal distributions, our evaluation showed the opposite: MxNorm failed to align complex, bimodal distribution such as ECAD, especially in heterogeneous samples. While the method may work adequately for simpler, unimodal markers like PD-L1, it struggles with more complex distributions. This is evidenced by the severe distribution distortion in **Figure 2a**, poor kBET and Silhouette score (**Figure 2b, 3b**), and the substantial shift in positive population percentage changes (**Figure 2c**).

Z-score normalization, while widely used for its simplicity, relies on assumptions of normally or symmetrically distributed data and consistent cell composition and signal distribution across samples. However, MTI data often show inherent right skewness and heterogeneity, making Z-score normalization less suitable. In right-skewed data, the mean is shifted toward higher values due to the tail. When samples have consistent cell composition (as with PD-L1 in previous examples), Z-score normalization may perform adequately. However, when cell composition varies significantly (as seen with ECAD), each sample’s reference point (mean) and scale (standard deviation) can be very different, resulting in misalignment. Furthermore, in right-skewed data, Z-score normalization causes lower-intensity data points to become overly compressed around zero, while high-intensity points may become exaggerated. This distorts the natural spread and distribution of data, causing large positive population percentage changes as evident in **Figures 2a** and **2c**.

Similar to Z-score normalization, ComBat assumes a common distribution and consistent cell composition across samples, with additive batch effects that can be linearly modeled. However, the right skewness, heterogeneous positive signals, and non-uniform technical variation in MTI data violate these assumptions. As a result, when the samples have significantly different cell compositions, ComBat may struggle to differentiate between technical batch effects and biological variation, potentially resulting in over-correction or under-correction. Additionally, ComBat can alter the overall distribution of data, particularly when data deviate from normality, evident in **Figure 2a** and **2c**, making it inherently unsuitable for MTI normalization.

The mean division transformation, while simple and computationally efficient, assumes that inter-sample variation arises primarily from uniform multiplicative scaling and that the sample mean serves as a reliable reference across images. However, this assumption fails in MTI data, where marker intensity distributions are typically right-skewed and cell compositions vary substantially between tissue samples. In such distributions, the mean is heavily influenced by the small subset of high-intensity (positive) cells, making it an unstable and biologically unrepresentative normalizing factor. Consequently, mean division can distort biologically meaningful differences—particularly when the proportion of marker-positive cells varies across samples—leading to misaligned distributions and inaccurate estimates of positive population estimate (**Figure 2a, 2c**). Although mean division may yield higher kBET acceptance rates by enforcing apparent batch mixing, this improvement often reflects artificial homogenization rather than true correction, as it suppresses biological variability in favor of uniformity.

FLINO uses a grid-based quantile normalization (Q75) approach, dividing each image into multiple grids and adjusting raw intensities using a multiplicative scaling factor. For each grid, raw intensity is normalized by multiplying it with the ratio of a global quantile (calculated across all images) to a local quantile (specific to the grids in each sample). This aligns intensities to a shared global quantile reference, assuming all samples share a common intensity distribution at the chosen quantile. However, this assumption falters with right-skewed and heterogeneous MTI marker distributions. For right-skewed markers, the Q75 value, influenced by a long tail of high-intensity values, poorly represents the central trend, leading to overcorrection in low-intensity grids or undercorrection in high-intensity grids. Additionally, FLINO does not distinguish between negative peaks (background) and positive peaks (biological signals), risking distortion of key biological features. Normalizing each grid independently can also amplify batch effects or local intensity differences, introducing artificial variability or failing to correct pre-existing inconsistencies. From a computational standpoint, FLINO’s R-based workflow and reliance on dividing images into grids make it resource-intensive, inefficient, and impractical for modern MTI datasets, which often contain millions of pixels and thousands of grids per slide.

Our universal normalization pipeline, UniFORM, leverages the negative population as a biologically invariant reference point and applies rigid functional data registration to align distributions, is free from restrictive assumptions. This makes UniFORM broadly applicable across diverse MTI datasets. By anchoring normalization on negative peak alignment, UniFORM effectively reduces technical variation while preserving biologically meaningful variability. The use of automated rigid landmark-based registration ensures that distributions are shifted consistently across samples without introducing distortion. **Table 3** summarizes and highlights that UniFORM is the most comprehensive method, outperforming others in all evaluating metrics and effectively correctly both inter-batch and intra-batch technical variations. However, the effectiveness of UniFORM depends on the quality of the MTI dataset and its cell/nuclei segmentation masks used to extract signals. In cases where imaging artifacts such as autofluorescence severely compromises the signal-to-noise ratio, it may become difficult or even impossible to reliably recover the true biological signal. In such instances, we recommend excluding those samples during the quality control (QC) process, as normalization cannot compensate for fundamentally compromised data. For reliable cell/nuclei segmentation, we employed the MESMER cell segmentation model^25^ (see **Methods**), a state-of-the-art tool, to generate these masks. For datasets with minimal cell loss, like CRC-ORION, we used the first-round DAPI channel and corresponding morphology channels to extract the masks. For datasets with significant cell loss in later staining rounds, like the PRAD-CyCIF dataset, we adjusted the masks to account for tissue loss by comparing the first-round DAPI masks with the last-round masks, ensuring cleaner and meaningful biological signals.

**Table 3:** Comparison of normalization methods analyzed in this study. UniFORM outperformed existing methods across all evaluated criteria, demonstrating robustness and effectiveness. An asterisk (*) denotes features claimed by the authors but lacking implementation or comprehensive evaluation.

| Methods | Feature-level normalization | Align correct signals | Preserve distribution | Preserve population counts | Pixel-level normalization |
| --- | --- | --- | --- | --- | --- |
| **UniFORM** | ✓ | ✓ | ✓ | ✓ | ✓ |
| MxNorm | ✓ | ✓ | ✗ | ✗ | ✗* |
| Z-Score | ✓ | ✗ | ✗ | ✗ | ✗ |
| ComBat | ✓ | ✗ | ✗ | ✗ | ✗ |
| MeanDiv | ✓ | ✗ | ✗ | ✗ | ✗ |
| FLINO | ✗ | ✗ | ✓ | ✗ | ✓ |

Normalization methods in MTI and single-cell analysis often face a trade-off between achieving a good mixture of cells (as indicated by kBET acceptance rates) and preserving biological signal^26–29^. In our study, UniFORM demonstrated the highest kBET metrics across all groups, outperforming other methods that rely on artificial homogenization to improve the mixture at the cost of destroying biological integrity. This highlights UniFORM’s superior ability to achieve both robust mixing and signal preservation, setting it apart as a more balanced and effective normalization approach.

In certain cases, such as the CD45 marker in the PRAD-CyCIF dataset, marker intensity distributions exhibit highly heterogeneous and irregular patterns, such as multimodal distributions or inconsistent positive-to-negative population ratios (**Figure S11a**). These irregularities violate the assumptions of the automatic normalization workflow and are automatically flagged using a Kullback–Leibler (KL) divergence-based metric, prompting the user to consider manual fine-tuning. To address this challenge, we implemented a guided manual fine-tuning option (see **Methods, Figure S11b**), which enables user-directed landmark selection on histogram representations of marker intensity distributions. This process allows users to manually adjust the position of key landmarks—typically corresponding to the negative population peak—to improve alignment in cases where automated registration may be suboptimal. This feature demonstrates the flexibility of the pipeline, enabling accurate signal alignment even in the most complex cases where automated methods alone may fall short.

A limitation of our normalization pipeline is its reliance on the linear application of normalization factors. While this approach is well-suited to IF-based techniques such as CyCIF and CODEX, which exhibit a linear relationship between signal intensity and antigen concentration, it is unsuitable for multiplexed immunohistochemistry (mIHC). In mIHC, chromogenic detection produces non-linear signal intensities because of enzyme-catalyzed colorimetric amplification, which disrupts the proportionality between signal intensity and antigen concentration^2–4^.

Applying a linear normalization factor to mIHC data can lead to misrepresentation, as the scaling may unevenly affect intensity levels, exaggerating or diminishing the actual signal distribution. To address this, a warping function might be needed to perform a non-linear normalization of the mIHC samples. Additionally, while UniFORM is currently optimized for within-cell normalization, the underlying framework could be adapted for extracellular regions with appropriate modifications, enabling broader application to spatial compartments beyond cell boundaries. Future work could explore this option.

Ultimately, this work demonstrates the strength of UniFORM as a versatile normalization pipeline, balancing automatic approaches with manual flexibility to achieve precise signal alignment, across diverse MTI data sets and cancer types. By addressing key challenges in batch correction and normalization, UniFORM sets the foundation for accurate, reproducible analysis of multiplex imaging data, with the potential to significantly improve downstream biological insights and clinical applications.

## Methods

### Dataset

Detailed information on CRC-ORION, PRAD-CyCIF, and TMA-Lunaphore datasets is outlined in **Table S1a-c** and the **Data Availability** section.

### Preprocessing

The CRC-ORION has already been processed and made available online^30^. Preprocessing steps of the PRAD-CyCIF and TMA-Lunaphore dataset are detailed in our previous publication^31^. All three datasets are prepared in the ome.tif format. Mesmer cell segmentation is used to segment cells and nuclei for both datasets. Nuclear marker DAPI, and membrane markers (a composite projection of Pan-cytokeratin and CD45 marker) were used for the CRC-ORION dataset. Nuclear marker DAPI, and membrane markers (a composite projection of EPCAM, CD45, and ECAD marker) were used for the PRAD-CyCIF and TMA-Lunaphore dataset. The masks were registered and refined to ensure each nucleus had a corresponding cell membrane. The cell ID, centroids, and cell mean intensities were extracted with scikit-image^32^.

### Quality control

Quality control procedures were implemented to account for tissue loss and ensure the removal of imaging defects and artifacts. Once the masks were refined to ensure each cell had a corresponding nucleus, a binary mask from the last-round nuclear mask was used to rapidly filter cells based on the overlap with the first-round nuclear mask. Cells were considered lost if their mean intensity overlapped with the last-round nuclear mask and fell below a specified threshold. We used 0.10 in our pipeline to remove the lost cells and maintain the cells that might have drifted mildly due to local tissue deformation. The filtered masks were processed to ensure that nuclear and cell masks maintained consistent IDs. Code to perform tissue loss filtering is available along with our normalization pipeline. Besides tissue loss, other common imaging defects and artifacts include tissue folding and wrinkling, tissue disintegration, significant autofluorescence, bright spots due to dye aggregation, and air bubbles and debris. Samples with these artifacts were either dropped or cropped depending on the severity.

### UniFORM: Feature-level normalization

UniFORM’s feature-level normalization pipeline can be divided into three steps: 1. Calculate histogram, 2. Automatic functional data registration, 3. Normalization. Feature-level normalization takes in the raw cell mean intensity data, undergoes the automatic max-correlation based functional data alignment, and then outputs the normalized cell mean intensity data.

#### 1. Calculate histogram

For each marker sample, we apply a logarithmic transformation step to enhance dynamic range and skewness. Then for each marker, it iterates through samples to determine a common global range, this ensures a uniform and consistent comparison across samples and avoids confounding effects due to varying axis scales. After a common global range is established, it then iterates through samples to produce histograms along with plots that provide an essential visualization of raw data distribution.

#### 2. Automatic functional data registration

In the second step of the automatic normalization pipeline, we align the distributions of cell mean intensities across samples using a Fast Fourier Transform (FFT)-based max-correlation algorithm. This approach ensures that for each marker, all sample histograms are aligned to a selected reference histogram. The reference sample is chosen to be representative of the marker’s characteristics and serves as the baseline for alignment.

The algorithm works by computing the cross-correlation between each sample’s histogram and the reference histogram to determine the optimal shift that maximizes their alignment. To facilitate this process, histograms are first transformed into functional data objects (FDataGrid), converting discrete histograms into continuous functional representations. This transformation allows for smooth mathematical operations and precise functional alignment.

The FFT-based cross-correlation efficiently identifies the shift that maximizes similarity between the sample and reference histograms. The calculated shifts are then applied to realign the sample histograms to the reference, standardizing the distribution of cell mean intensities across all samples. The algorithm is summarized in the pseudocode below:

##### Algorithm 1

Correlation-Based Normalization Using FFT

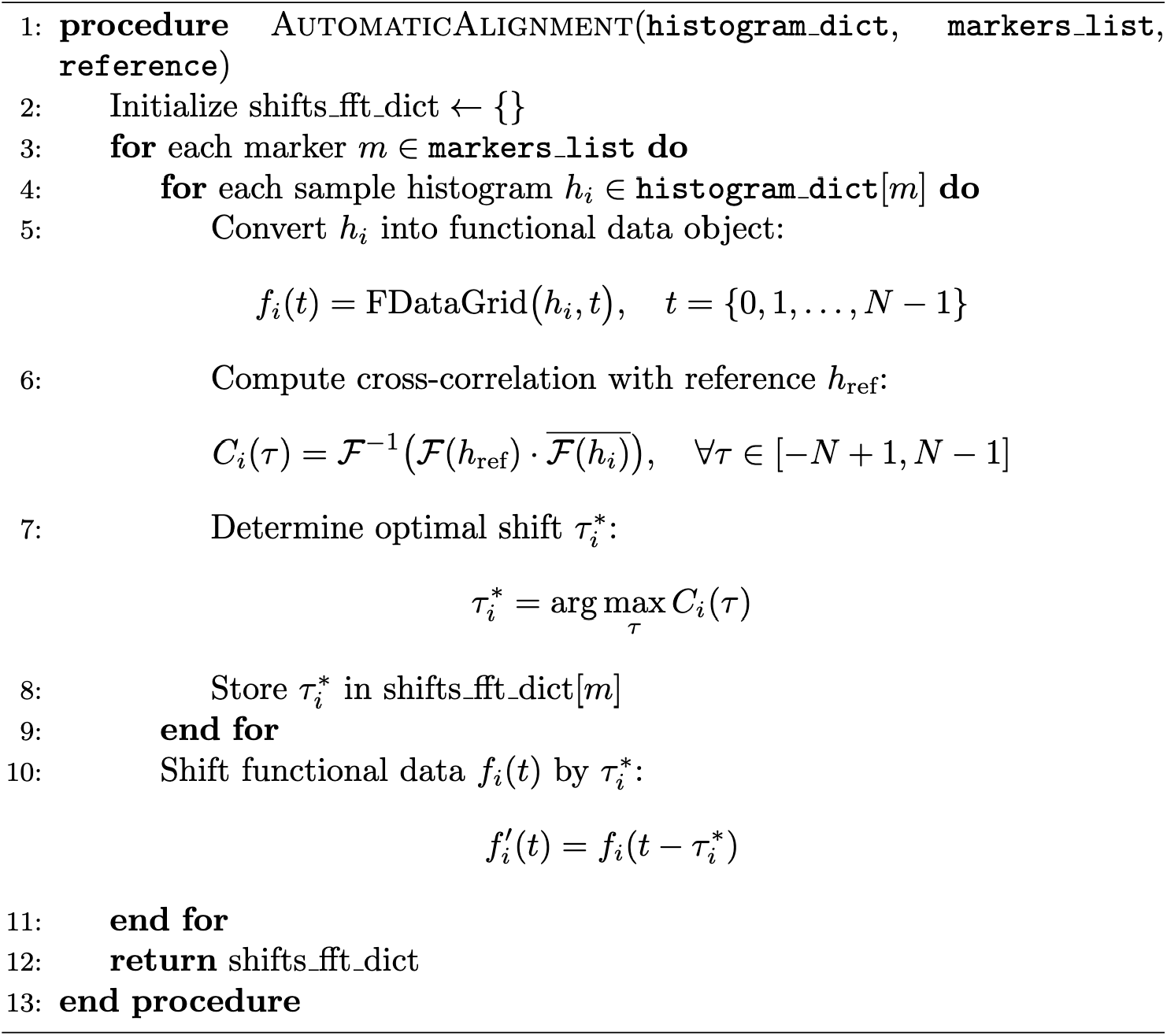

#### 3. Normalization

In step 3 of the normalization pipeline, we apply the normalization factors derived from FFT-based shifts to the raw data to achieve normalization across all samples. This is achieved by first translating the normalization factors in the FData scale to the logarithmic scale to ensure these shifts correspond accurately to the scale of the log-transformed raw data. Then for each sample and marker, the normalization factor is applied by scaling the original marker intensity data *e*^*shift*^to make sure the shift is applied at the scale of the raw data. The normalized data is then saved as float64 data type. The algorithm can be described by the pseudocode below:

##### Algorithm 2

Performing Data Normalization

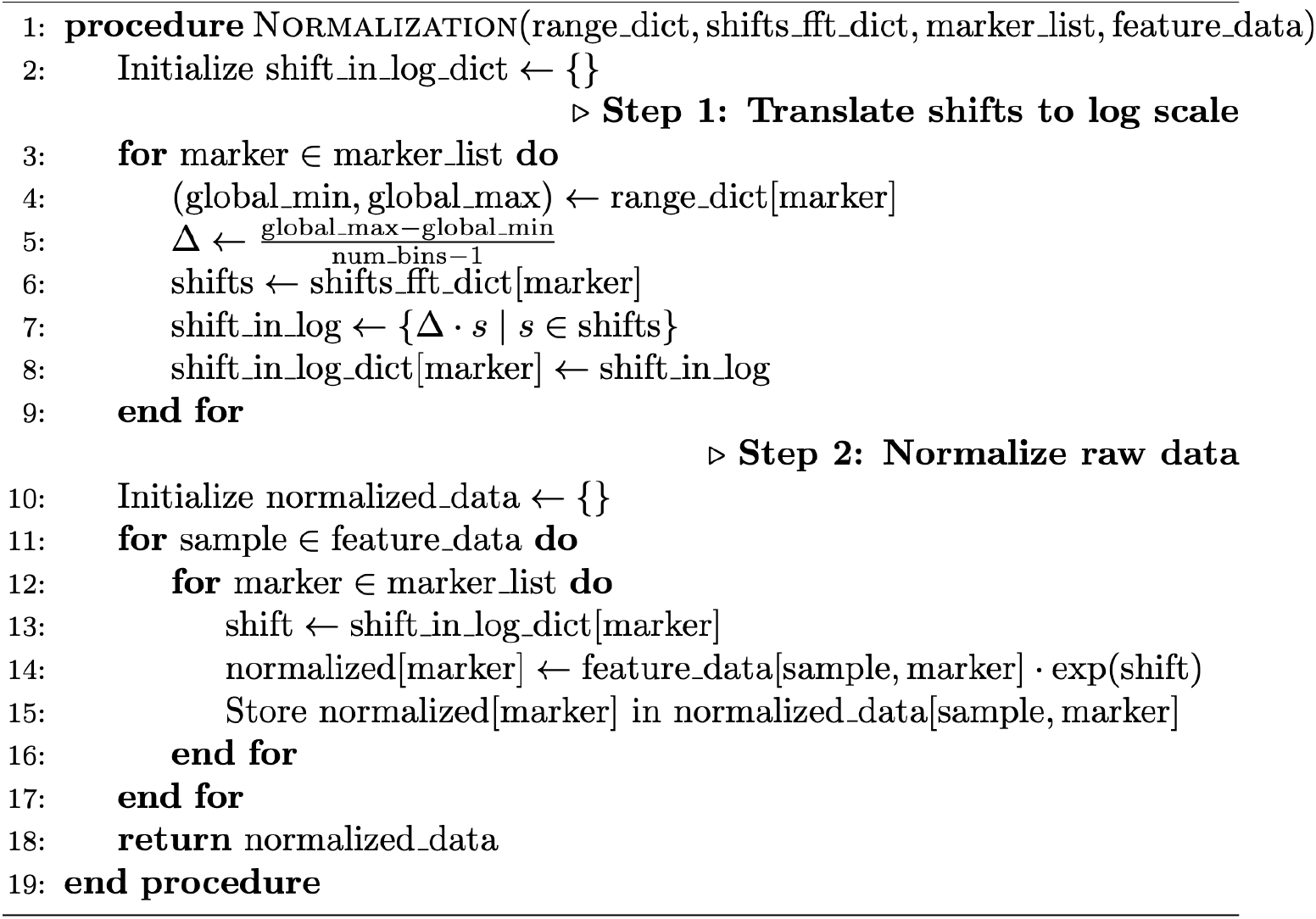

### UniFORM: Pixel-level normalization

Pixel-level normalization follows a very similar workflow to feature-level, differing only in how the input is processed and how the normalization factor is applied to the raw image data. The pixel-level pipeline handles raw pixel data in the format of Dask arrays for computational efficiency and uses cell/nuclei segmentation masks to filter out non-relevant areas and ensure that only pixels corresponding to cells or nuclei are used for the histogram calculation. Then it undergoes a similar workflow of histogram calculation and automatic normalization factor estimation. However, the two workflows differ in how the normalization factor is applied to the raw data. As described in the results section, compression and overflow issues are specific to pixel-level data as they need to be maintained in uint16 after normalization.

The stretching mechanism in the pixel-level normalization pipeline is designed to address compression artifacts that arise when applying normalization factors < 1 to raw pixel data. When pixel intensity values are scaled-down and converted back to uint16, the same number of unique intensity values needs to be rounded to a narrow range, this restriction can introduce artifacts causing spikes in the histogram where multiple original values round to the same output value. To determine the stretching factor, for each marker, the most compressed normalization factor (corresponds to the sample with the most left-shift) is determined, and we will be able to calculate a stretching factor that ensures the range of compressed data can be expanded to its pre-normalization level. All normalized samples will be linearly stretched by this stretching factor for consistency. The algorithm can be described by the pseudocode below:

#### Algorithm 3

Pixel-level Normalization and Stretching Mechanism

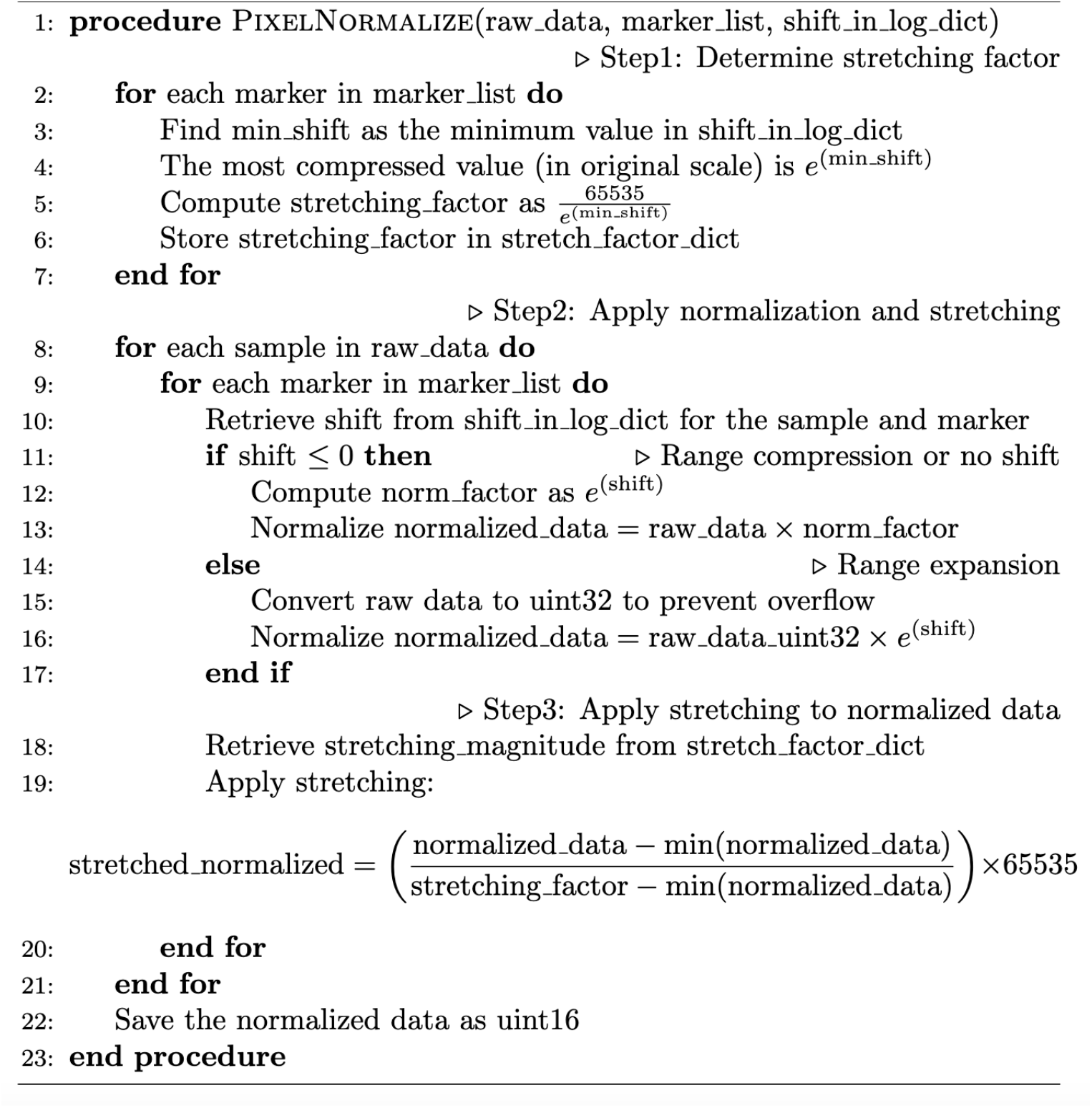

The normalized pixel-level data is saved as an uint16 image stack in the ome.tiff format, the same as the input image format.

UniFORM performs pixel-level normalization exclusively within nuclear or cellular segmentation masks. This design choice is intentional: including background or non-cellular regions would introduce substantial bias into the intensity distribution, especially in the negative population, which we rely on as a biologically invariant reference for normalization. By focusing on pixels within cell boundaries, we ensure that the normalization process reflects true biological signal and avoids distortion from non-informative background.

### UniFORM: Manual fine-tuning option (landmark normalization)

The manual landmark normalization step in the pipeline allows for normalization adjustment based on selected landmarks that represent user-defined negative peaks. These landmarks are critical points chosen by the user, either manually or with guided assistance using Gaussian Mixture Model (GMM) curves as a visual aid. The process starts by plotting functional data curves of the histogram distribution for each marker on an interactive *plotly* plot (https://github.com/plotly/plotly.py), allowing the user to quickly and easily identify landmarks.

The landmark function calculates the shift needed to align the functional data curves of each sample to the reference negative peak. If landmarks are provided for a marker, the landmark function adjusts the curves so that the user-defined locations become aligned with the user-defined reference. If landmarks are not provided, the pipeline uses the automatic max-correlation based method.

### Implementation of normalization benchmarking techniques

* **MxNorm Normalization:** We followed the guidelines on the MxNorm Github repository: https://github.com/ColemanRHarris/mxnorm to conduct normalization. We used the best-reported normalization methods for analysis, specifically transform = “log10_mean_divide” and method = “registration” (meaning that performing a log10 and mean division transformation of the data before performing B-spline function data registration).
* **ComBat Normalization:** Our ComBat normalization is implemented using pyComBat, a new Python implementation of ComBat algorithm to adjust for batch effects within our data. Prior to applying ComBat, a log10 transformation with a small offset was applied to the data. We followed the guidelines on pyComBat website: https://epigenelabs.github.io/pyComBat/ to conduct normalization.
* **Z-score Normalization:** We perform Z-score normalization on the sample level using Z-score implementation from SciPy (scipy.stats). Prior to applying the Z-score, a log10 transformation with a small offset was applied to the data.

Mean Division Normalization: We performed mean division normalization using the MxNorm package by setting transform = “mean_divide” and method = “none”

### Gaussian Mixture Model (GMM) Analysis

2-component GMMs, statistical methods that fit two Gaussian distributions to data to identify underlying subpopulations, were used in our study. We adopted the SCIMAP^24^ implementation, which uses a simple GMM model, for visualization purposes in Figure 2a. For more accurate positive population thresholding (in our positive population change analysis), we implemented a more complex GMM, which uses a more complex model initiation and uses the Gaussian probability density functions (PDFs) to determine the best threshold for positive population. The algorithm can be described by the pseudocode below:

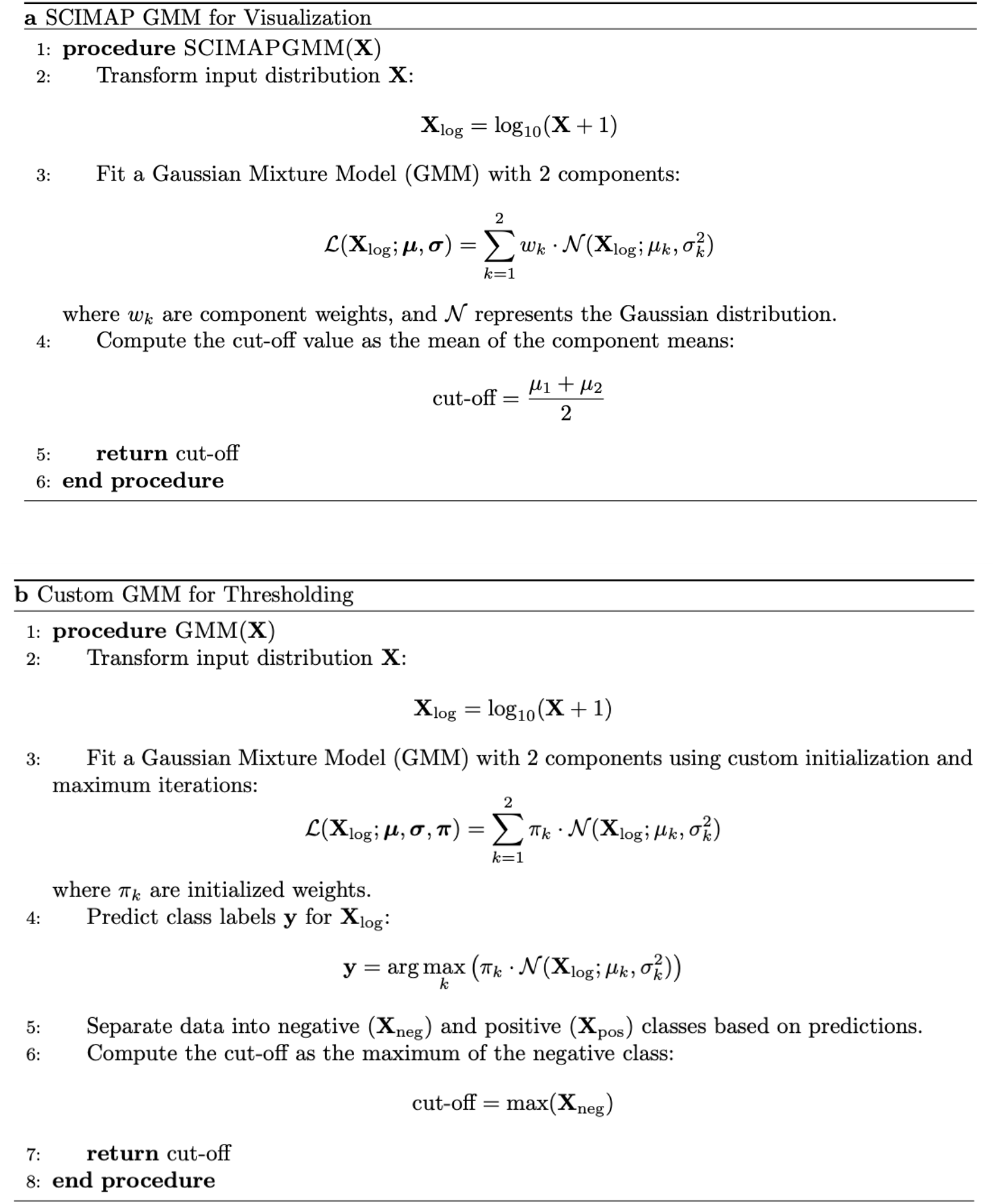

### Positive population change analysis

For each normalization method, we conducted a positive population change analysis using the GMM approach described above to quantify its efficacy. For each marker, a local positive population threshold is calculated for each raw sample, while a global threshold is determined for the combined normalized samples. Then we evaluated the proportions of cells above the local threshold and global threshold for each sample and the mean and standard deviation of changes in positive population proportions can be determined.

### kBET analysis

Due to the computational inefficiency of the R implementation of kBET, we performed kBET analysis using pegasuspy, a python package with kBET implementation, to measure batch effects before and after normalization. We followed the guidelines on the package’s website: https://pegasus.readthedocs.io/en/stable/ for implementation. Preprocessing included data standardization (Z-score) and dimensionality reduction using principal component analysis (PCA), followed by the construction of a neighborhood graph and a UMAP embedding to enhance the representation of local data structures. The kBET algorithm was applied to the UMAP representation to assess batch effects more effectively. The reported p-values and Chi-square statistics are three outputs from the Pegasus kBET pipeline. These metrics are not derived from independently fitted statistical models but are integral to the kBET framework.

kBET leveraged the neighborhood graph generated from the UMAP representation to evaluate how well samples from different batches mixed within their local neighborhoods. This analysis involved determining the k-nearest neighbors (kNN) for each cell and assessing whether the observed distribution of batch labels within these neighborhoods deviated from an ideal, uniform distribution. The deviation was quantified using a chi-squared test, resulting in a kBET score that reflects the extent of batch mixing. The kBET acceptance rate, calculated from the proportion of tests where the null hypothesis (no batch effect) was accepted, provided an overall measure of batch effect mitigation. A higher acceptance rate indicated more successful batch effect removal and a more homogeneous sample distribution across batches.

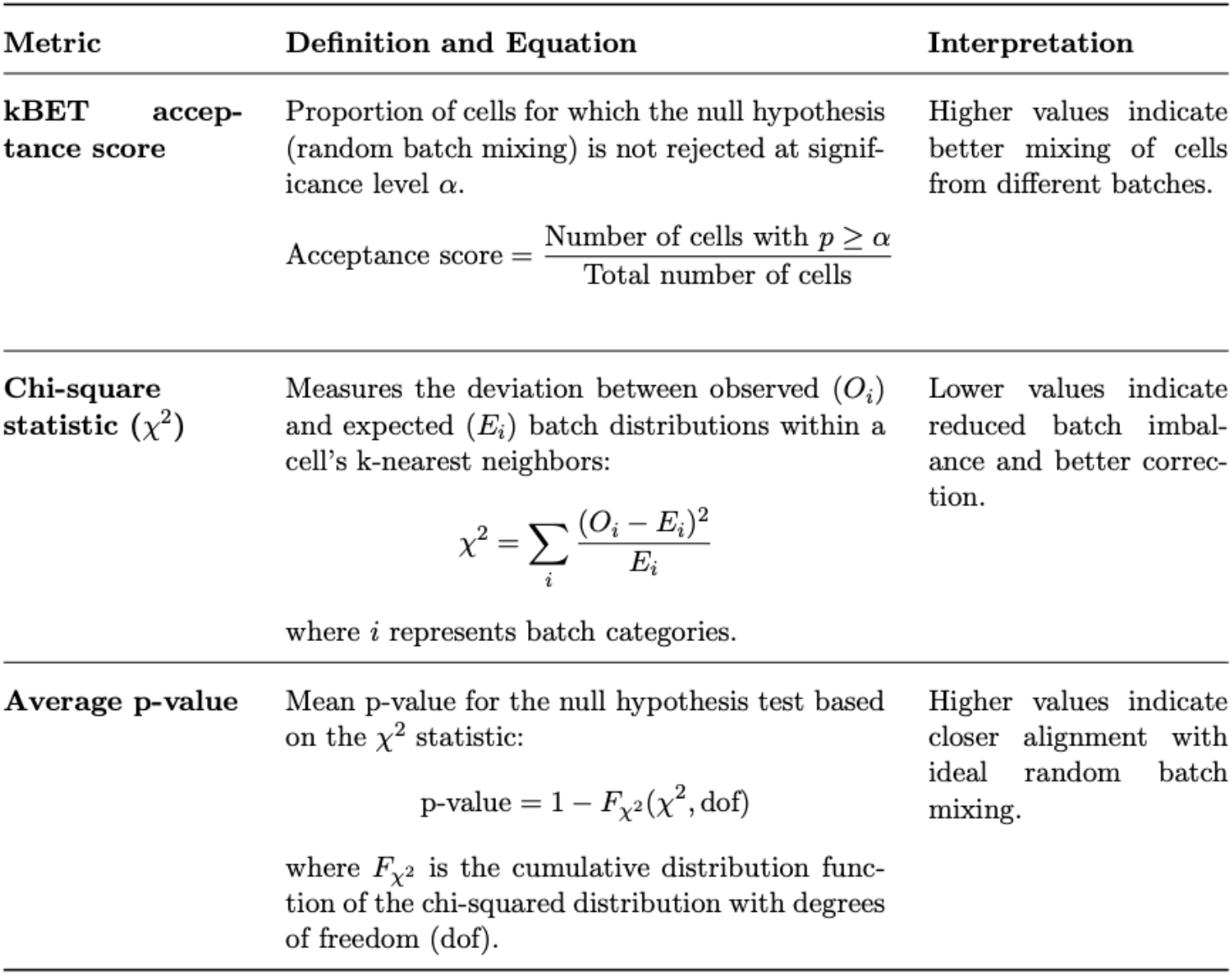

### Silhouette coefficient analysis

To evaluate how well normalization preserved biologically distinct cell populations, we performed Silhouette score analysis using the Silhouette score function from scikit-learn (https://scikit-learn.org/stable/modules/generated/sklearn.metrics.silhouette_score.html). Manual thresholding of mutually exclusive markers on the raw dataset—ECAD vs. CD45 and αSMA vs. ECAD—was used to define three broad and biologically meaningful cell types: tumor epithelial cells (ECAD^+^), immune cells (CD45^+^), and non-immune stromal cells (αSMA^+^). These manually annotated cell type labels served as ground truth cluster assignments for computing the Silhouette score, which quantifies the separation between these mutually exclusive populations. Prior to analysis, all datasets were standardized using Z-score scaling to ensure comparable marker expression ranges. A scale-invariant cosine distance metric was employed for Silhouette score calculation, enabling assessment of clustering quality based on the relative orientation of expression profiles rather than their absolute magnitude.

### UMAP visualization

For UMAP visualization, we utilized the pegasuspy package (https://pegasus.readthedocs.io/en/stable/index.html) to construct UMAP embeddings on top of the neighborhood graph. The UMAP was derived from the dataset after neighborhood graph construction to effectively capture the local and global structure of the data. To visualize the UMAP, we colored the plots by sample names and by CD45, αSMA, and ECAD marker intensity values.

### Leiden clustering analysis

We performed Leiden clustering analysis using Scanpy^33^ to identify distinct groups within the dataset before and after normalization. First data is transformed by Z-score scaling, A neighborhood graph was constructed using 50 nearest neighbors, followed by Leiden clustering at a resolution of 0.3. To evaluate sample representation in each cluster, we calculated proportions by dividing the count of cells per sample by the total cell count in each cluster.

### Data visualization

Matplotlib and seaborn packages in Python were used to generate all plots in this manuscript.

### Statistics and reproducibility

All analyses were conducted using publicly available MTI dataset CRC-ORION and our in-house PRAD-CyCIF and TMA-Lunaphore datasets, encompassing pixel-level and feature-level data across multiple batches. Data preprocessing involved log transformation with an offset to stabilize the variance. Benchmarking of UniFORM against Z-score, ComBat, and MxNorm was performed using kBET to evaluate batch effect mitigation and Gaussian Mixture Models (GMMs) to assess the preservation of positive population percentages. Pixel-level normalization was evaluated by aligning pixel intensity distributions with feature-level data, with Spearman correlations quantifying the consistency between levels.

For reproducibility, we provide the necessary dataset and code used in our analysis.

## Data availability

The CRC-ORION dataset is publicly available through the Human Tumor Atlas Network (HTAN) at https://humantumoratlas.org/, under participant IDs: HTA7_926, HTA7_927, HTA7_932, HTA7_934, HTA7_966, and HTA7_947. Our in-house PRAD-CyCIF dataset is accessible on Zenodo at https://zenodo.org/records/14257384.

## Code availability

All software used in the study is detailed in the Methods section and supplementary material. All scripts are available via GitHub: https://github.com/kunlunW/UniFORM

## Acknowledgments

This work was carried out with major support from the National Cancer Institute (NCI) Human Tumor Atlas Network (HTAN) Research Centers at OHSU (U2CCA233280), and funding (CEDAR3410918) from the Cancer Early Detection Advanced Research Centre at Oregon Health & Science University, Knight Cancer Institute (S.E.E.). Y.H.C. is supported by R01 CA253860, R01CA276224, U01 294548, Kuni Foundation Imagination Grants, and the Cancer Early Detection Advanced Research Center (CEDAR) pilot project grant. The resources of the Exacloud high-performance computing environment developed jointly by OHSU and Intel and the technical support of the OHSU Advanced Computing Center are gratefully acknowledged.

## Author Contributions

Conceptualization: K.W., Y.H.C. Software implementation: K.W., Y.H.C. Data analysis: K.W., S.K., K.A. Quality check and code review: K.W., Y.H.C., S.E.E., Z.S., E.C. Data generation: Z.S., J.Y. Manuscript writing: K.W., Y.H.C., S.E.E. Funding: M.W., G.B.M., S.E.E., Y.H.C. All authors contributed to the paper either through discussions or directly.

## Competing interests

The authors declare the following competing interests: G.B.M. is a SAB member or Consultant: for Amphista, Astex, AstraZeneca, BlueDot, Chrysallis Biotechnology, Ellipses Pharma, GSK, ImmunoMET, Infinity, Ionis, Leapfrog Bio, Lilly, Medacorp, Nanostring, Nuvectis, PDX Pharmaceuticals, Qureator, Roche, Signalchem Lifesciences, Tarveda, Turbine, Zentalis Pharmaceuticals. G.B.M. has Stock/Options/Financial relationships with: Bluedot, Catena Pharmaceuticals, ImmunoMet, Nuvectis, SignalChem, Tarveda, and Turbine. G.B.M. has Licensed Technology: HRD assay to Myriad Genetics, DSP patents with Nanostring. G.B.M. has Sponsored research with AstraZeneca. The other authors declare no competing interests.

## Supplementary material

See separate supplementary material PDF

## Notes

### Summary of Updates

updated context and added more analyses

