## Supplemental information for "UniFORM: Towards Universal ImmunoFluorescence Normalization for Multiplex Tissue Imaging"

| Specimen ID | HTAN Participant ID | Cohort |
|---|---|---|
| CRC01 | HTA7_926 | 1 |
| CRC02 | HTA7_927 | 1 |
| CRC03 | HTA7_932 | 1 |
| CRC04 | HTA7_934 | 1 |
| CRC05 | HTA7_966 | 1 |
| CRC06 | HTA7_947 | 1 |

| Marker | Channel | Function |
|---|---|---|
| Hoechst | 1 | Nuclear/DNA stain |
| AF1 | 2 | Autofluorescence control |
| CD31 | 3 | Endothelial marker; cell adhesion molecule |
| CD45 | 4 | General lymphocyte marker |
| CD68 | 5 | Macrophage (M1) |
| Argo550 | 6 | Autofluorescence control |
| CD4 | 7 | T helper cells |
| FOXP3 | 8 | Regulatory T cells (Tregs) |
| CD8a | 9 | Cytotoxic T cells |
| CD45RO | 10 | Memory T-cell marker |
| CD20 | 11 | B-cell marker |
| PD-L1 | 12 | Immune checkpoint ligand |
| CD3e | 13 | T-cell receptor complex |
| CD163 | 14 | Macrophage (M2) |
| E-cadherin | 15 | Epithelial marker |
| PD-1 | 16 | Immune checkpoint receptor |
| Ki67 | 17 | Proliferation marker |
| Pan-CK | 18 | Epithelial/tumor marker |
| SMA | 19 | Smooth muscle actin (fibroblast/muscle marker) |

**Table S1a CRC-ORION metadata information.**

| Sample Name | Batch ID |
|---|---|
| PRAD-01 | batch4 |
| PRAD-02 | batch1 |
| PRAD-03 | batch1 |
| PRAD-04 | batch4 |
| PRAD-05 | batch1 |
| PRAD-06 | batch4 |
| PRAD-07 | batch2 |
| PRAD-08 | batch2 |
| PRAD-09 | batch3 |
| PRAD-10 | batch3 |
| PRAD-11 | batch5 |
| PRAD-12 | batch2 |
| PRAD-13 | batch7 |
| PRAD-14 | batch7 |
| PRAD-15 | batch6 |
| PRAD-16 | batch6 |
| PRAD-17 | batch6 |
| PRAD-18 | batch6 |
| PRAD-19 | batch6 |
| PRAD-20 | batch3 |



| Marker | Channel | Function |
|---|---|---|
| DAPI | 1 | DNA/nuclear stain |
| EPCAM | 2 | Epithelial/tumor marker |
| CD56 | 3 | Natural killer (NK) cell marker |
| CD45 | 4 | General lymphocyte marker |
| aSMA | 5 | Smooth muscle actin (fibroblast/muscle marker) |
| ChromA | 6 | Chromogranin A (neuroendocrine marker) |
| CK14 | 7 | Epithelial cell marker |
| Ki67 | 8 | Proliferation marker |
| GZMB | 9 | T cell activation; cytotoxic granules |
| PD1 | 10 | Immune checkpoint receptor |
| ECAD | 11 | Epithelial marker (E-cadherin) |
| CD31 | 12 | Endothelial marker; cell adhesion molecule |
| CD45RA | 13 | Naive T-cell marker |
| HLADRB1 | 14 | MHC Class II antigen |
| CD3 | 15 | T-cell receptor complex |
| p53 | 16 | Tumor suppressor protein |
| CDX2 | 17 | GI origin carcinoma marker |
| CD20 | 18 | B-cell marker |
| NOTCH1 | 19 | Regulates cell fate decisions |

**Table S1b CRC-ORION metadata information.**

| Sample Name | Batch/Replicate (TMA) |
|---|---|
| Breast Cancer | TMA4<br>TMA5<br>TMA6 |
| Breast MCF7 | TMA4<br>TMA5<br>TMA6 |
| Colon Ad | TMA4<br>TMA5<br>TMA6 |
| Colon HCT116 | TMA4<br>TMA5<br>TMA6 |
| Glioma | TMA4<br>TMA5<br>TMA6 |
| Liver Ad | TMA4<br>TMA5<br>TMA6 |
| Normal Tonsil | TMA4<br>TMA5<br>TMA6 |
| Pancreas Panc-1 | TMA4<br>TMA5<br>TMA6 |
| Prostate 22RV1 | TMA4<br>TMA5<br>TMA6 |
| Prostate LNCaP | TMA4<br>TMA5<br>TMA6 |
| Prostate RWPE-1 | TMA4<br>TMA5<br>TMA6 |
| Prostate PC3 | TMA4<br>TMA5<br>TMA6 |

| Marker | Channel | Function |
|---|---|---|
| DAPI | 1 | Nuclear stain |
| TRITC (ms555) | 2 | Fluorescent label |
| Cy5 (rb647) | 3 | Fluorescent label |
| TOMM20 | 4 | Mitochondrial marker |
| CD90 | 5 | Fibroblast/stem cell marker |
| CD45 | 6 | Leukocyte marker |
| ERG | 7 | Endothelial/prostate marker |
| HLA-DR | 8 | MHC class II antigen |
| CD11b | 9 | Myeloid lineage marker |
| CD3 | 10 | T cell marker |
| AR | 11 | Androgen receptor |
| TUBB3 | 12 | Neuronal marker |
| GZMB | 13 | Cytotoxic granule |
| E-cad | 14 | Cell adhesion (epithelial) |
| CK5 | 15 | Basal epithelial marker |
| CD68 | 16 | Macrophage marker |
| TH | 17 | Dopaminergic marker |
| aSMA | 18 | Smooth muscle/fibroblast marker |
| AMACR | 19 | Prostate cancer marker |
| CD56 | 20 | NK cell marker |
| NFKB | 21 | Inflammatory signaling |
| HIF-1 | 22 | Hypoxia response |
| CD4 | 23 | Helper T cell marker |
| FOXA1 | 24 | Luminal transcription factor |
| ADAM10 | 25 | Metalloproteinase |
| DCX | 26 | Neuronal migration marker |
| CD11c | 27 | Dendritic cell marker |
| CD20 | 28 | B cell marker |
| Ki67 | 29 | Proliferation marker |
| CD8 | 30 | Cytotoxic T cell marker |
| CD31 | 31 | Endothelial marker |
| VIM | 32 | Mesenchymal marker |
| NRXN1 | 33 | Synaptic adhesion |
| NLGN4X | 34 | Synaptic adhesion |
| ChromA | 35 | Neuroendocrine marker |
| TRYP | 36 | Melanocyte marker |
| CD44 | 37 | Adhesion/stemness marker |
| NLGN1 | 38 | Synaptic protein |
| CK8 | 39 | Luminal epithelial marker |
| B-catenin | 40 | Wnt signaling/adhesion |
| H3K4 | 41 | Histone modification |
| H3K27ac | 42 | Enhancer activation |
| CD163 | 43 | M2 macrophage marker |

**Table S1c TMA-Lunaphore metadata information.**

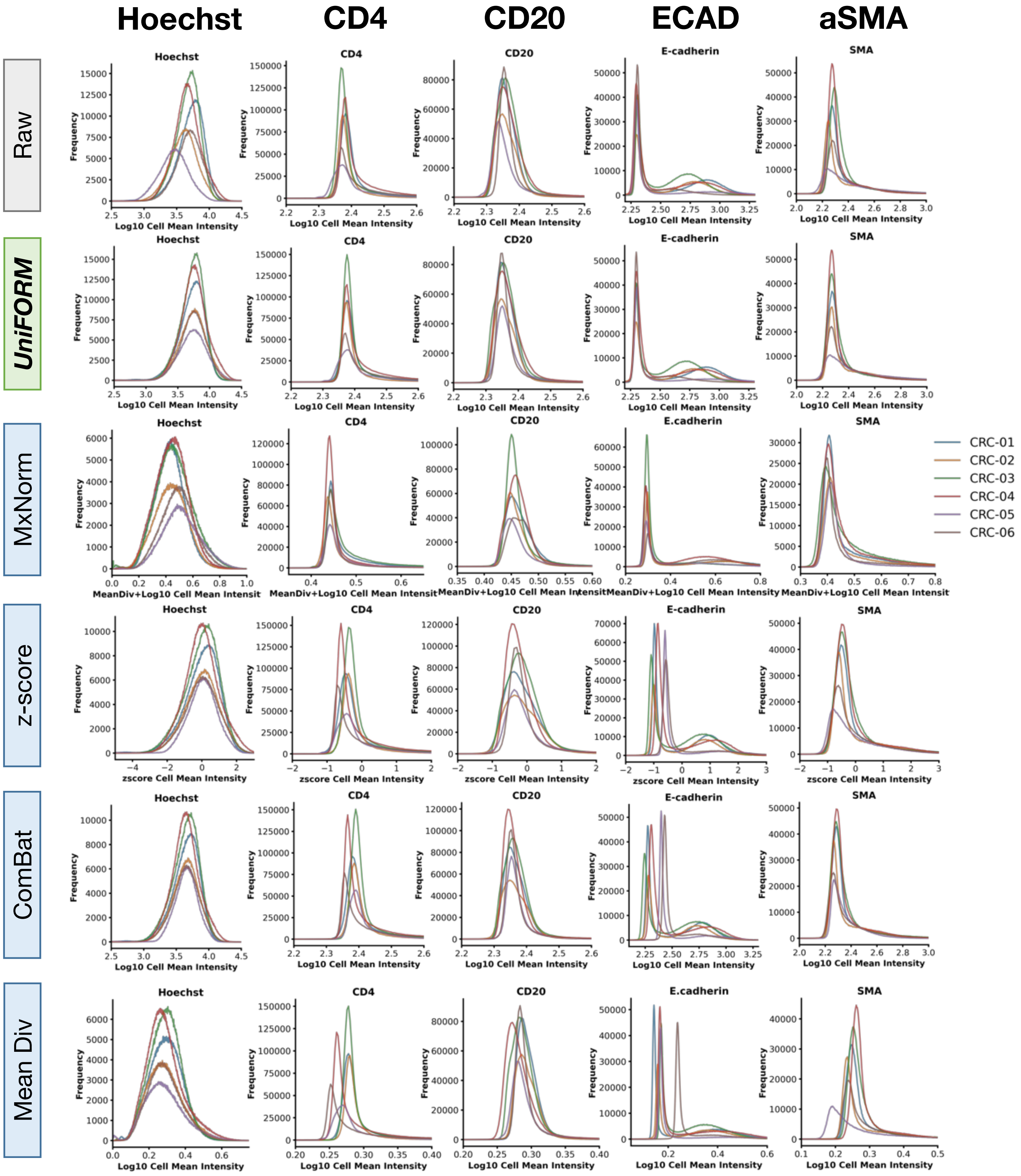


**Figure S1a CRC-ORION feature-level normalization across methods.** Markers such as CD45RO, PD-L1, and PD-1 exhibit substantial technical variation in cell intensity distributions across samples, despite all originating from the same cohort and batch. UniFORM effectively harmonizes inter-batch variation while preserving distribution shape and positive population counts. In contrast, MxNorm aligns simpler distributions but at the cost of distorting their distributions (e.g., aSMA). z-score, ComBat, and mean division all fail to properly align many markers and significantly distort distribution shapes.

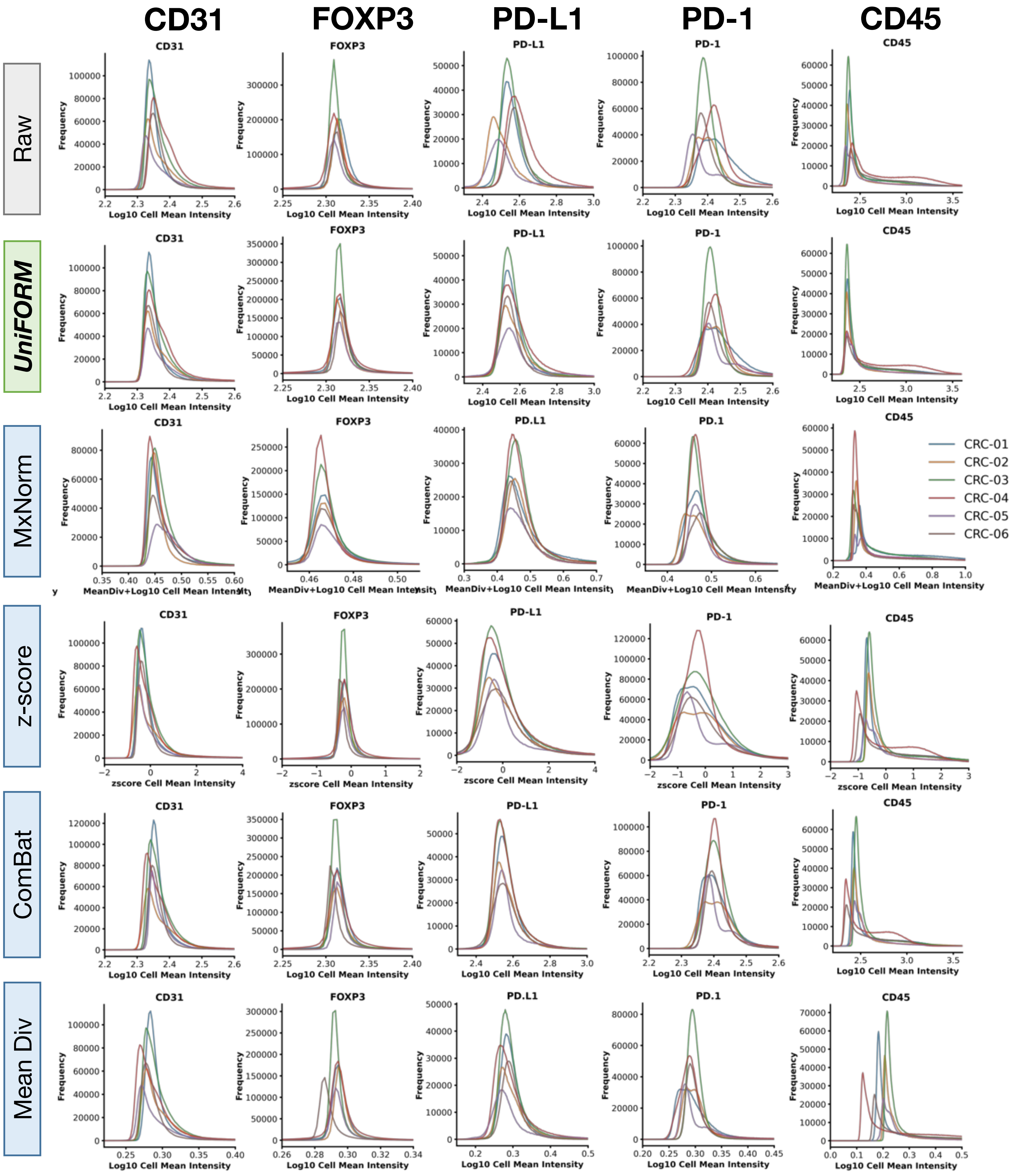


**Figure S1b CRC-ORION feature-level normalization across methods.** Markers such as CD45RO, PD-L1, and PD-1 exhibit substantial technical variation in cell intensity distributions across samples, despite all originating from the same cohort and batch. UniFORM effectively harmonizes inter-batch variation while preserving distribution shape and positive population counts. In contrast, MxNorm aligns simpler distributions but at the cost of distorting their distributions (e.g., aSMA). z-score, ComBat, and mean division all fail to properly align many markers and significantly distort distribution shapes.

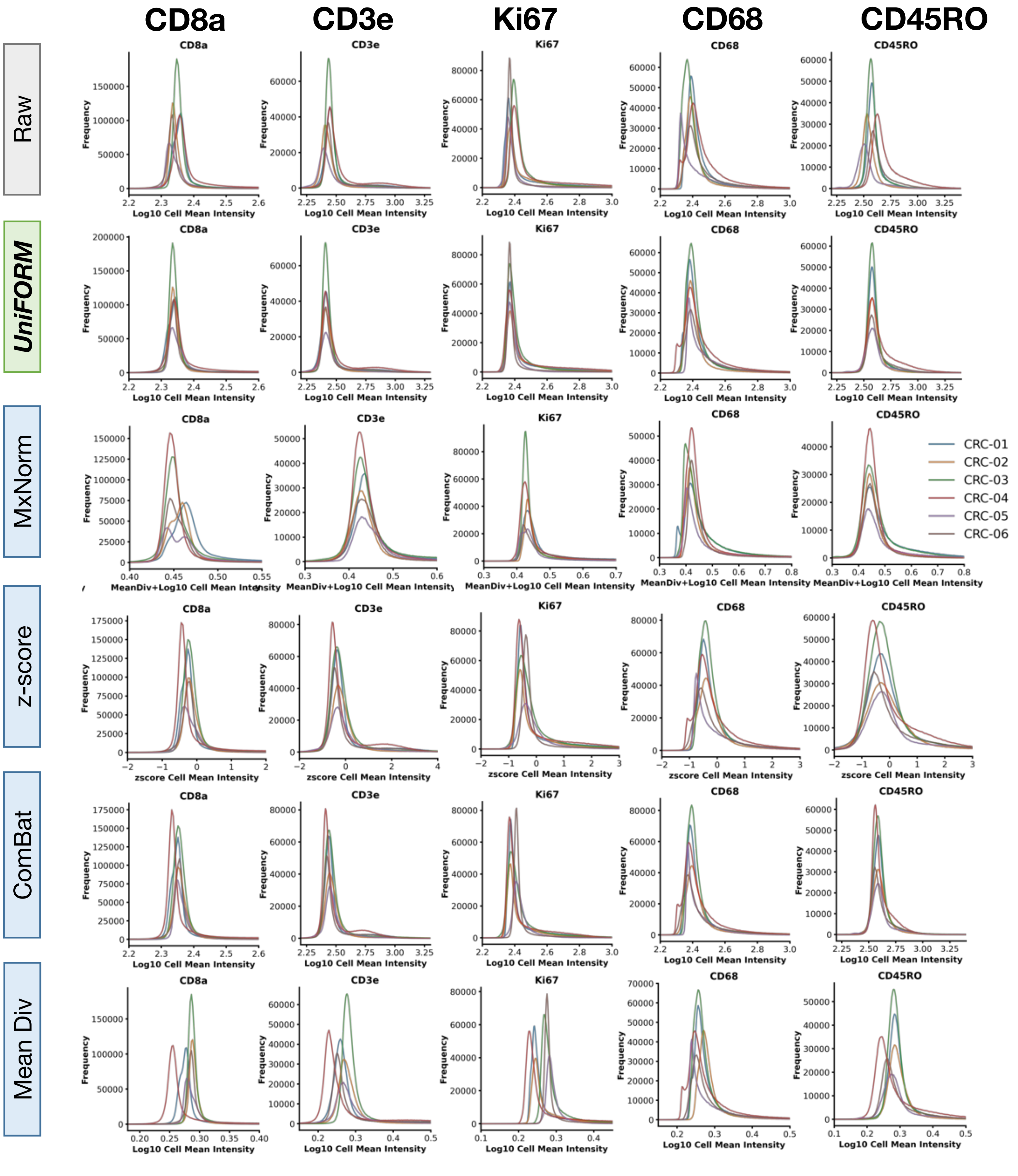


**Figure S1c CRC-ORION feature-level normalization across methods.** Markers such as CD45RO, PD-L1, and PD-1 exhibit substantial technical variation in cell intensity distributions across samples, despite all originating from the same cohort and batch. UniFORM effectively harmonizes inter-batch variation while preserving distribution shape and positive population counts. In contrast, MxNorm aligns simpler distributions but at the cost of distorting their distributions (e.g., aSMA). z-score, ComBat, and mean division all fail to properly align many markers and significantly distort distribution shapes.

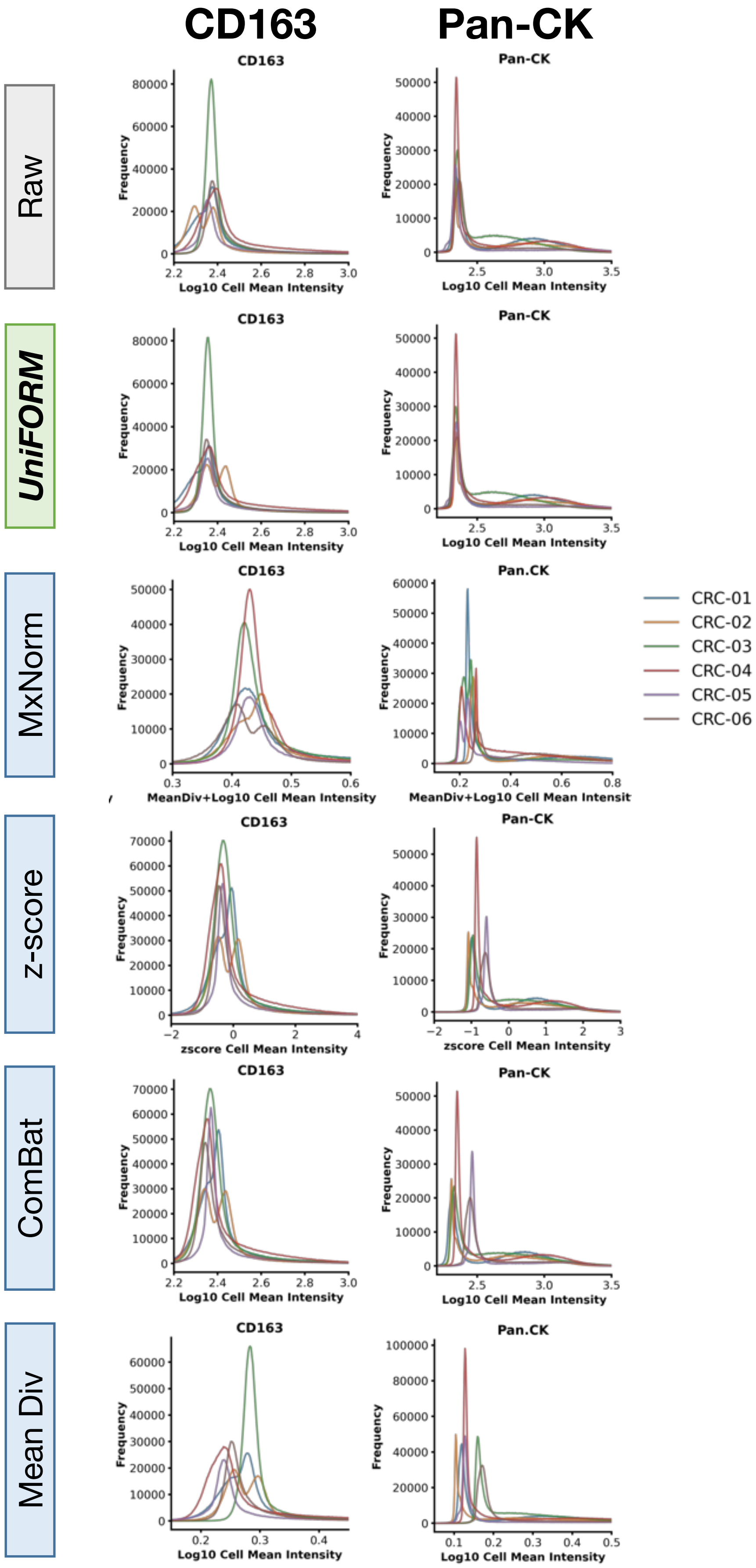


**Figure S1d CRC-ORION feature-level normalization across methods.** Markers such as CD45RO, PD-L1, and PD-1 exhibit substantial technical variation in cell intensity distributions across samples, despite all originating from the same cohort and batch. UniFORM effectively harmonizes inter-batch variation while preserving distribution shape and positive population counts. In contrast, MxNorm aligns simpler distributions but at the cost of distorting their distributions (e.g., aSMA). z-score, ComBat, and mean division all fail to properly align many markers and significantly distort distribution shapes.

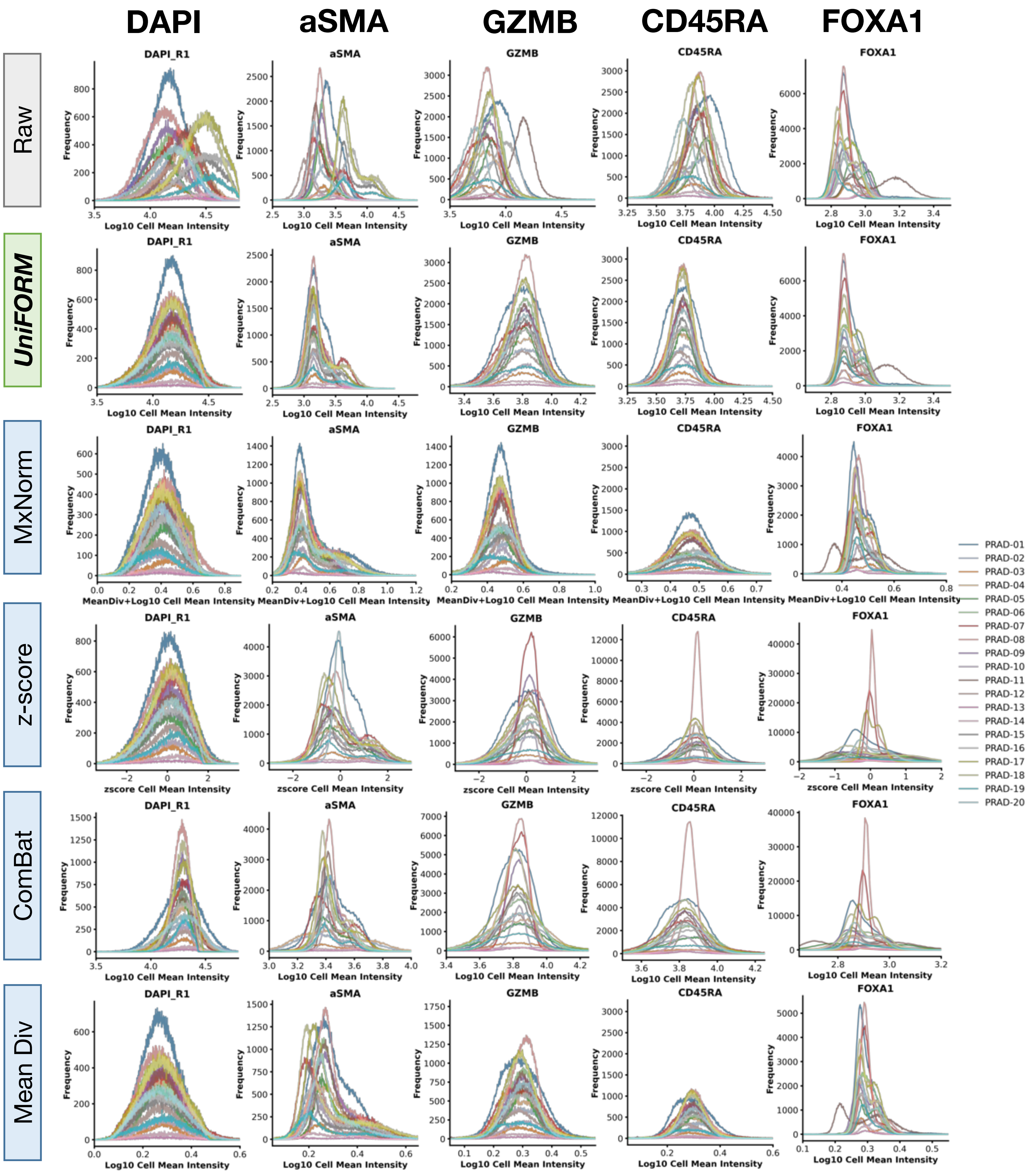


**Figure S2a. PRAD-CyCIF feature-level normalization across methods.** Raw data show substantial batch effects and heterogeneous marker distributions. UniFORM accurately aligns distributions while preserving shape and positive population counts. In contrast, MxNorm, z-score, ComBat, and mean division all fail to align most markers and significantly distort distribution shapes, leading to potential loss of biological information—especially for complex markers like ECAD.

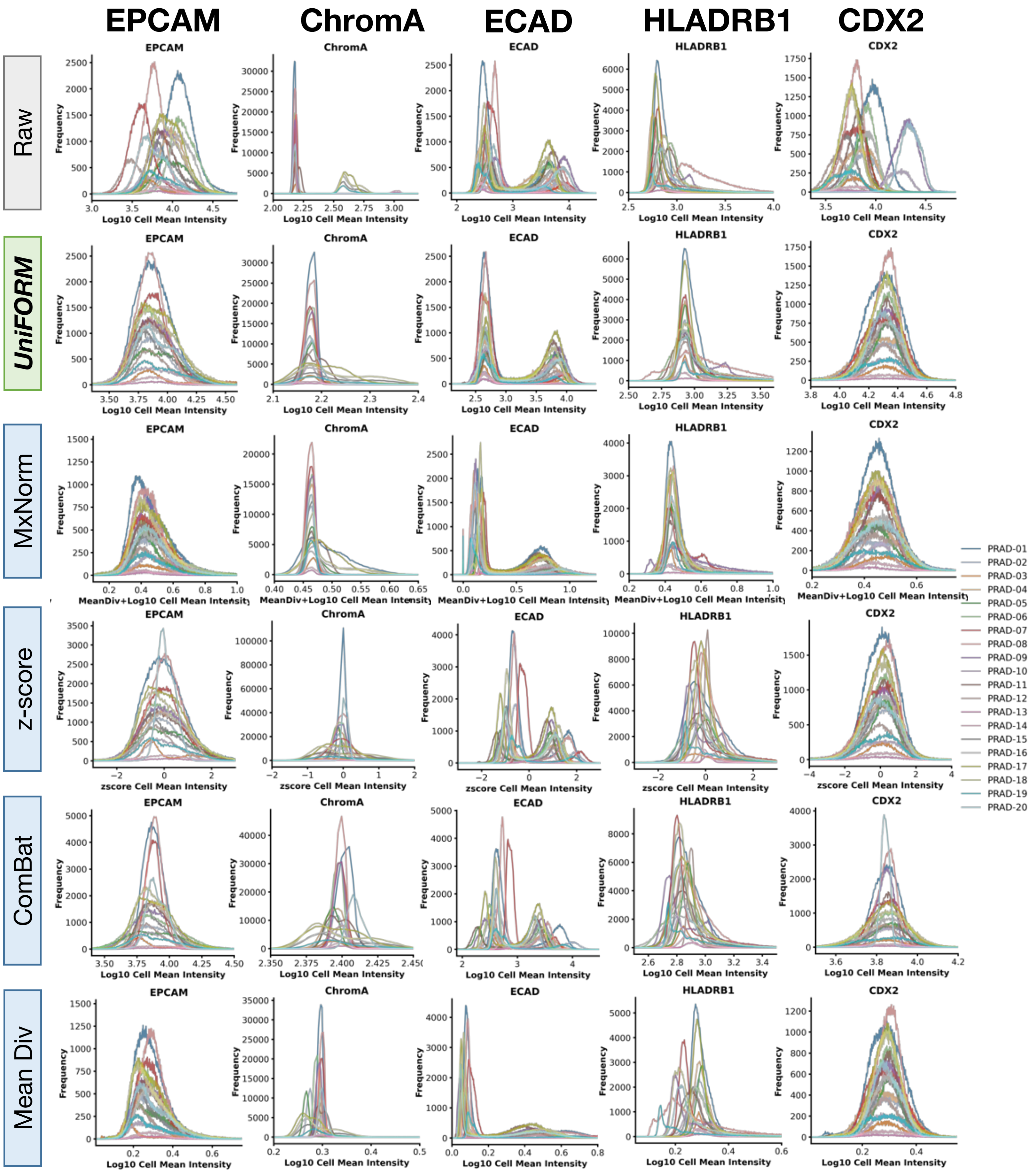


**Figure S2b. PRAD-CyCIF feature-level normalization across methods.** Raw data show substantial batch effects and heterogeneous marker distributions. UniFORM accurately aligns distributions while preserving shape and positive population counts. In contrast, MxNorm, z-score, ComBat, and mean division all fail to align most markers and significantly distort distribution shapes, leading to potential loss of biological information—especially for complex markers like ECAD.

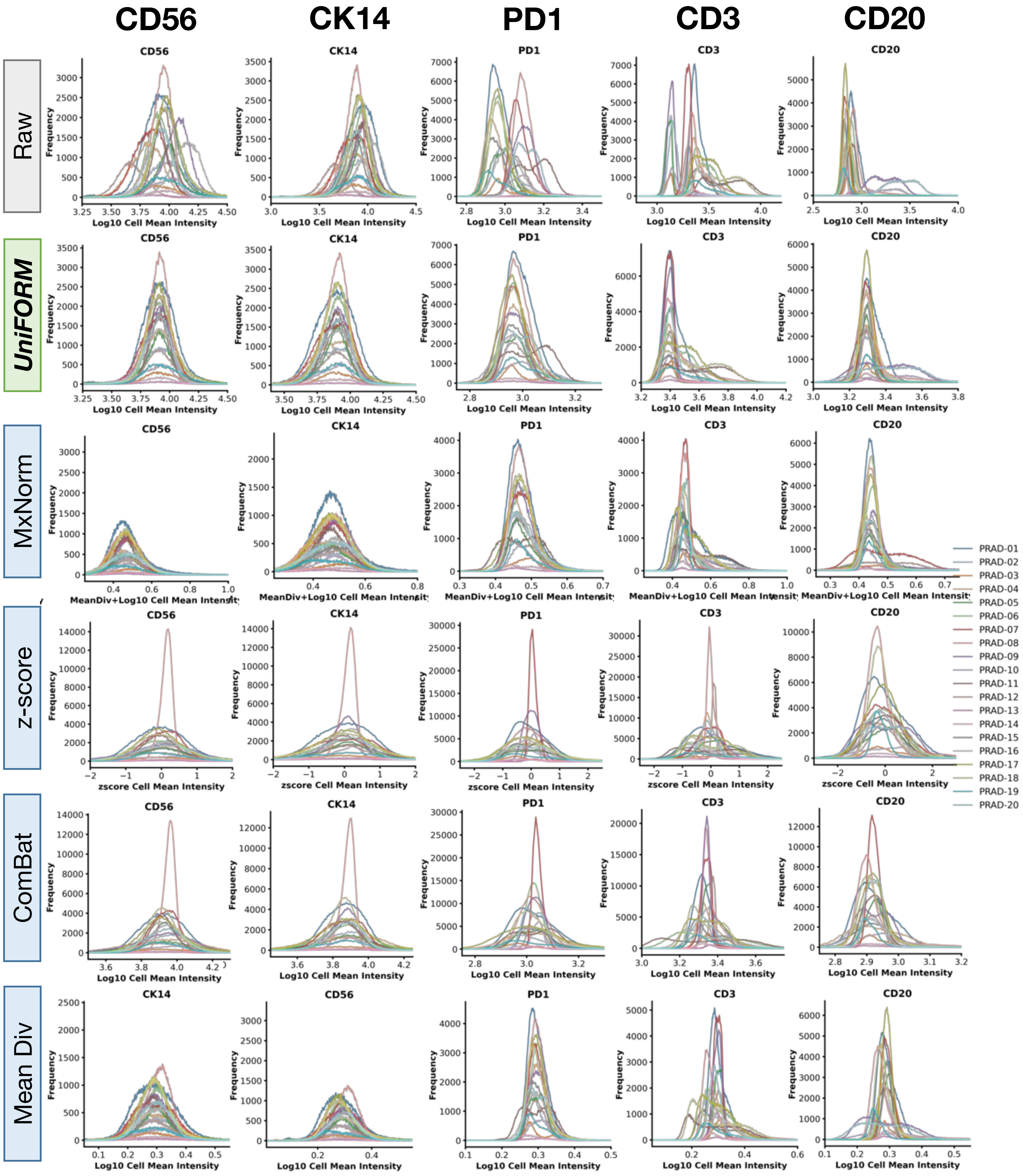


**Figure S2c. PRAD-CyCIF feature-level normalization across methods.** Raw data show substantial batch effects and heterogeneous marker distributions. UniFORM accurately aligns distributions while preserving shape and positive population counts. In contrast, MxNorm, z-score, ComBat, and mean division all fail to align most markers and significantly distort distribution shapes, leading to potential loss of biological information—especially for complex markers like ECAD.

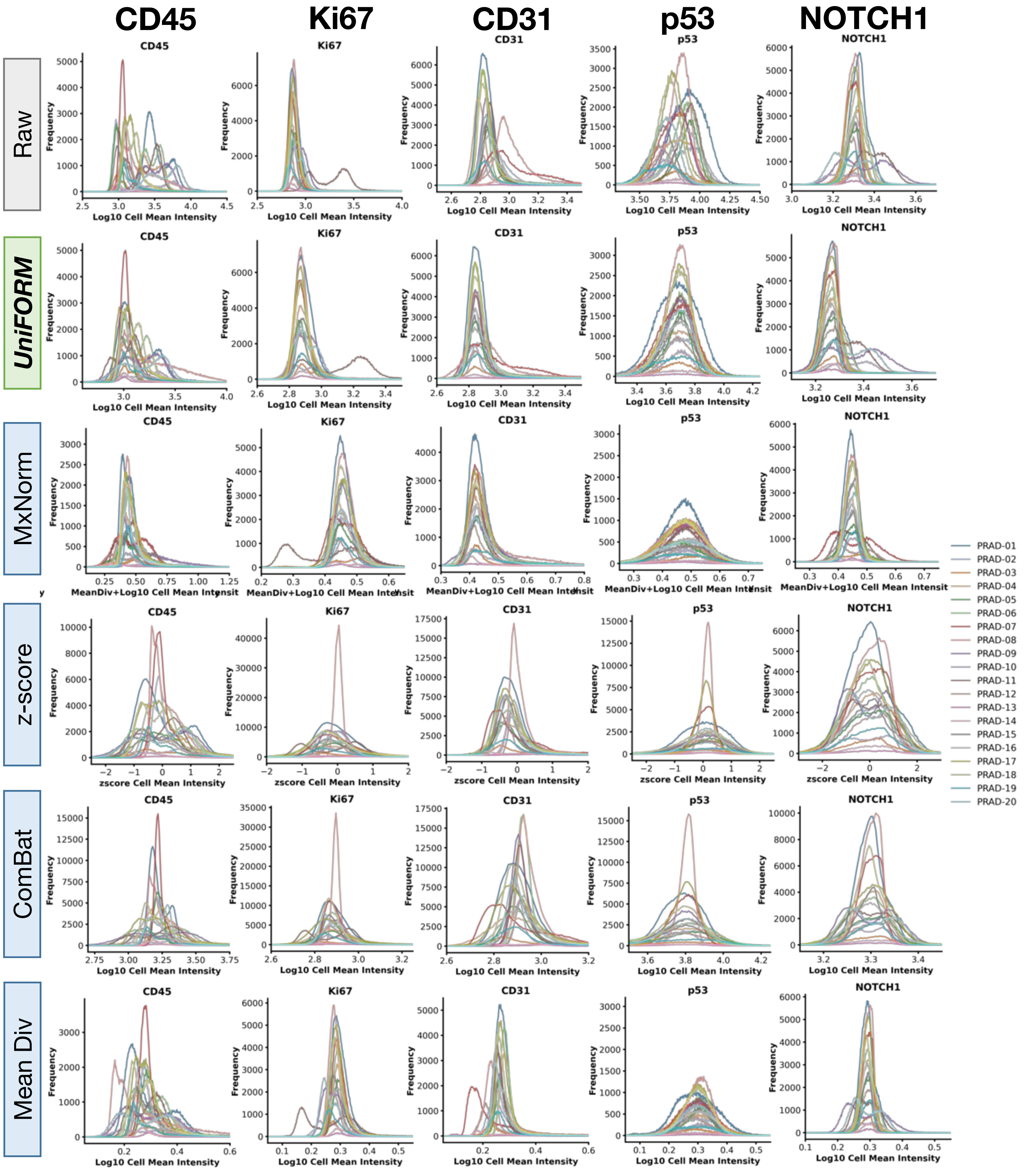


**Figure S2d. PRAD-CyCIF feature-level normalization across methods.** Raw data show substantial batch effects and heterogeneous marker distributions. UniFORM accurately aligns distributions while preserving shape and positive population counts. In contrast, MxNorm, z-score, ComBat, and mean division all fail to align most markers and significantly distort distribution shapes, leading to potential loss of biological information—especially for complex markers like ECAD.

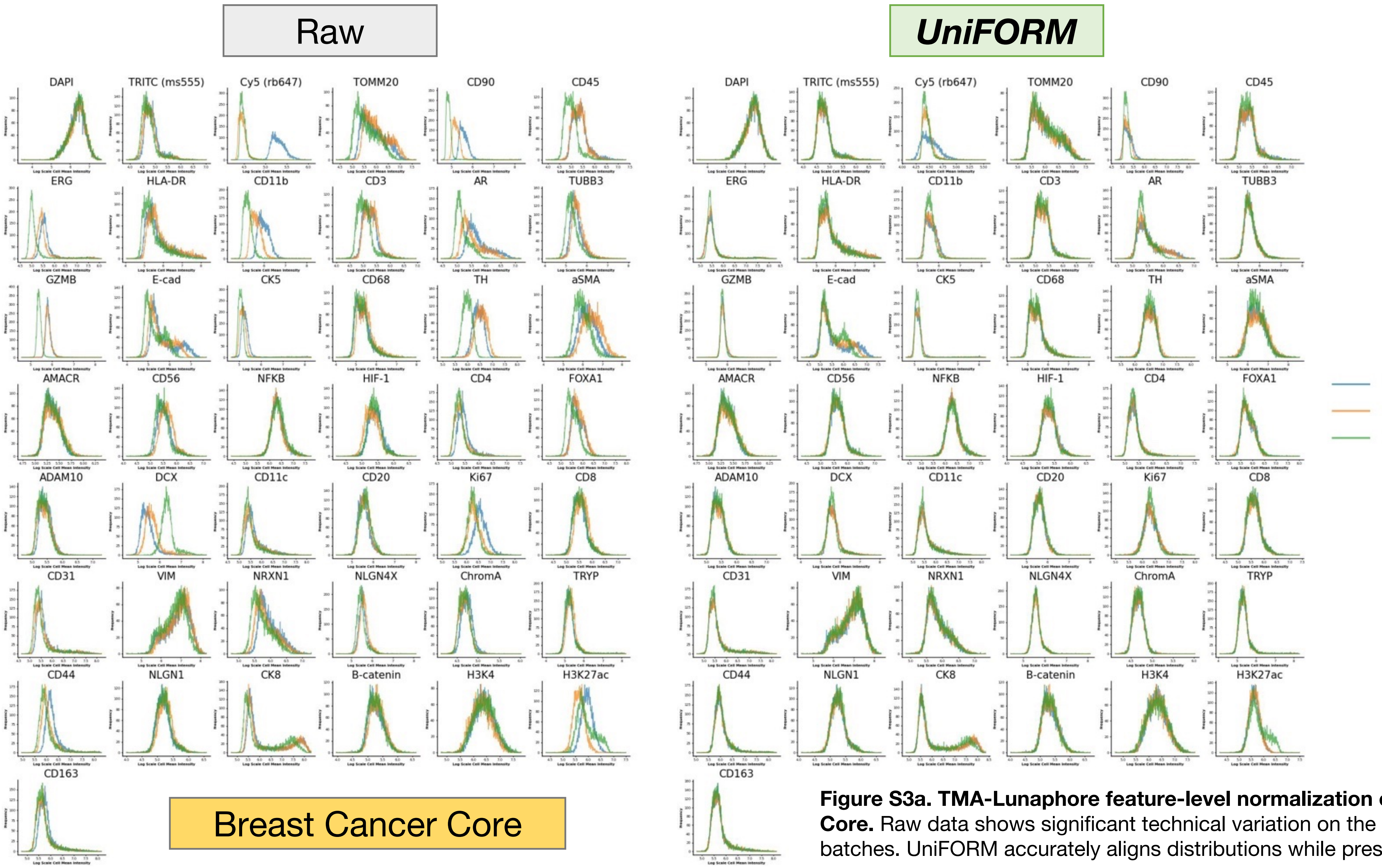


**Figure S3a. TMA-Lunaphore feature-level normalization on Breast Cancer Core.** Raw data shows significant technical variation on the same slide across batches. UniFORM accurately aligns distributions while preserving biology.

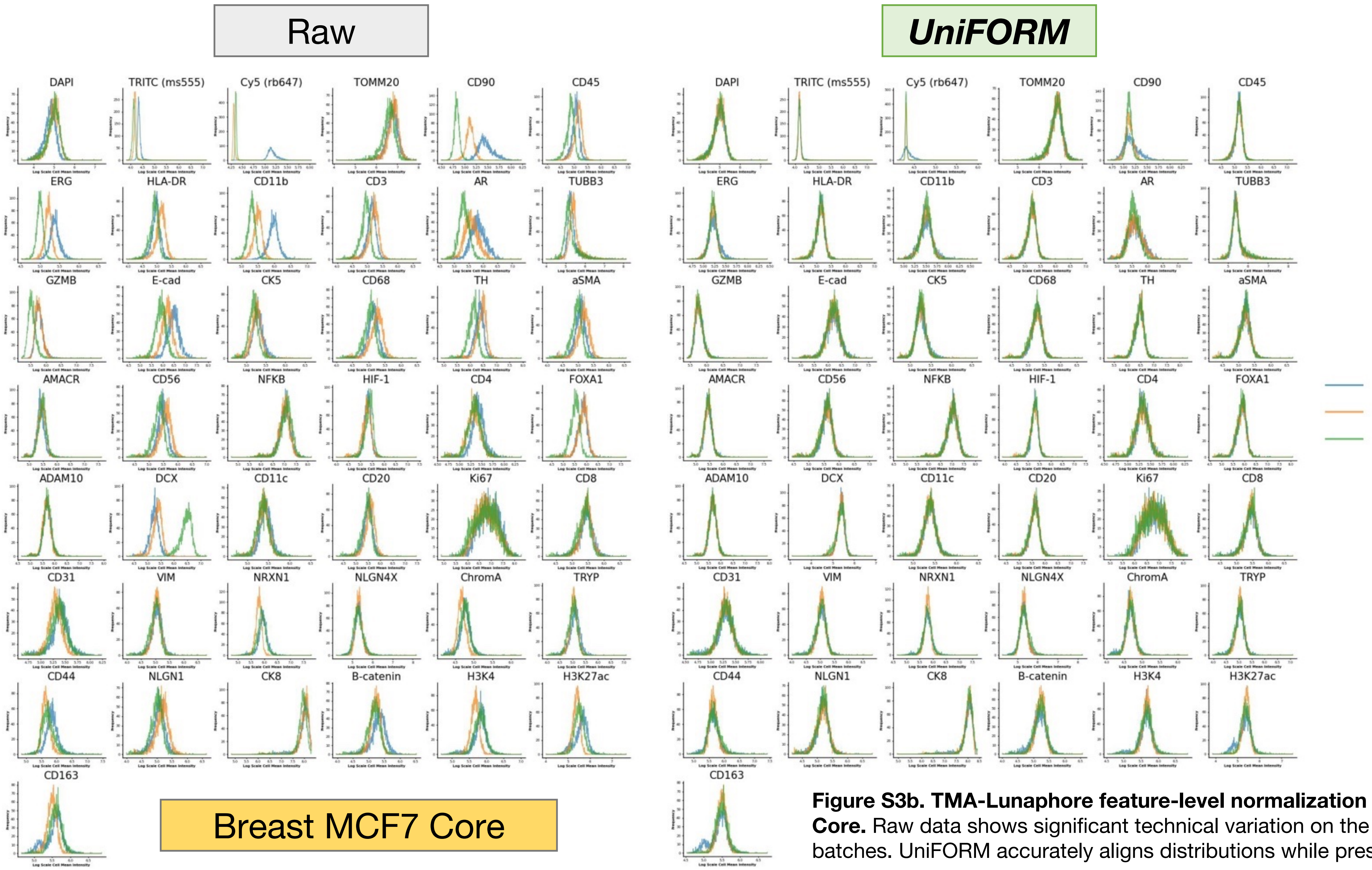


**Figure S3b. TMA-Lunaphore feature-level normalization on Breast MCF7 Core.** Raw data shows significant technical variation on the same slide across batches. UniFORM accurately aligns distributions while preserving biology.

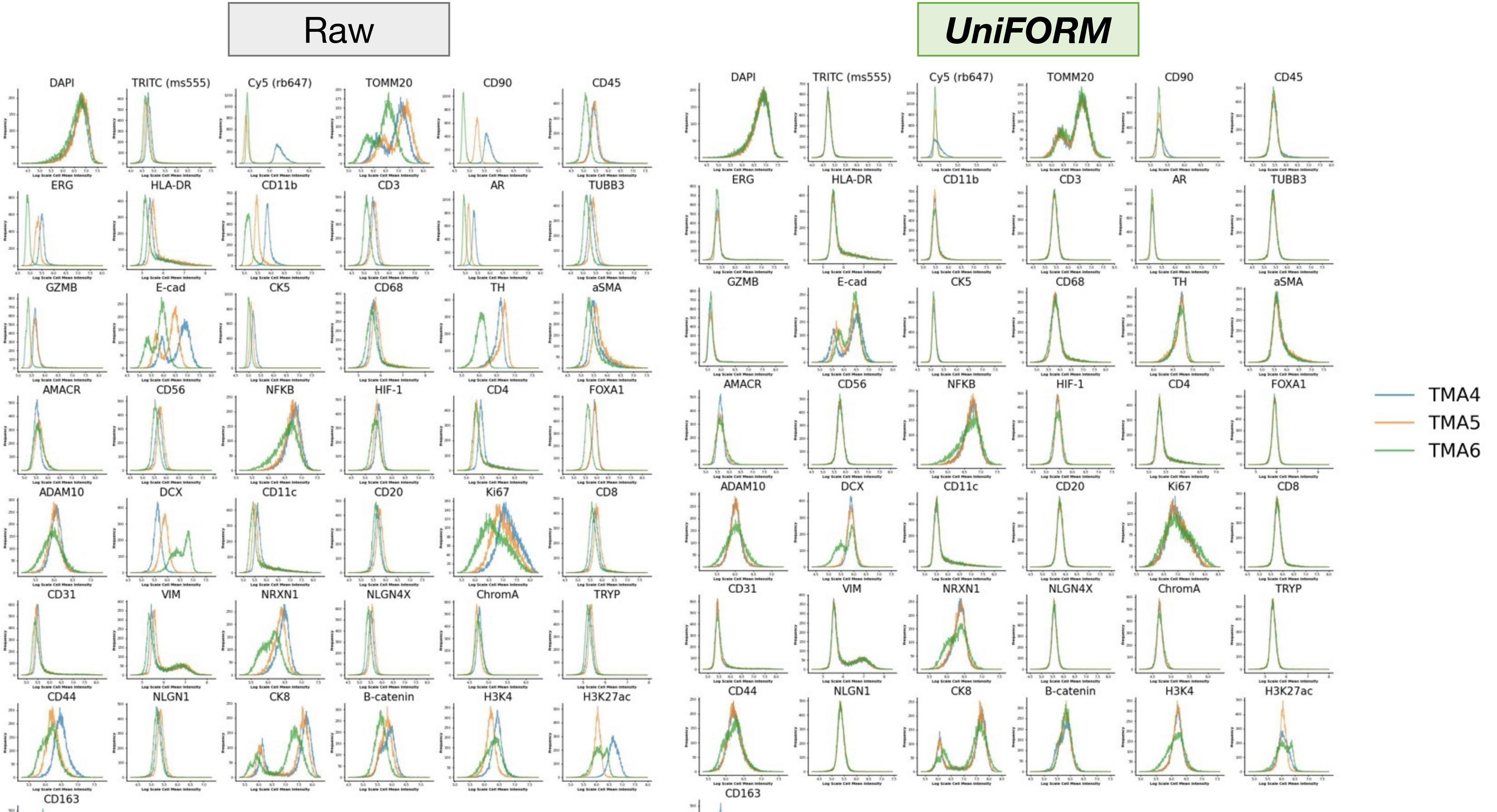

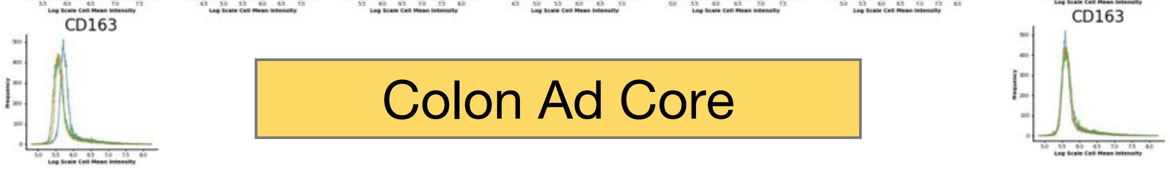


**Figure S3c. TMA-Lunaphore feature-level normalization on Colon Ad Core.** Raw data shows significant technical variation on the same slide across batches. UniFORM accurately aligns distributions while preserving biology.

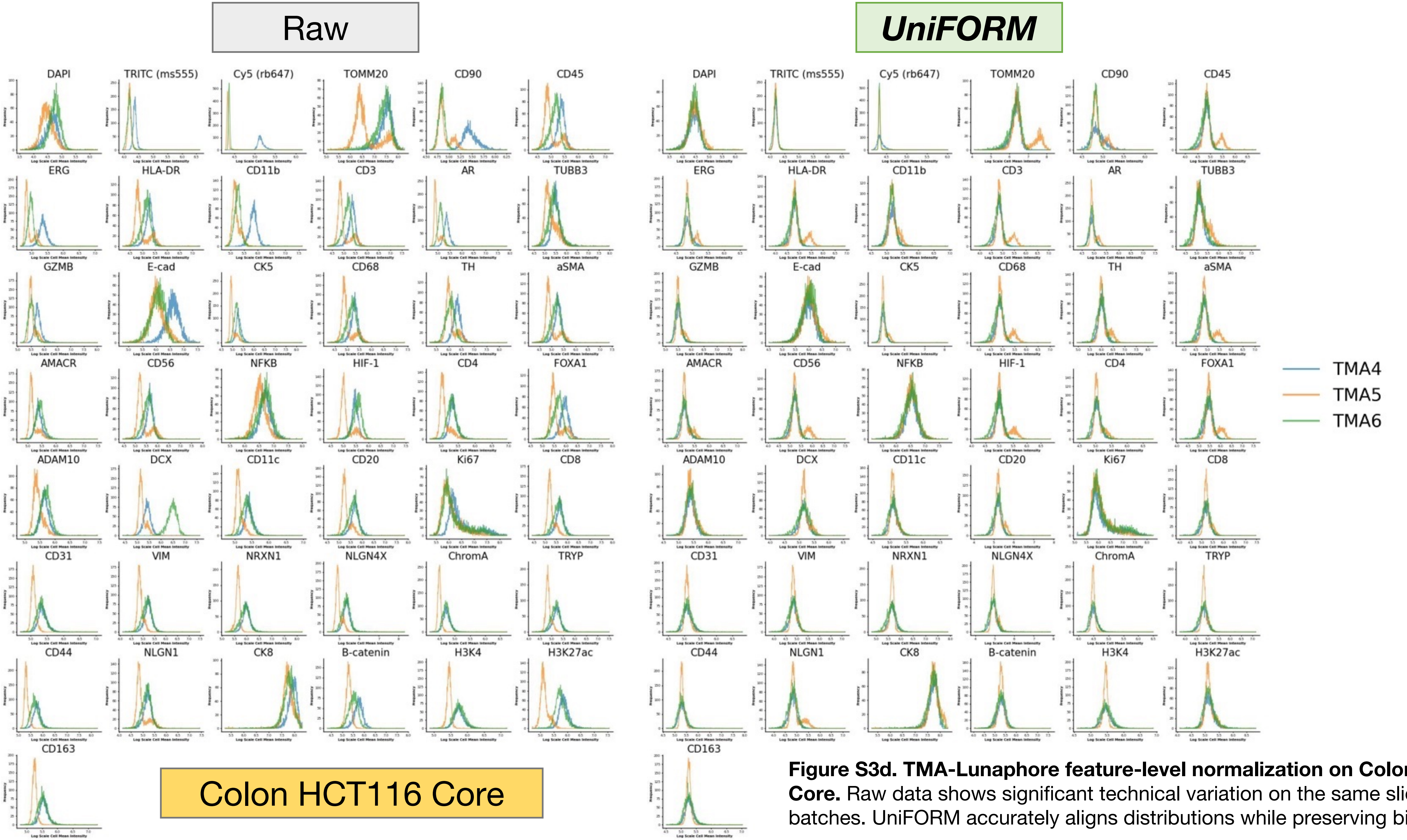


**Figure S3d. TMA-Lunaphore feature-level normalization on Colon HCT116 Core.** Raw data shows significant technical variation on the same slide across batches. UniFORM accurately aligns distributions while preserving biology.

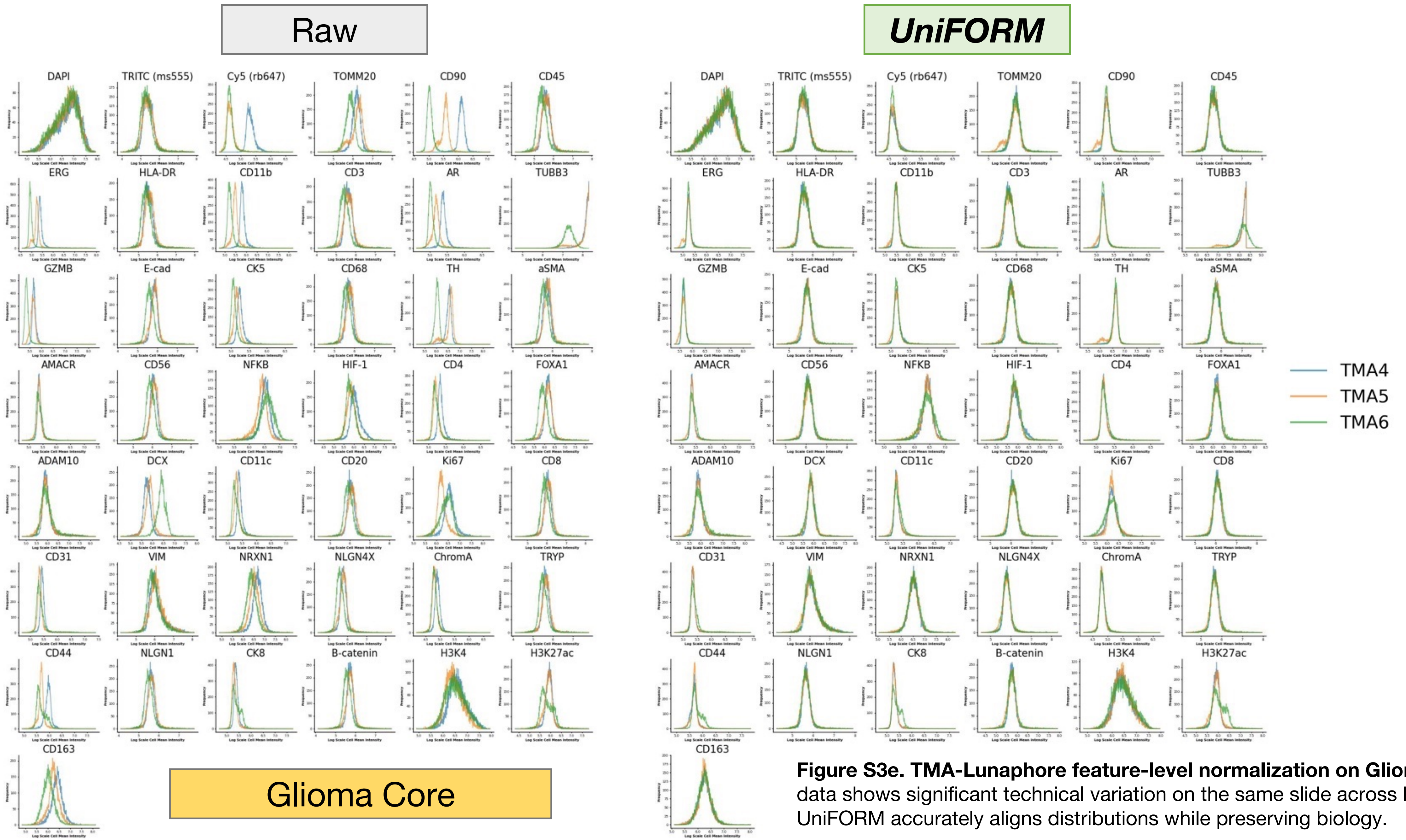


**Figure S3e. TMA-Lunaphore feature-level normalization on Glioma Core.** Raw data shows significant technical variation on the same slide across batches. UniFORM accurately aligns distributions while preserving biology.

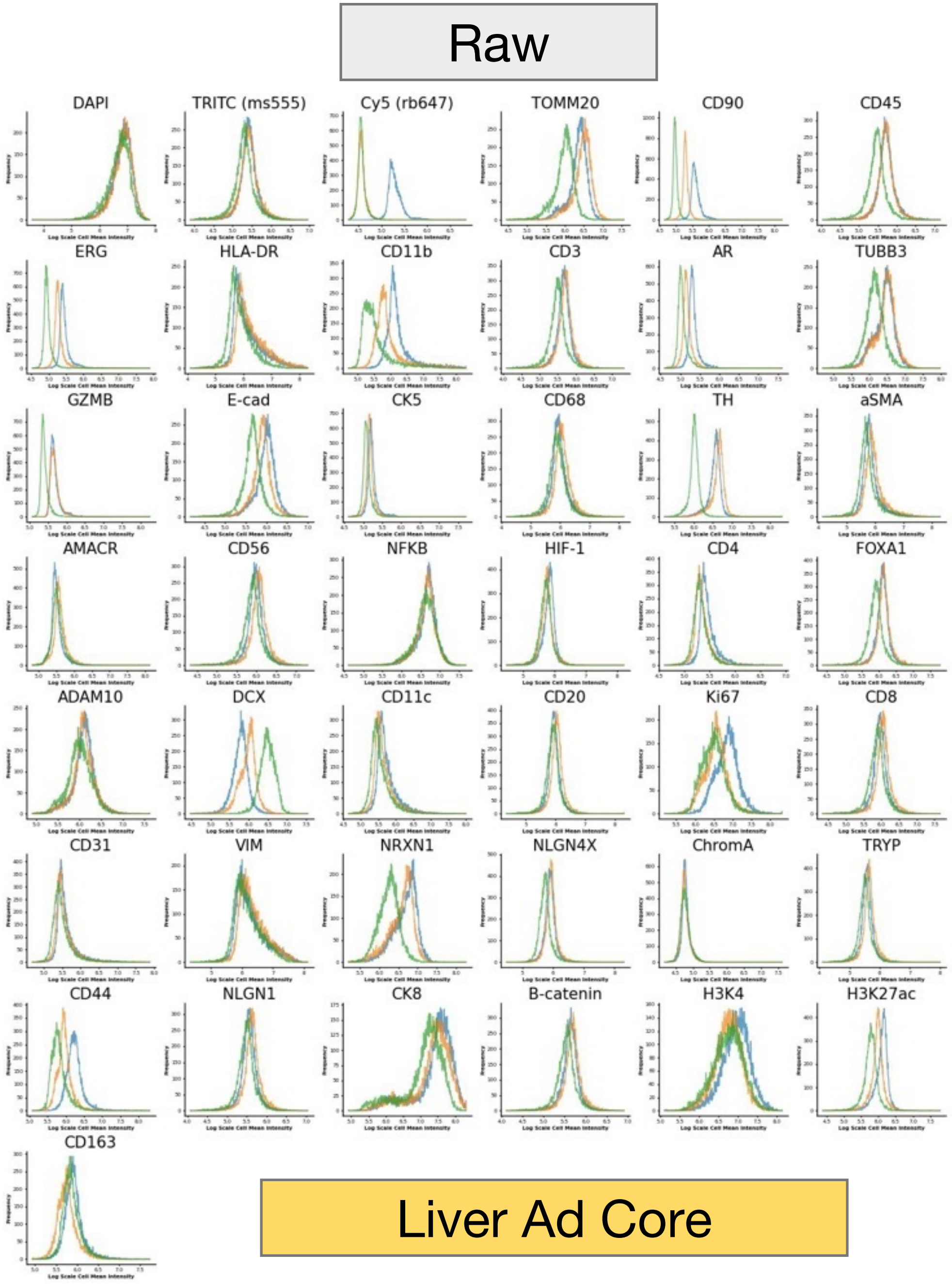


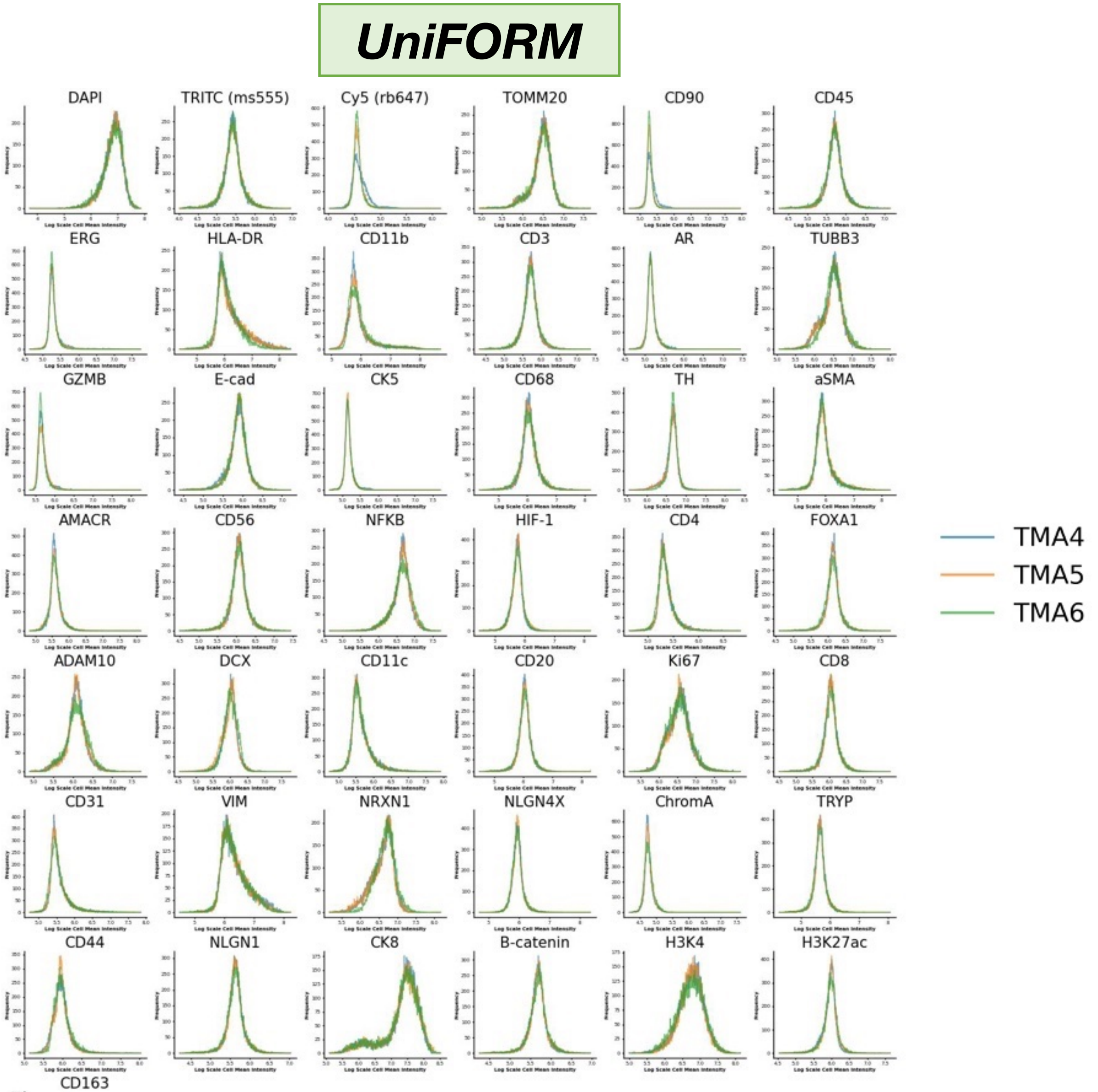


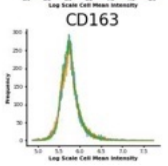


**Figure S3f. TMA-Lunaphore feature-level normalization on Liver Ad Core.** Raw data shows significant technical variation on the same slide across batches. UniFORM accurately aligns distributions while preserving biology.

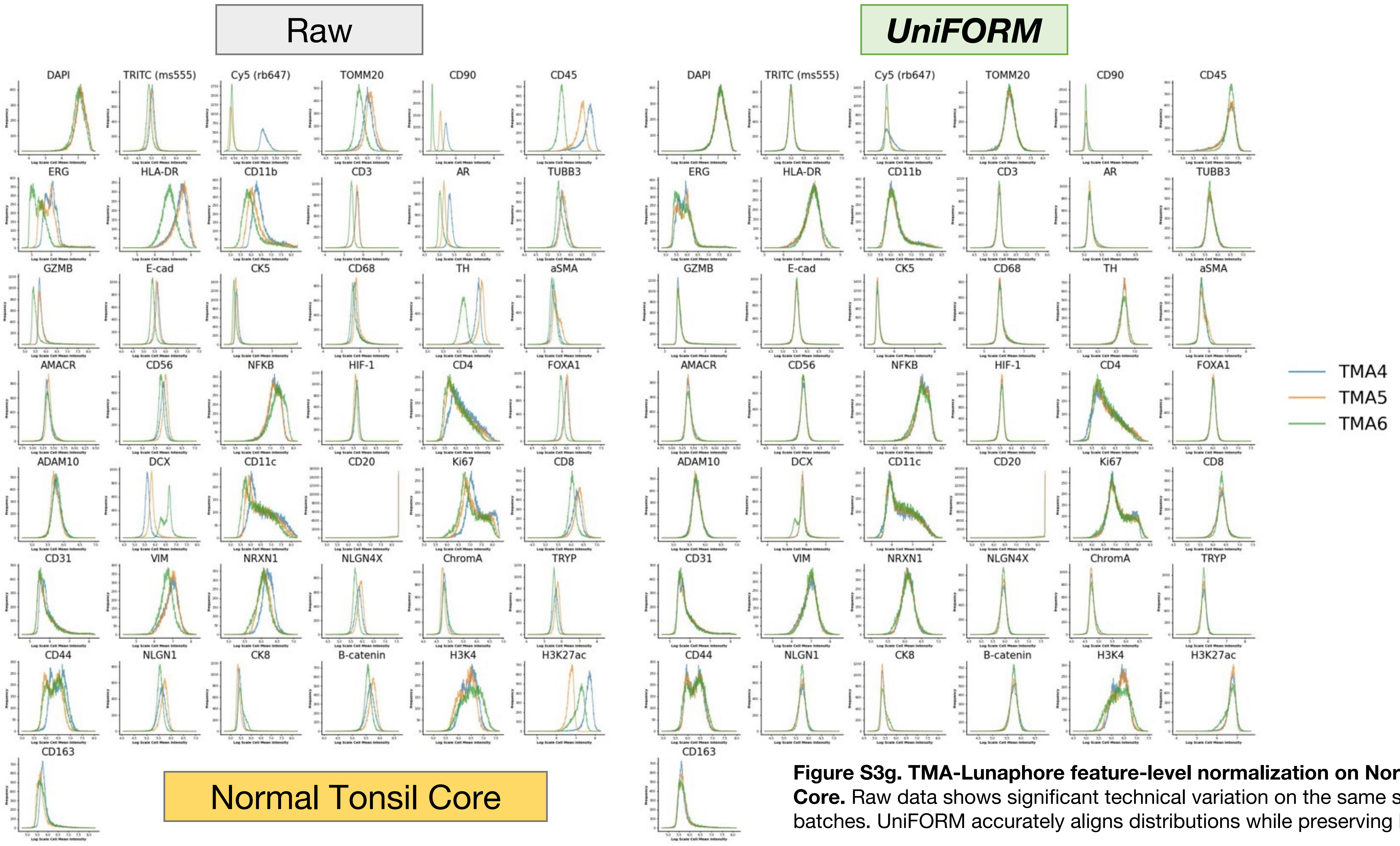


**Figure S3g. TMA-Lunaphore feature-level normalization on Normal Tonsil Core.** Raw data shows significant technical variation on the same slide across batches. UniFORM accurately aligns distributions while preserving biology.

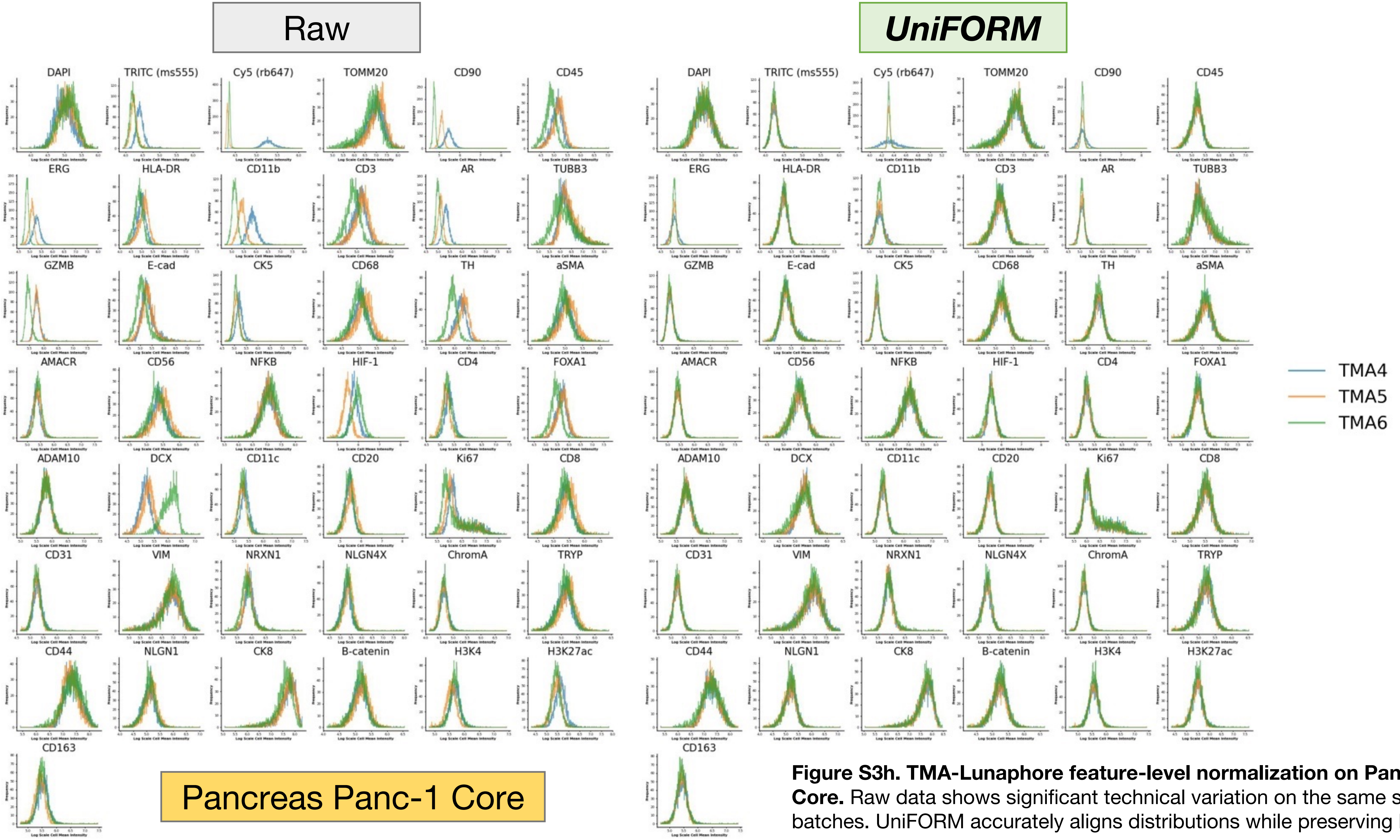


**Figure S3h. TMA-Lunaphore feature-level normalization on Pancreas Panc-1 Core.** Raw data shows significant technical variation on the same slide across batches. UniFORM accurately aligns distributions while preserving biology.

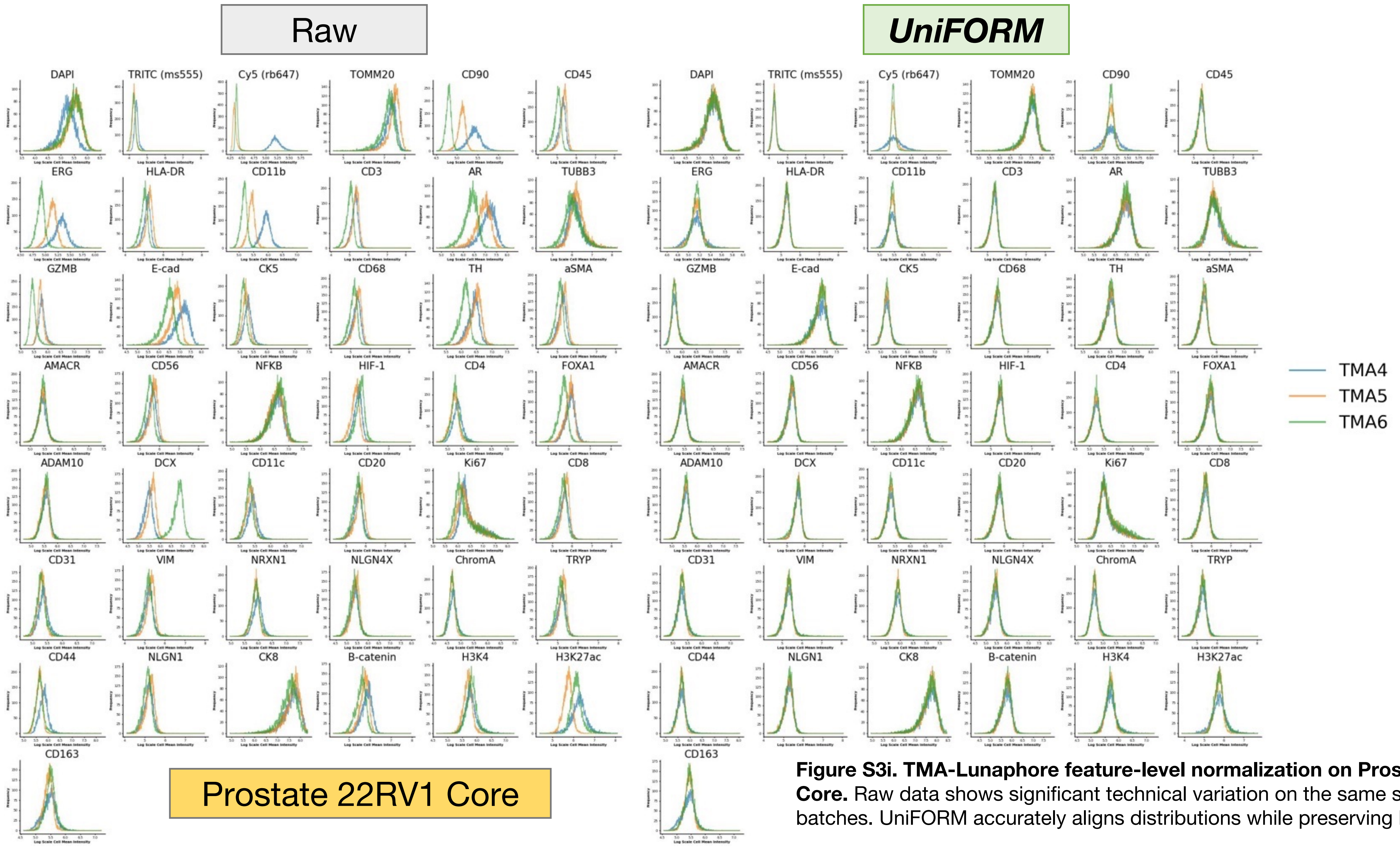


**Figure S3i. TMA-Lunaphore feature-level normalization on Prostate 22RV1 Core.** Raw data shows significant technical variation on the same slide across batches. UniFORM accurately aligns distributions while preserving biology.

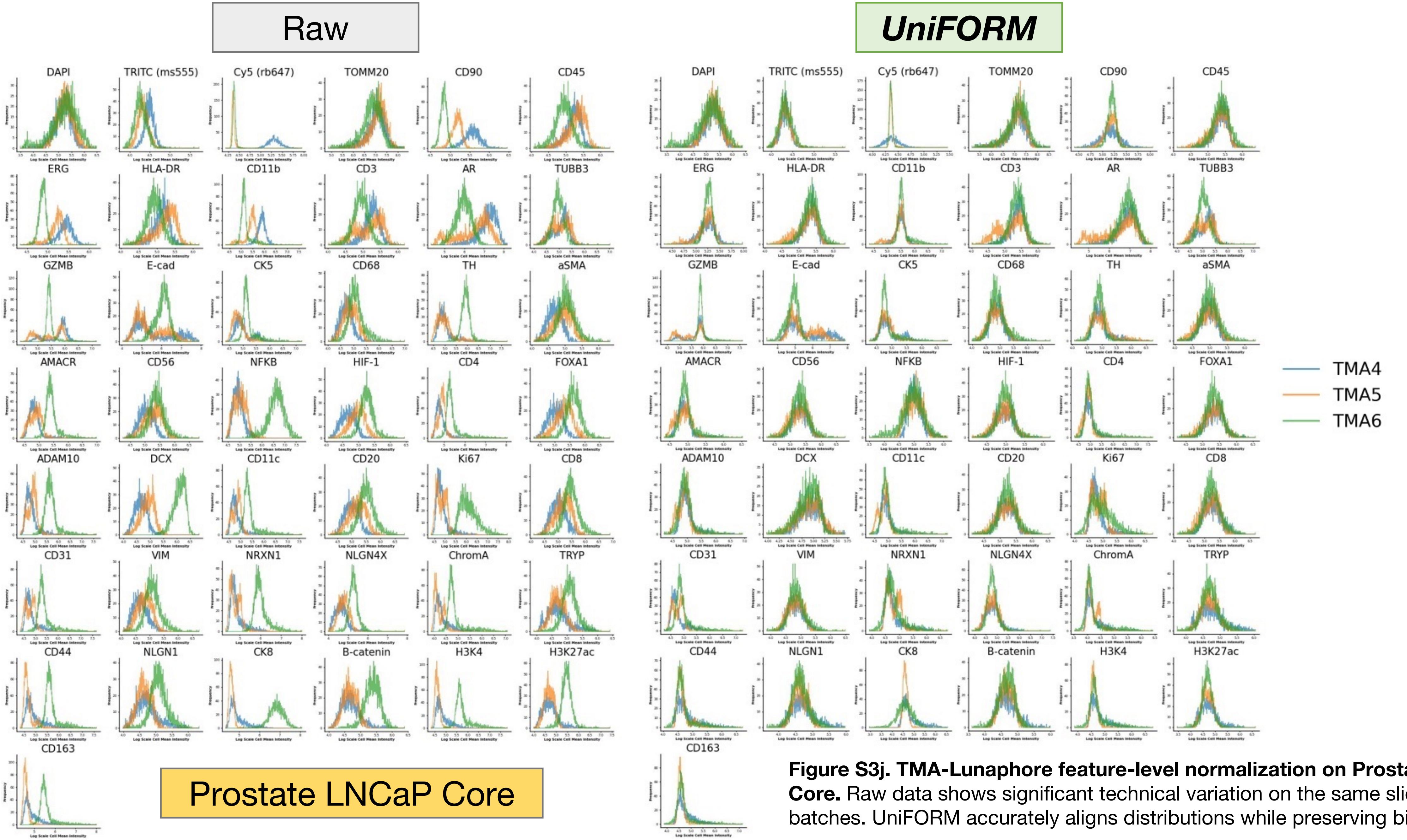


**Figure S3j. TMA-Lunaphore feature-level normalization on Prostate LNCaP Core.** Raw data shows significant technical variation on the same slide across batches. UniFORM accurately aligns distributions while preserving biology.

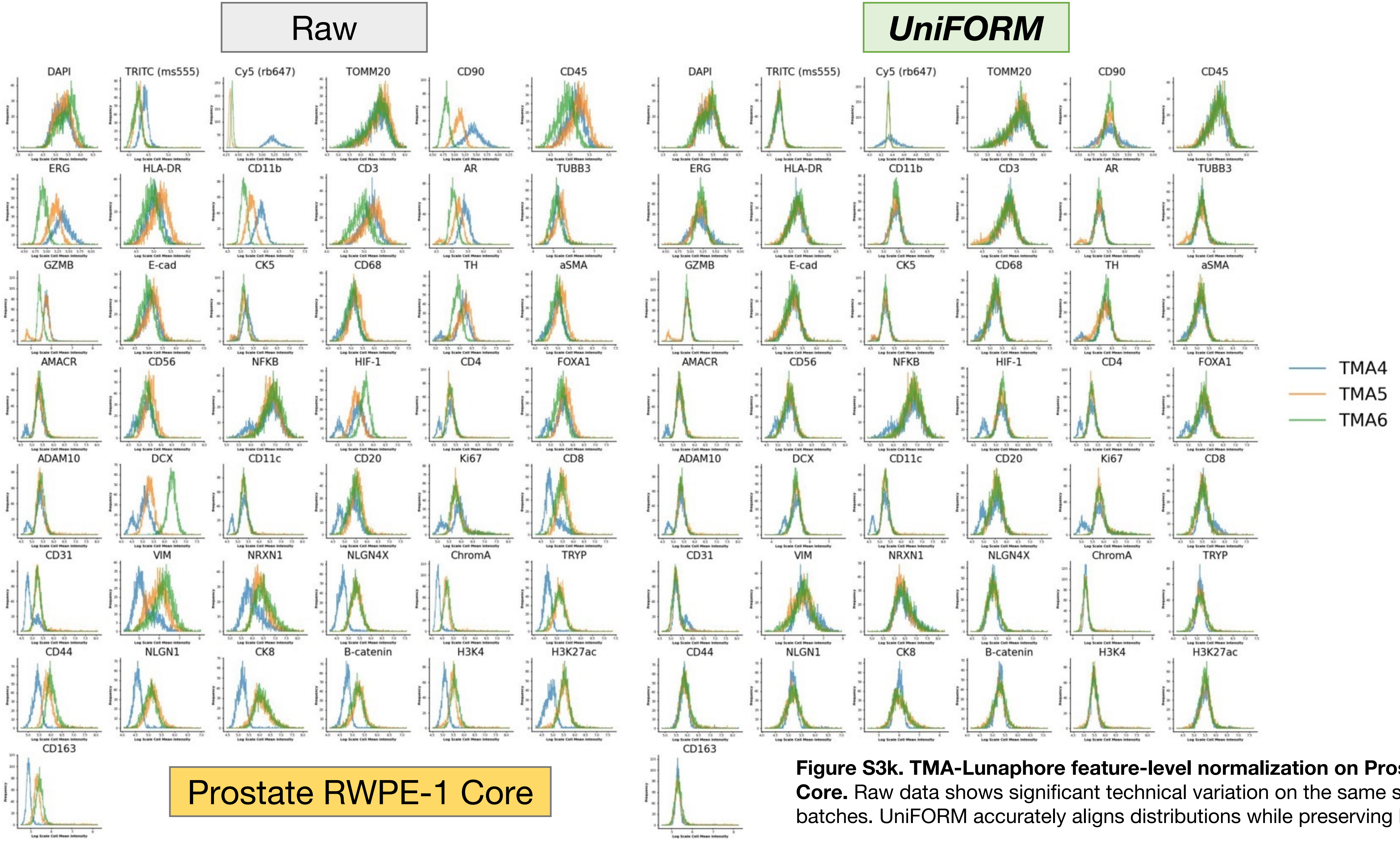


**Figure S3k. TMA-Lunaphore feature-level normalization on Prostate RWPE-1 Core.** Raw data shows significant technical variation on the same slide across batches. UniFORM accurately aligns distributions while preserving biology.

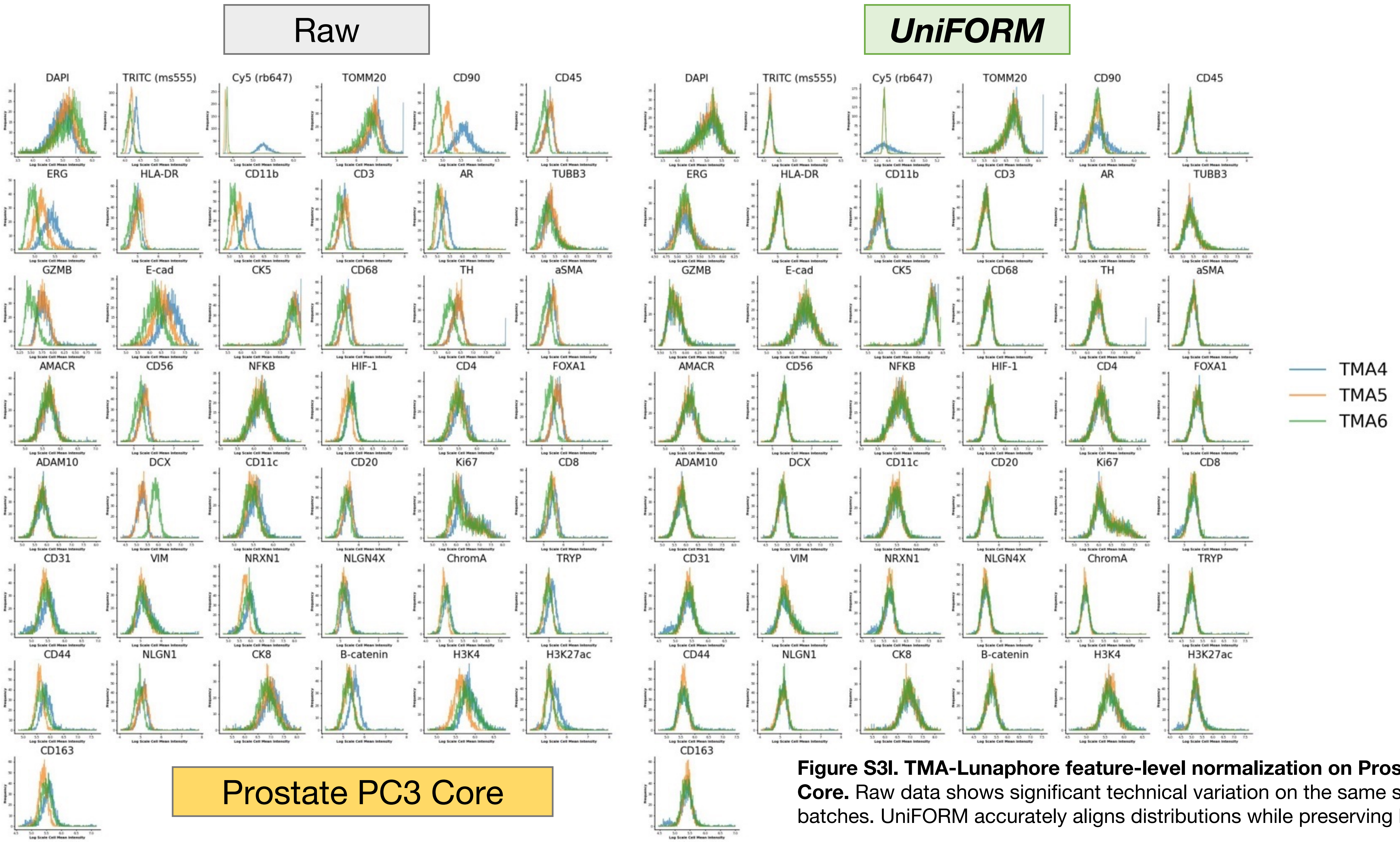


**Figure S3I. TMA-Lunaphore feature-level normalization on Prostate PC3 Core.** Raw data shows significant technical variation on the same slide across batches. UniFORM accurately aligns distributions while preserving biology.

| Sample Pair | Raw | | | UniFORM | | | MxNorm | | | z-score | | | ComBat | | | MeanDiv | | |
|---|---|---|---|---|---|---|---|---|---|---|---|---|---|---|---|---|---|---|
| | kBET | Chi-square | p-value | kBET | Chi-square | p-value | kBET | Chi-square | p-value | kBET | Chi-square | p-value | kBET | Chi-square | p-value | kBET | Chi-square | p-value |
| PRAD-02 & PRAD-14 | 0.87496 | 24.659 | 0.05509 | **0.91320** | 4.2913 | 0.35847 | 0.88442 | 13.550 | 0.21950 | 0.89178 | 10.987 | 0.26118 | 0.88295 | 6.7732 | 0.29147 | 0.90841 | **3.9433** | **0.36155** |
| PRAD-01 & PRAD-02 | 0.00064 | 24.862 | 0.00138 | 0.67208 | 5.3327 | 0.31685 | 0.43894 | 9.5486 | 0.18245 | 0.38175 | 10.160 | 0.15637 | 0.42424 | 10.448 | 0.19091 | **0.70299** | **4.0201** | **0.32868** |
| PRAD-05 & PRAD-12 | 0.00059 | 24.915 | 0.00017 | **0.61642** | **5.8098** | **0.24522** | 0.49168 | 6.8864 | 0.18599 | 0.47464 | 9.0445 | 0.21229 | 0.27958 | 13.985 | 0.11999 | 0.59799 | 6.1684 | 0.23986 |
| PRAD-07 & PRAD-19 | 0.00128 | 24.864 | 0.00375 | **0.58262** | **8.9142** | **0.23530** | 0.27883 | 11.450 | 0.10426 | 0.25590 | 17.858 | 0.07892 | 0.32481 | 15.534 | 0.10738 | 0.51087 | 10.348 | 0.17366 |
| PRAD-02 & PRAD-07 | 0.00143 | 5.7325 | 0.00051 | **0.56263** | **5.7325** | **0.23605** | 0.34209 | 11.945 | 0.13959 | 0.03052 | 22.231 | 0.01080 | 0.06202 | 21.228 | 0.02096 | 0.35778 | 9.9925 | 0.16052 |

**Table S2: kBET Analysis Results Across Normalization Methods.** kBET analysis was performed on five pairs of PRAD-CyCIF samples selected for similar cell compositions to ensure fair assessment of batch correction performance. UniFORM consistently demonstrated strong performance across all three kBET evaluation metrics, achieving the highest average acceptance rate among all methods. While mean division showed slightly better scores in two sample pairs, this improvement may result from artificial homogenization that suppresses both technical variation and biologically meaningful differences, potentially compromising biological fidelity.

| Sample Pair | Raw | UniFORM | MxNorm | z-score | ComBat | MeanDiv |
|---|---|---|---|---|---|---|
| PRAD-02 & PRAD-14 | 0.42690 | 0.48853 | -0.01072 | 0.46040 | **0.48896** | 0.40923 |
| PRAD-01 & PRAD-02 | 0.25328 | 0.34803 | -0.01928 | 0.31553 | **0.42326** | 0.27119 |
| PRAD-05 & PRAD-12 | 0.35813 | **0.46952** | -0.01747 | 0.24083 | 0.27185 | 0.23688 |
| PRAD-07 & PRAD-19 | 0.43697 | **0.55053** | -0.02566 | 0.43746 | 0.47941 | 0.48244 |
| PRAD-02 & PRAD-07 | 0.53156 | **0.57780** | -0.03393 | 0.47141 | 0.52634 | 0.47710 |

**Table S3: Silhouette Coefficient Analysis Results Across Normalization Methods.** Silhouette analysis was conducted on the same five pairs of PRAD-CyCIF samples used in the kBET analysis to evaluate the separation of mutually exclusive cell types. UniFORM consistently achieved high silhouette coefficients across all pairs, with the highest average score among the methods, reflecting robust preservation of biologically meaningful cluster structure. While ComBat yielded slightly higher scores in two sample pairs, this may result from excessive compression of the data, which artificially reduces intra-cluster distances and inflates the silhouette score without necessarily preserving biological fidelity.

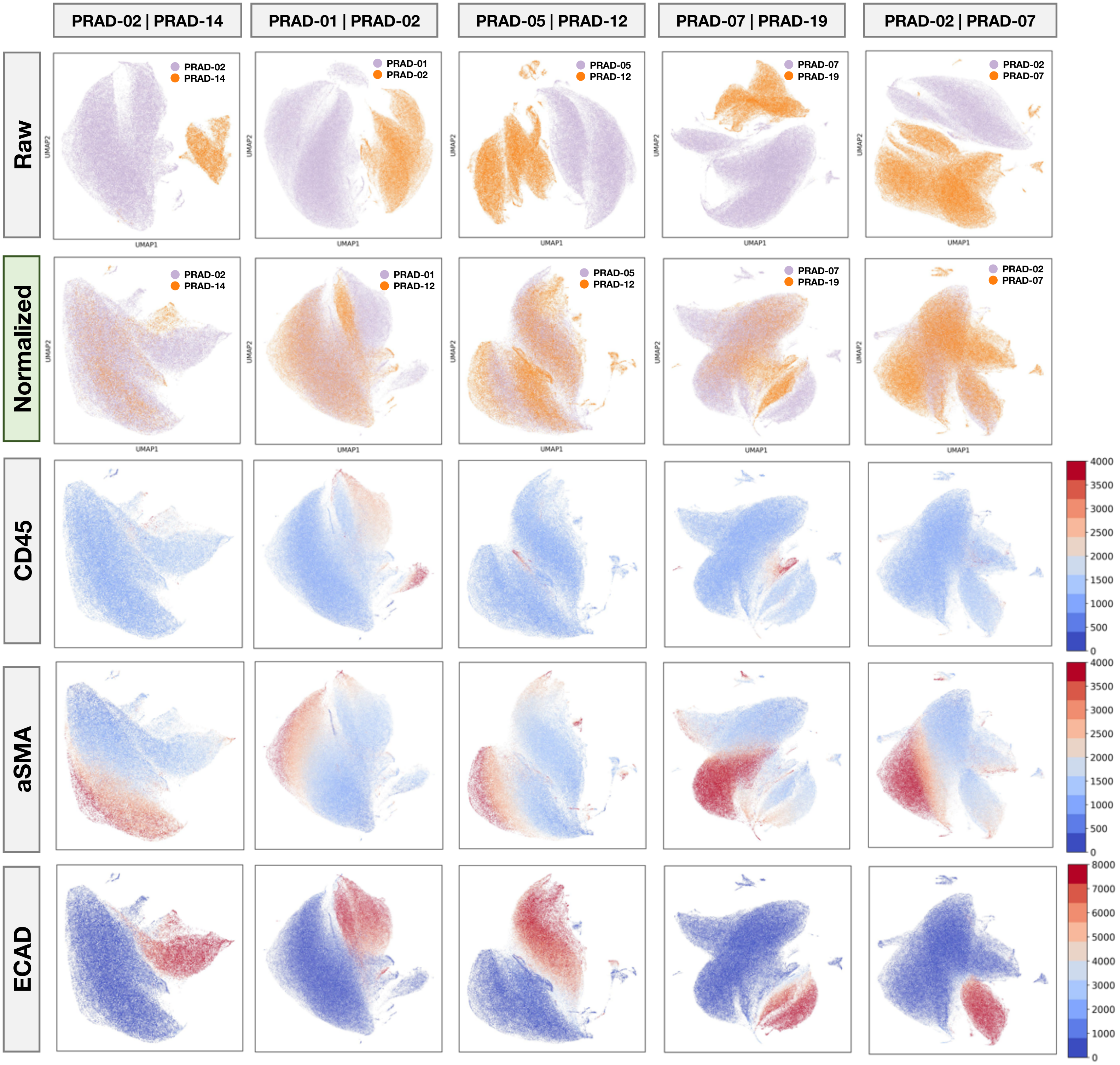


**Figure S4: Improved data integration and marker clustering with UniFORM normalization.** UMAP visualizations of raw and UniFORM-normalized data from five PRAD-CyCIF patient pairs, colored by sample IDs, highlight enhanced sample mixing post-normalization. The UniFORM-normalized data demonstrates significantly improved clustering of cells with higher intensity values for key markers, including CD45, α-SMA, and ECAD, reflecting better biological alignment across samples

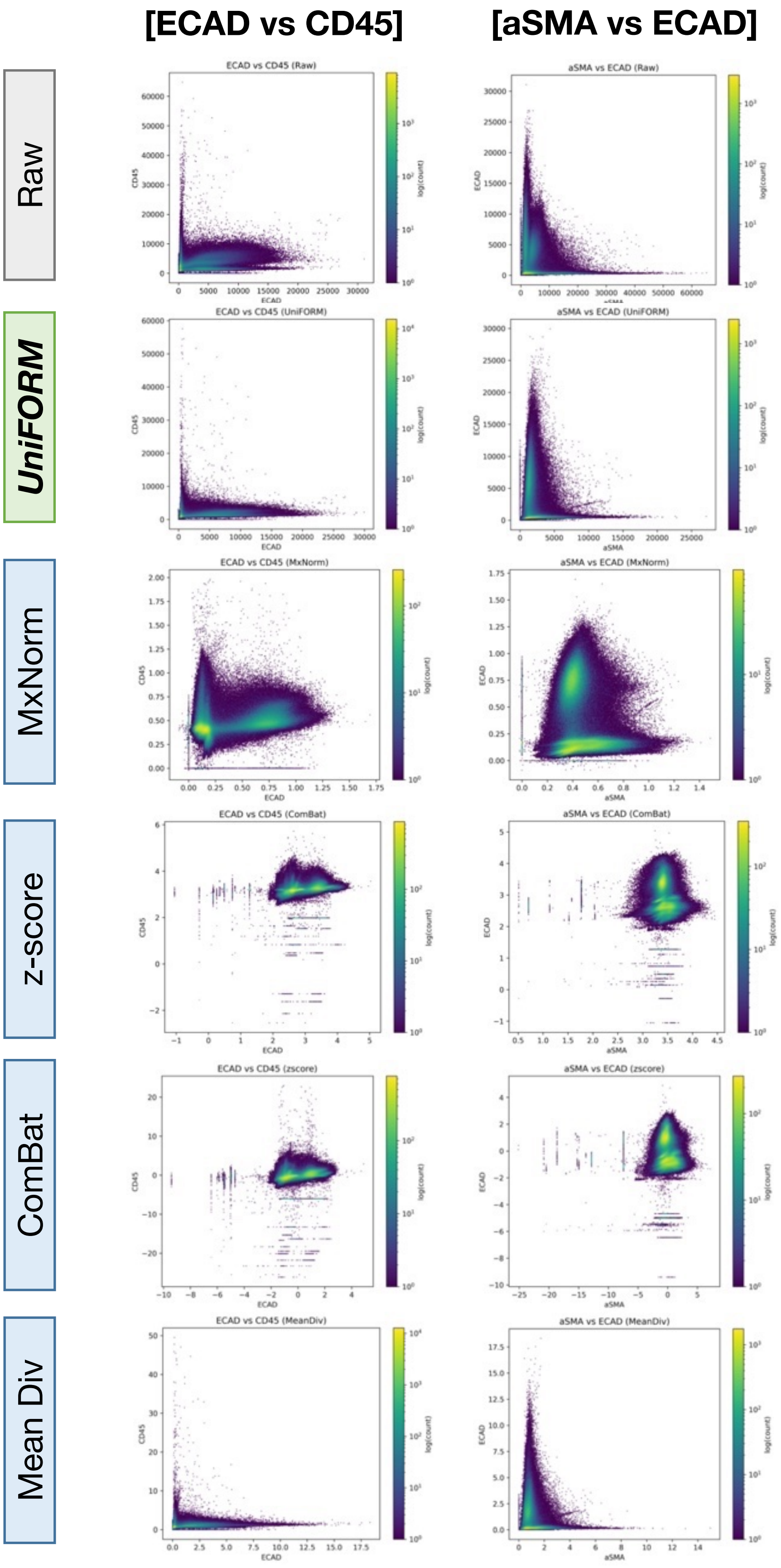


**Figure S5. Scatter Plots of Mutually Exclusive Marker Expression.** Scatter plots of mutually exclusive marker pairs—ECAD vs. CD45 and aSMA vs. ECAD—illustrate how different normalization methods affect marker exclusivity. UniFORM and mean division substantially improved the separation between mutually exclusive markers, restoring expected biological patterns. In contrast, MxNorm (registration-based), z-score, and ComBat distort marker expression relationships, reducing exclusivity and potentially compromising biological interpretability. The compressed expression ranges observed in z-score and ComBat may artificially inflate silhouette scores without preserving meaningful biological variation.

| Marker | UniFORM | | MxNorm | | z-score | | ComBat | | MeanDiv | |
|---|---|---|---|---|---|---|---|---|---|---|
| | Mean % Change | Std Dev | Mean % Change | Std Dev | Mean % Change | Std Dev | Mean % Change | Std Dev | Mean % Change | Std Dev |
| CD31 | 3.8807 | **1.3940** | 2.7941 | 2.7362 | **2.7219** | 2.2739 | 3.5785 | 2.3865 | 2.7266 | 1.5815 |
| CD45 | **2.2817** | **1.2222** | 19.219 | 9.6910 | 7.3230 | 6.2264 | 18.135 | 6.7914 | 15.638 | 7.0022 |
| CD68 | **3.6009** | **4.5310** | 10.860 | 8.1556 | 5.4402 | 7.1554 | 7.3337 | 9.1228 | 5.8881 | 7.7298 |
| CD4 | **2.1124** | **1.1236** | 13.357 | 7.7832 | 5.1557 | 6.1330 | 14.355 | 5.8997 | 8.6208 | 5.7388 |
| FOXP3 | **0.91994** | **0.40661** | 3.4118 | 2.3611 | 1.7271 | 0.73822 | 1.9501 | 1.3005 | 1.3724 | 1.1603 |
| CD8a | **1.2097** | **1.2520** | 7.5454 | 6.0071 | 2.2988 | 2.1913 | 4.4001 | 2.4769 | 4.1304 | 2.3902 |
| CD45RO | **1.2807** | **1.2753** | 11.004 | 6.6268 | 5.5626 | 3.9647 | 5.3298 | 4.2621 | 4.6460 | 3.9447 |
| CD20 | **1.6591** | 1.5589 | 2.8773 | 1.8676 | 2.2067 | **1.0494** | 3.1440 | 3.0739 | 1.7791 | 1.8990 |
| PD-L1 | **2.3122** | 2.2189 | 8.3193 | 3.977 | 4.2656 | 2.3773 | 4.2459 | 1.6703 | 2.8229 | **1.2632** |
| CD3 | **1.1070** | **0.67126** | 8.8278 | 3.8668 | 3.3427 | 1.8078 | 3.3892 | 2.1547 | 3.3807 | 2.5188 |
| CD163 | **4.0568** | **1.7896** | 9.6176 | 3.5602 | 4.7440 | 3.2642 | 5.4731 | 4.4389 | 6.8842 | 4.1504 |
| ECAD | **1.3844** | **0.72127** | 18.212 | 11.813 | 3.9440 | 2.3471 | 9.8169 | 3.6952 | 12.539 | 24.973 |
| PD1 | **5.4169** | **4.8181** | 9.3586 | 10.48 | 7.9479 | 5.0211 | 7.2189 | 6.3596 | 6.9638 | 8.1048 |
| Ki67 | **1.2271** | **1.2199** | 11.885 | 6.5472 | 5.5387 | 3.9662 | 7.2577 | 4.5328 | 11.066 | 4.6917 |
| Pan-CK | **2.1303** | **1.0093** | 16.267 | 13.333 | 5.9269 | 4.2509 | 14.791 | 6.6299 | 7.2812 | 4.4392 |
| aSMA | 3.8978 | 4.3698 | 11.169 | 6.5845 | **3.7237** | 2.2731 | 4.6741 | **2.0914** | 4.9642 | 2.2414 |

**Table S4a: Positive Population Change Analysis for CRC-ORION dataset.** UniFORM dominates in having the lowest mean percentage change and standard deviation among all methods.

| Marker | UniFORM | | MxNorm | | z-score | | ComBat | | MeanDiv | |
|---|---|---|---|---|---|---|---|---|---|---|
| | Mean % Change | Std Dev | Mean % Change | Std Dev | Mean % Change | Std Dev | Mean % Change | Std Dev | Mean % Change | Std Dev |
| EPCAM | 7.7091 | 6.0188 | 7.9861 | 4.1146 | 5.9916 | 3.7327 | **3.8079** | **3.2534** | 22.242 | 9.3646 |
| CD56 | 1.9391 | 4.2764 | 3.8908 | **2.1719** | **1.5639** | 3.8530 | 2.4082 | 4.4461 | 14.980 | 3.8385 |
| CD45 | 7.5275 | 4.9072 | 13.468 | 9.3487 | 6.6940 | 4.7590 | 5.1941 | 3.8894 | **4.0705** | **3.7570** |
| aSMA | **6.7804** | **5.1111** | 19.254 | 11.295 | 10.550 | 14.234 | 22.890 | 12.300 | 16.131 | 8.2366 |
| ChromA | **2.7201** | **5.4197** | 9.4856 | 13.618 | 5.3158 | 8.3392 | 5.9512 | 11.297 | 5.3989 | 10.109 |
| CK14 | 0.7459 | 1.5869 | 1.9219 | **1.2432** | **0.7350** | 1.4290 | 1.0341 | 1.2751 | 4.3356 | 2.3652 |
| Ki67 | **1.9166** | **2.5194** | 5.9232 | 13.668 | 5.1354 | 15.463 | 5.2582 | 15.302 | 4.3646 | 13.726 |
| GZMB | 2.1503 | 3.7004 | 3.2859 | **2.4451** | 1.9156 | 3.4084 | **1.8701** | 2.7989 | 5.3294 | 2.5839 |
| ECAD | 2.9117 | **1.3709** | 17.022 | 11.053 | 4.3243 | 6.3498 | 4.3131 | 6.2733 | **2.0499** | 1.4043 |
| PD1 | 3.4994 | 5.9252 | 4.9479 | 5.8721 | 3.2809 | 5.8419 | **3.0950** | **5.0202** | 12.962 | 6.8823 |
| CD31 | **3.2084** | **2.8055** | 8.4386 | 11.791 | 4.7342 | 10.633 | 5.7803 | 12.643 | 4.5534 | 9.5648 |
| CD45RA | 1.4004 | 2.6855 | 20.785 | 5.9765 | **1.0231** | **2.3166** | 1.3204 | 2.5947 | 3.2153 | 2.7192 |
| HLADRB1 | **2.6441** | **4.0506** | 12.692 | 10.657 | 7.6595 | 10.614 | 7.9911 | 10.330 | 5.9993 | 8.6299 |
| CD3 | 4.9536 | **5.3458** | 15.634 | 12.187 | 6.2817 | 14.724 | **4.5291** | 9.3660 | 9.3239 | 8.8390 |
| FOXA1 | **1.4572** | 2.7276 | 2.9280 | 3.0964 | 1.5438 | 2.6850 | 2.0928 | 2.9014 | 5.6331 | **2.0071** |
| CDX2 | 6.3853 | 3.1297 | **2.6099** | **2.3422** | 10.125 | 3.0272 | 4.3563 | 3.7742 | 5.0081 | 2.9358 |
| CD20 | **1.9214** | **3.7294** | 17.442 | 18.518 | 8.7831 | 19.371 | 9.1211 | 19.596 | 5.8570 | 13.937 |
| NOTCH1 | **4.7104** | **5.6383** | 21.342 | 19.632 | 38.493 | 18.141 | 15.238 | 22.597 | 14.464 | 20.351 |

**Table S4b: Positive Population Change Analysis for PRAD-CyCIF dataset.** UniFORM still has the lowest mean percentage change and standard deviation in most markers despite the fact that PRAD-CyCIF is a more heterogeous dataset.

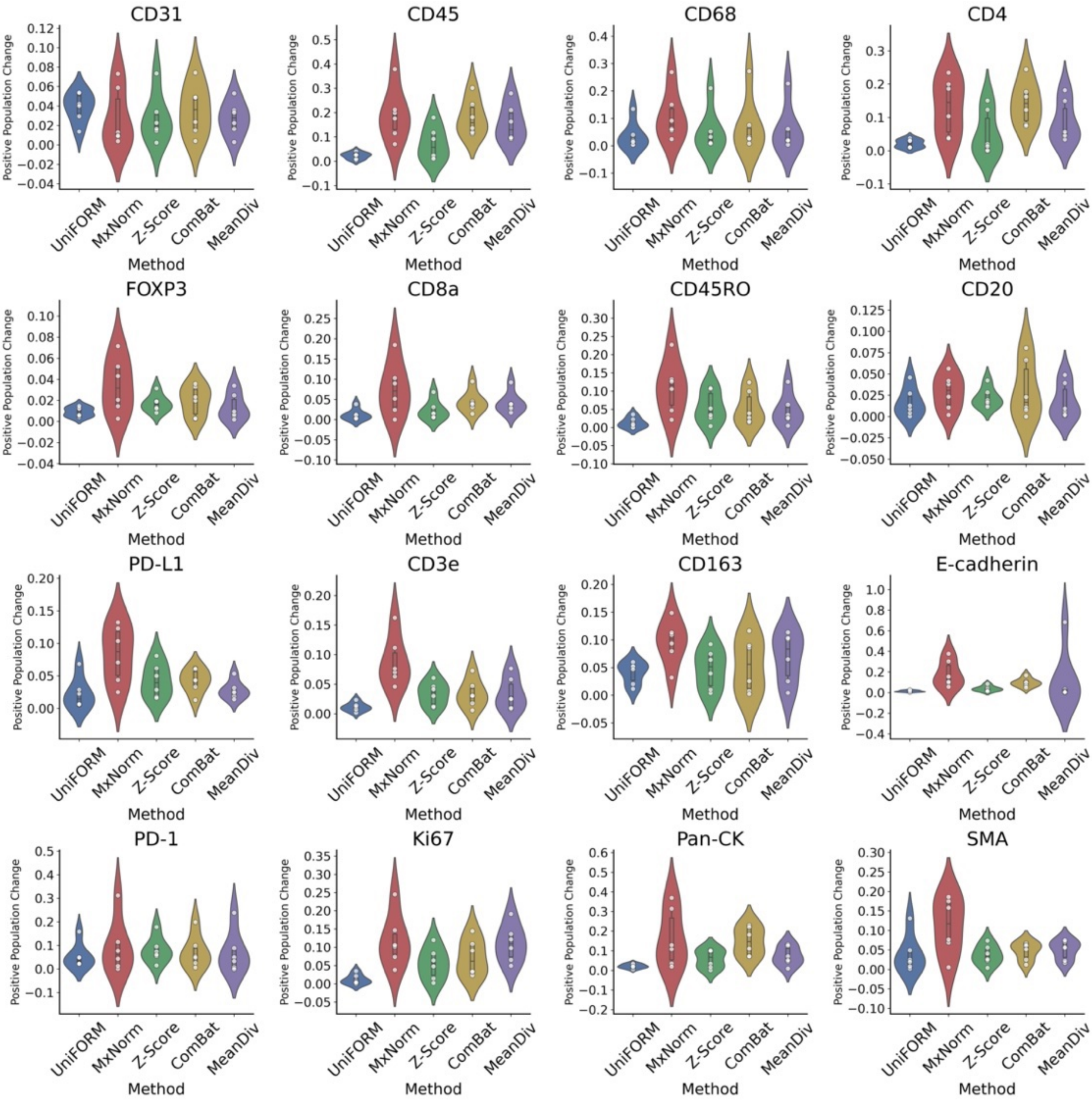


**Figure S6a: Violin plots for the positive population change analysis for the CRC-ORION dataset.** UniFORM consistently shows the lowest central tendency and spread in all markers across methods.

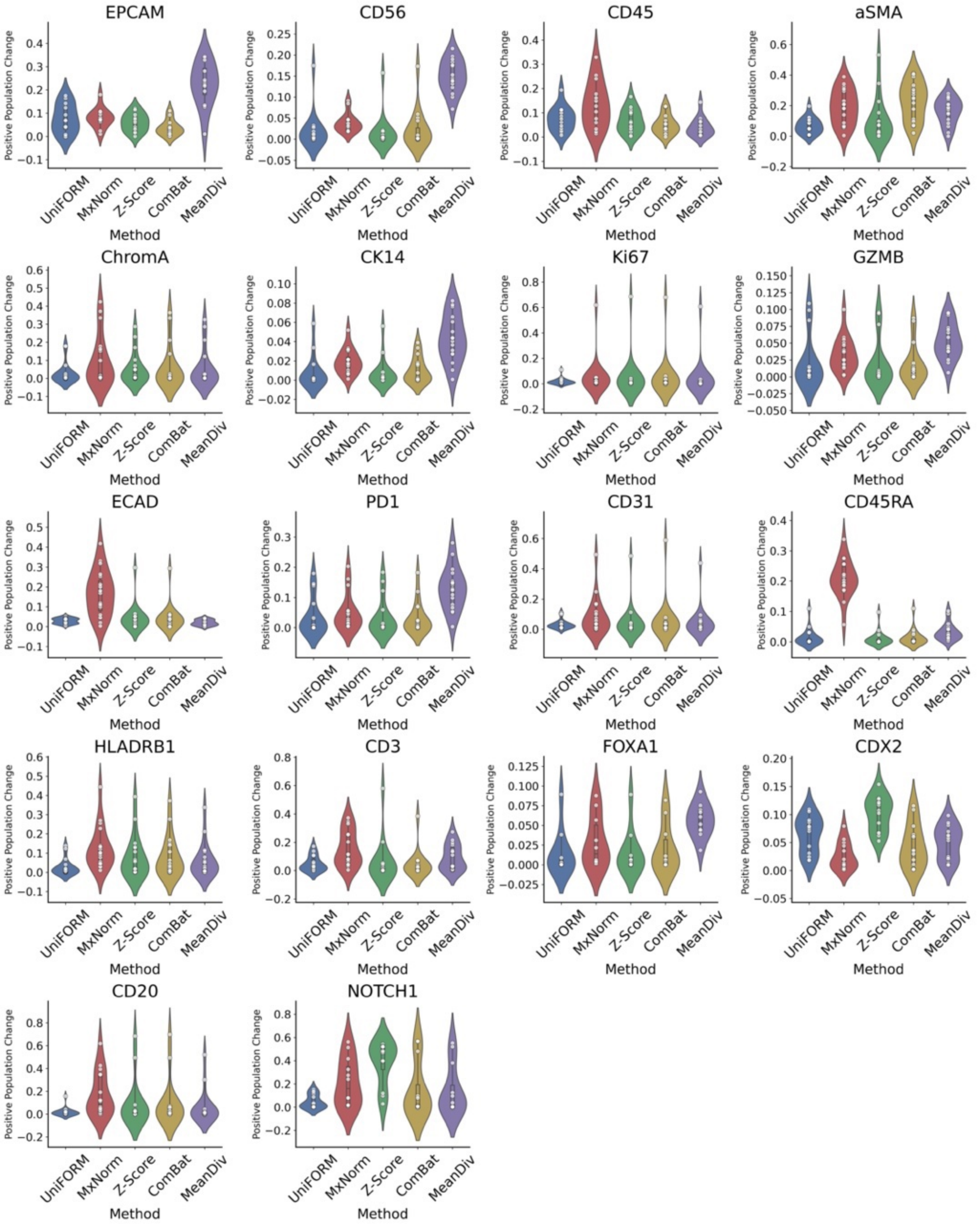


**Figure S6b: Violin plots for the positive population change analysis for the PRAD-CyCIF dataset.** UniFORM shows the lowest central tendency and spread in most markers across methods. UniFORM has relatively higher percentage change and SD due to data being very heterogenous and UniFORM's nature to preserve distribution shape.

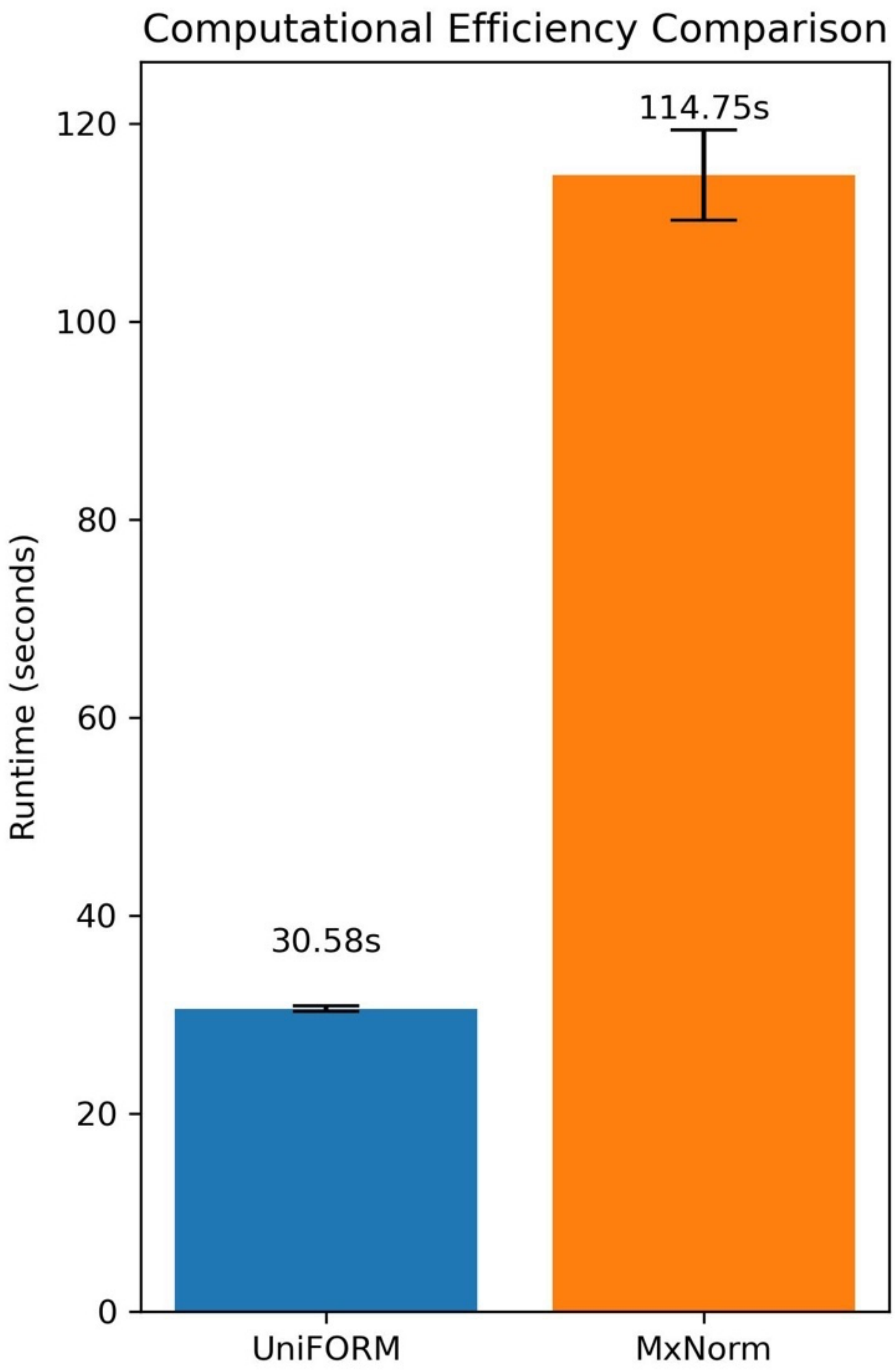


**Figure S7: Comparison of computational efficiency between UniFORM and MxNorm.** Average runtime (in seconds) was measured across five independent runs for each method using the PRAD-CyCIF dataset and hardware (Apple MacBook Pro, M2 Max chip, 32GB RAM). UniFORM demonstrated significantly lower runtime (30.58 ± 0.30 s) compared to MxNorm (114.75 ± 4.35 s), indicating a nearly fourfold improvement in computational efficiency. Error bars represent standard deviation.

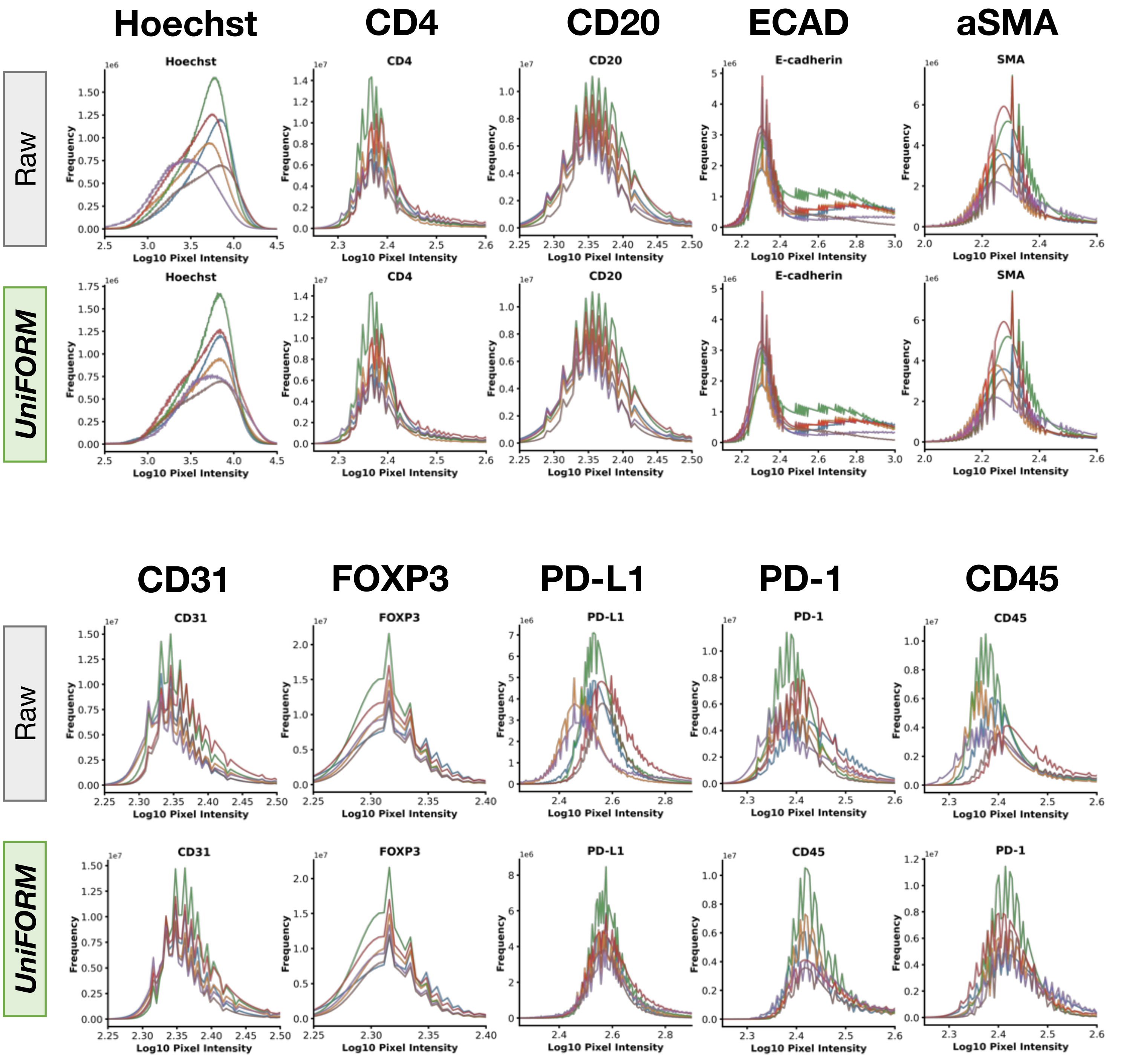


**Figure S8a: CRC-ORION pixel-level normalization.** Some markers such as CD45RO, PD-L1, and PD-1 show variations in cell intensity distribution despite being from the same batch. UniFORM harmonizes inter-batch variations while preserving the distribution shape and positive population counts.

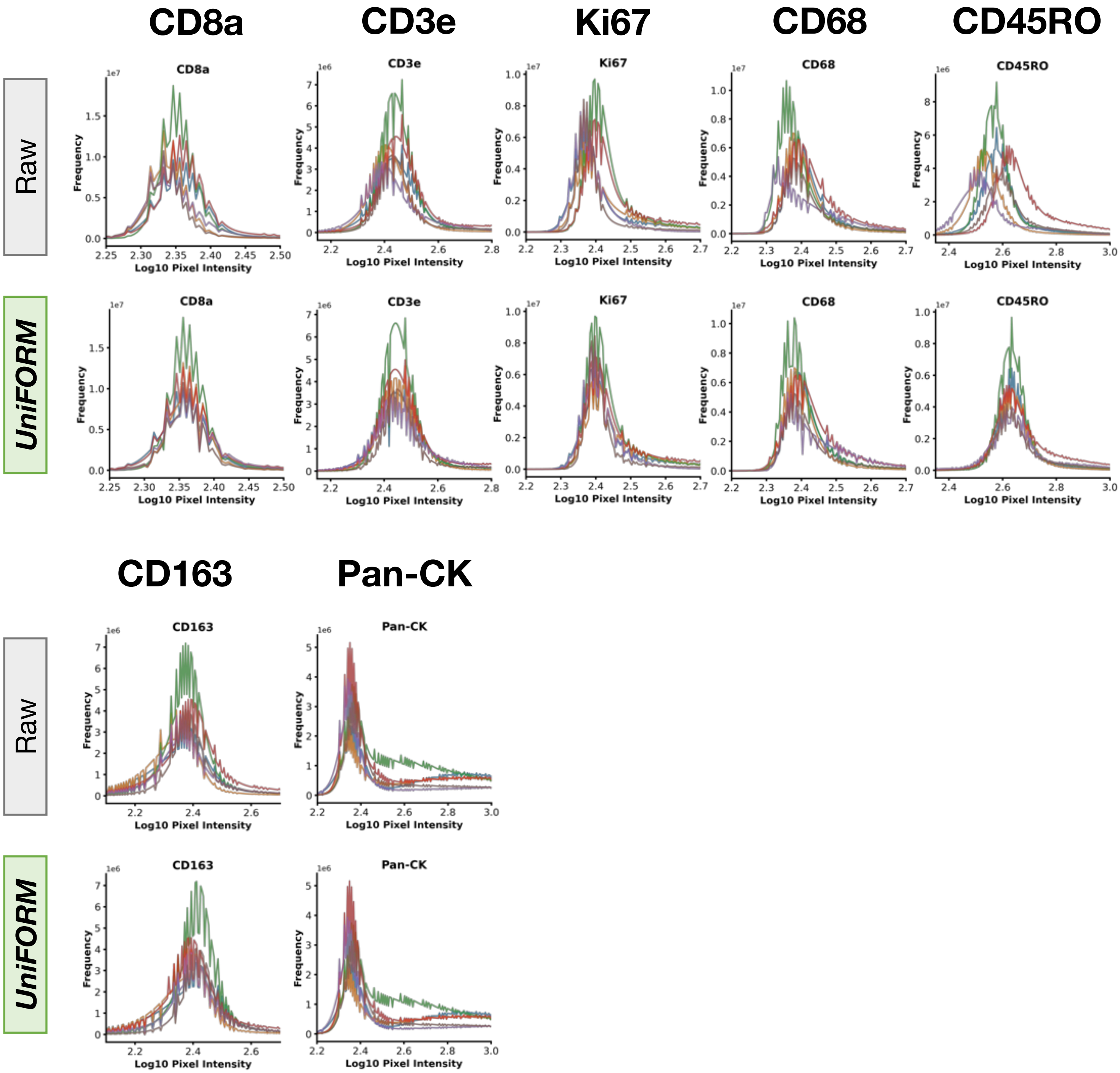


**Figure S8a: CRC-ORION pixel-level normalization.** Some markers such as CD45RO, PD-L1, and PD-1 show variations in cell intensity distribution despite being from the same batch. UniFORM harmonizes inter-batch variations while preserving the distribution shape and positive population counts.

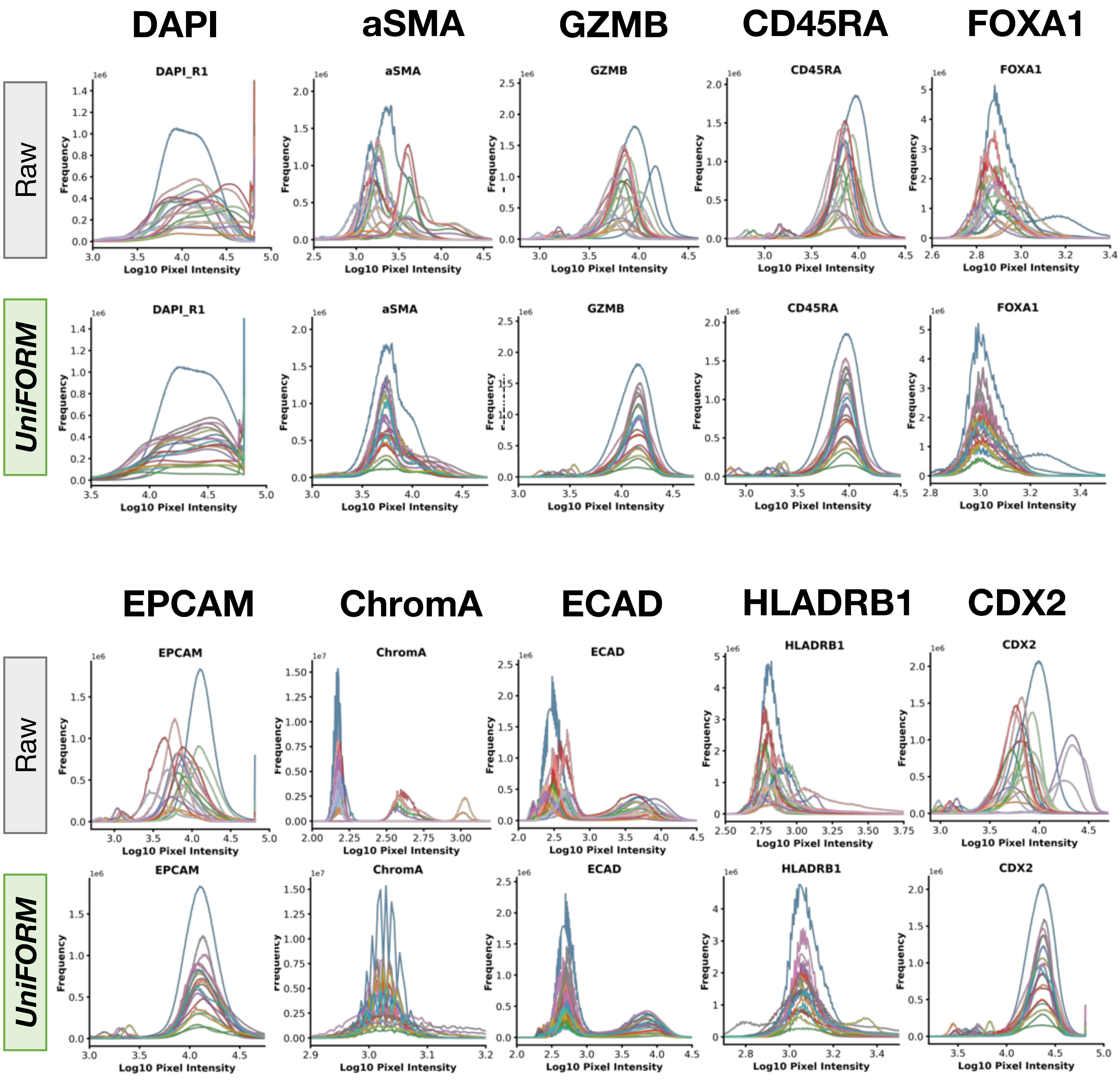


**Figure S9a: PRAD-CyCIF pixel-level normalization.** PRAD raw data shows significant batch effect and heterogenous distribution pattern. UniFORM accurately and efficiently aligns the distribution while preserving the distribution shape and positive population counts.

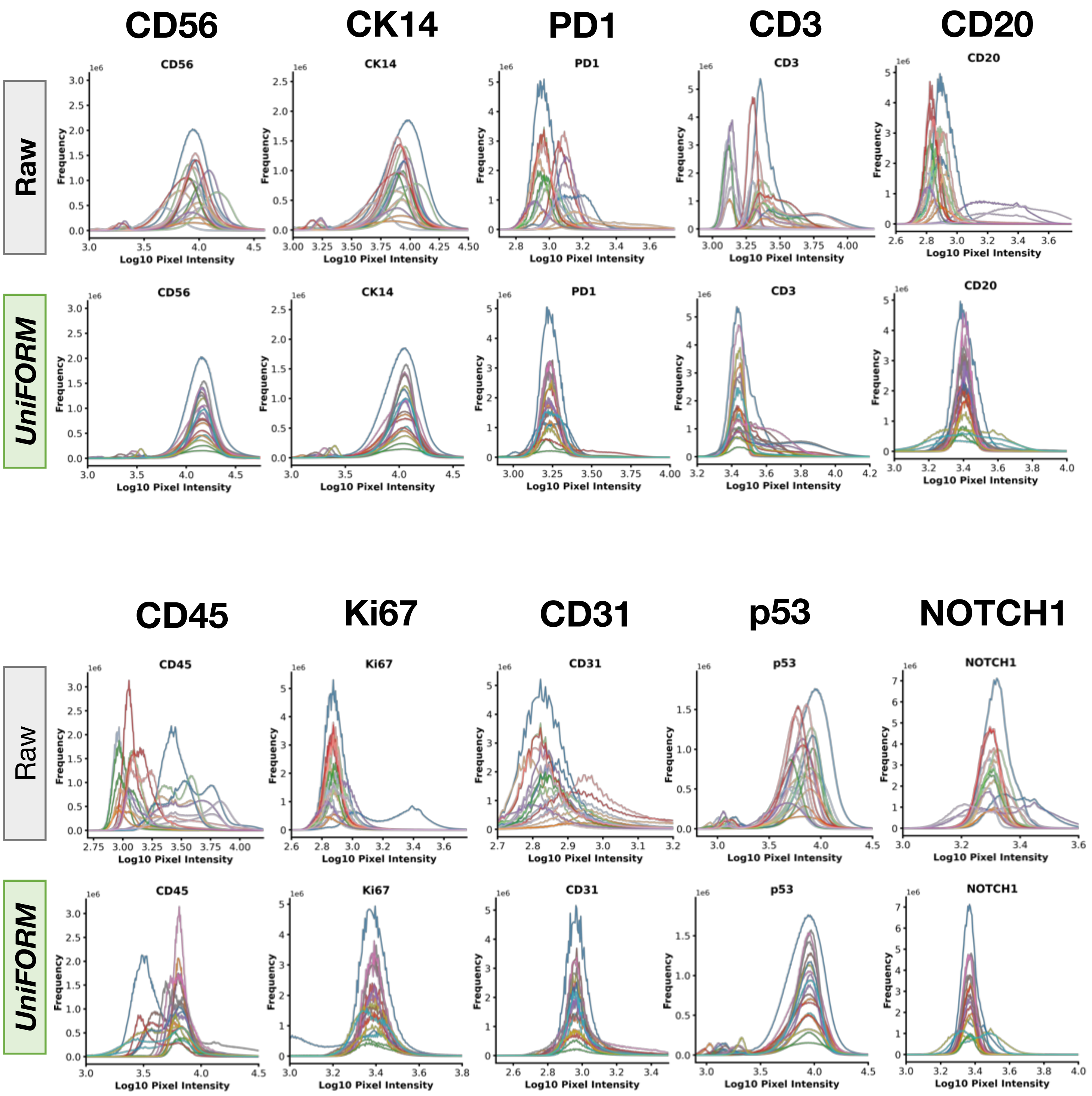


**Figure S9a: PRAD-CyCIF pixel-level normalization.** PRAD raw data shows significant batch effect and heterogenous distribution pattern. UniFORM accurately and efficiently aligns the distribution while preserving the distribution shape and positive population counts.

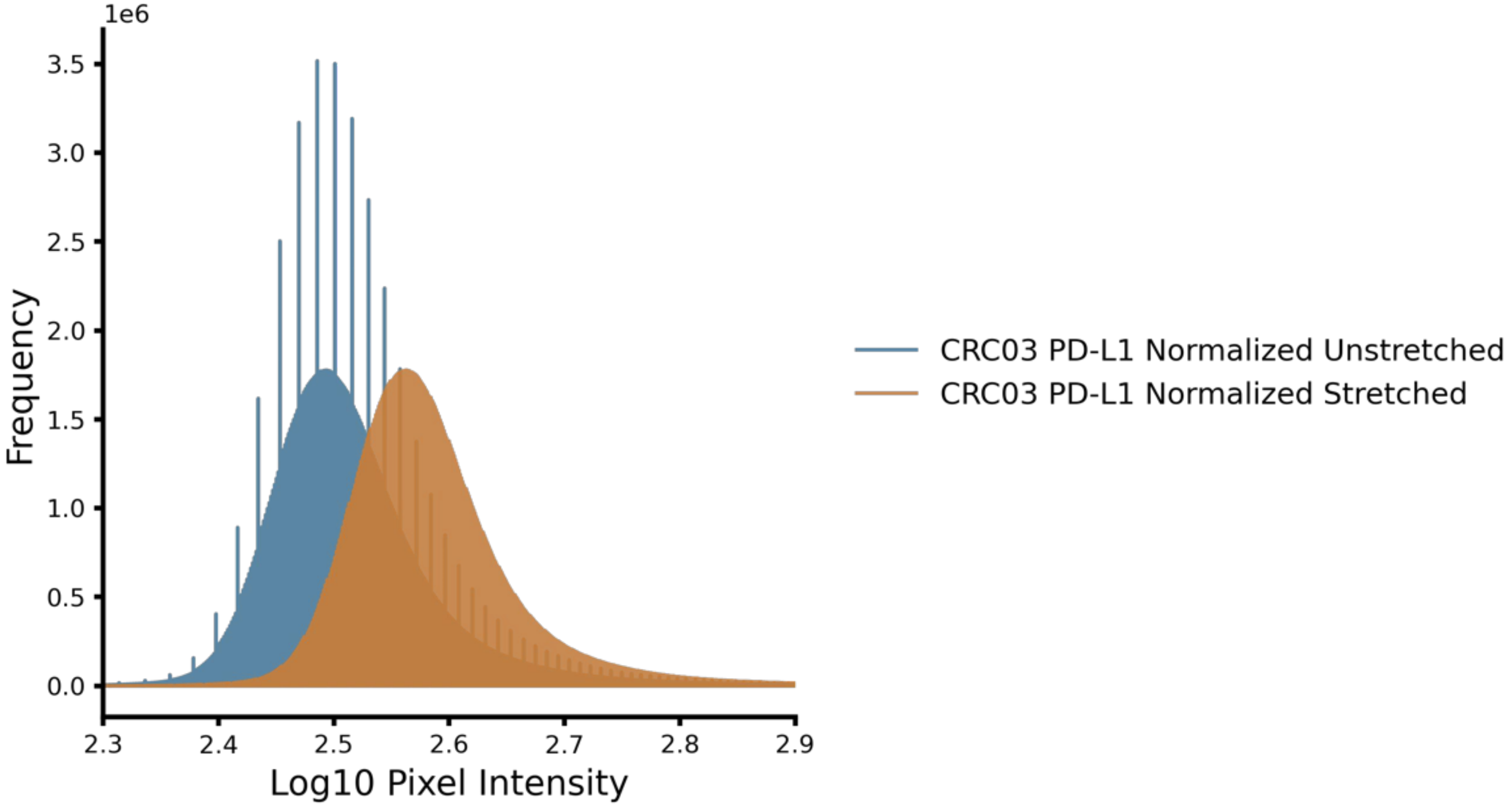


**Figure S10: Effect of Stretching Mechanism in UniFORM Pixel-Level Normalization.** Distribution plots comparing an unstretched (blue) and stretched (orange) marker intensity distribution after pixel-level normalization. When a normalization factor less than 1 is applied and values are converted back to uint16, the reduced dynamic range and limited integer precision introduce quantization artifacts, visible as discrete spikes in the unstretched distribution. The stretched version avoids these artifacts by preserving a broader intensity range, illustrating the importance of managing data type conversion in pixel-level normalization workflows.

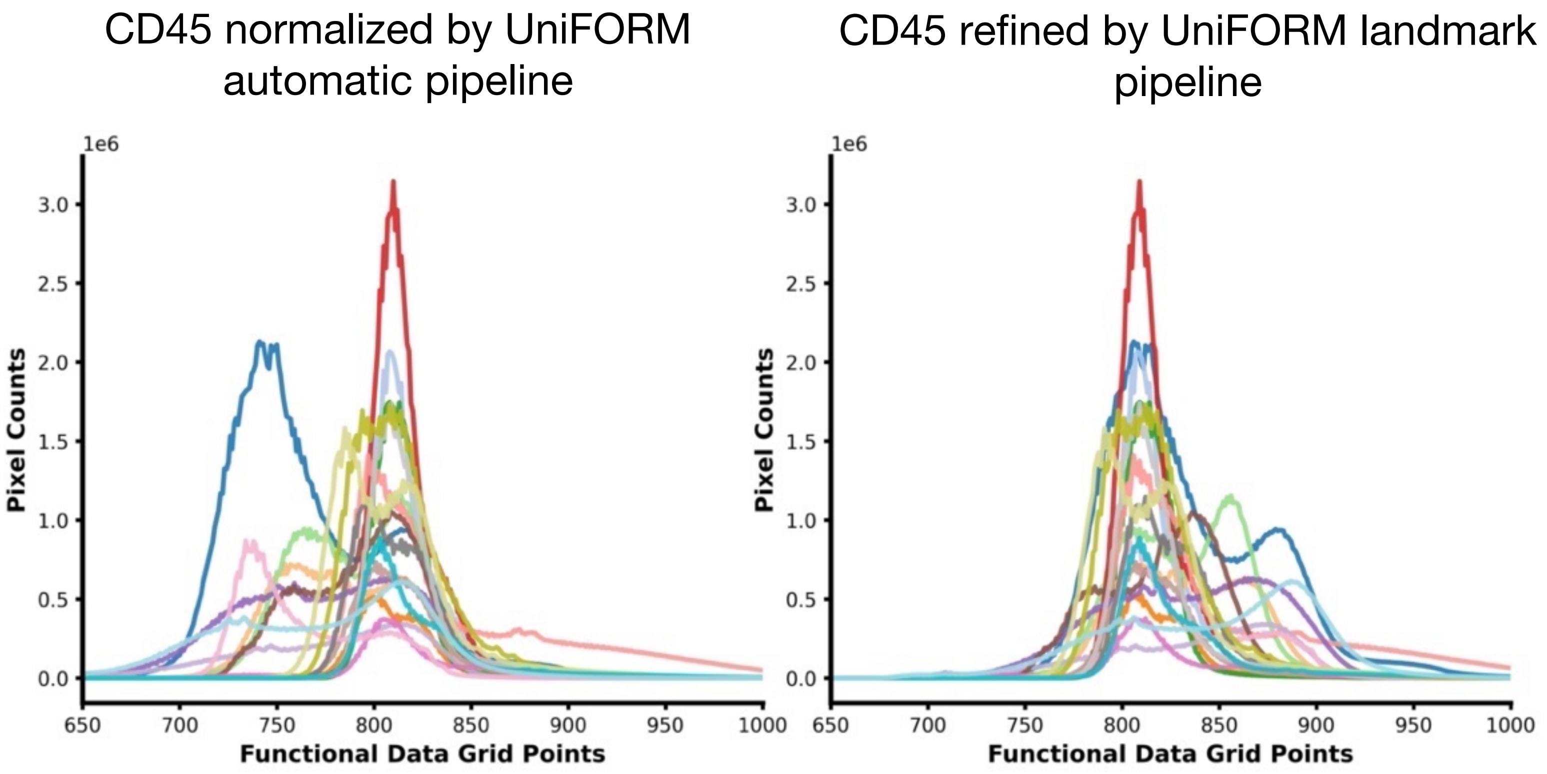


**Figure S11a: Markers that fail the automatic UniFORM normalization pipeline due to severe heterogeneity can be corrected using the guided landmark fine-tuning approach.** The example illustrates how UniFORM's automatic normalization pipeline struggled to align CD45 marker intensities across some samples in the PRAD-CyCIF dataset. By applying the efficient guided landmark fine-tuning option, these discrepancies were resolved, achieving consistent and biologically plausible alignment across samples.

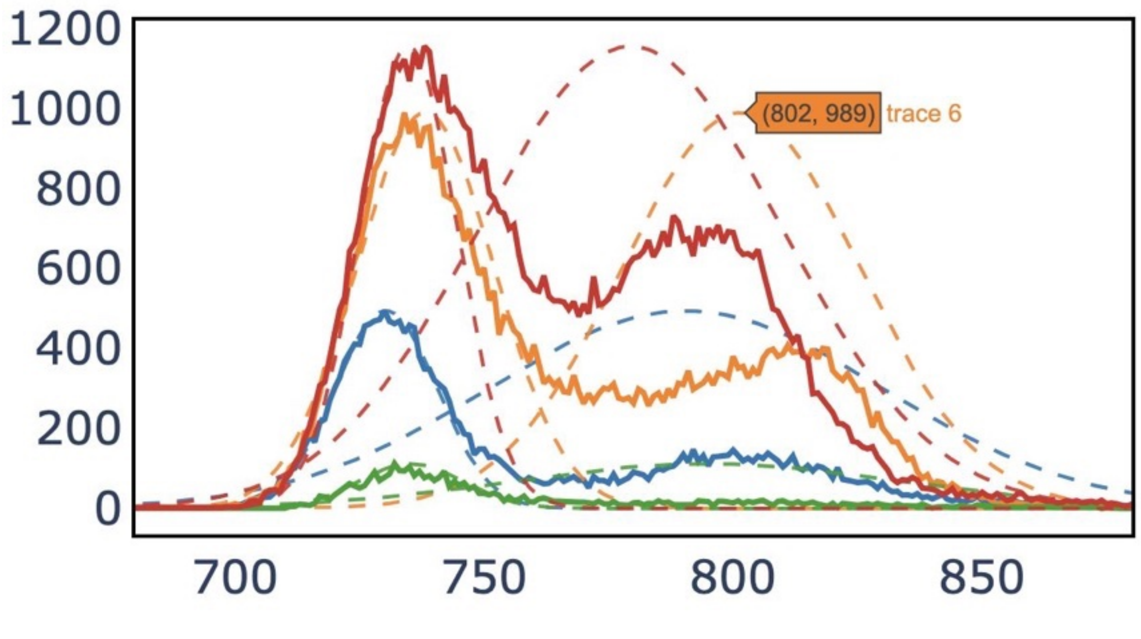


**Figure S11b: The guided landmark normalization pipeline estimates Gaussian curves for the negative and positive populations, aiding users in selecting appropriate landmarks.** The example illustrates the CD45 marker distribution, with Gaussian curves annotated to provide visual guidance for determining landmarks.
